# LemonCatcher Acidic Pull-Down Enables Selective In-Cell Hydrogen-Deuterium Exchange Mass Spectrometry

**DOI:** 10.64898/2026.08.26.747387

**Authors:** Dietmar Hammerschmid, Mahjoobeh Eshani, Anthony H. Keeble, Benjamin Russell Lewis, Valeria Calvaresi, Polina Heatley, Di Zhu, Henry Hayward, Weston B. Struwe, Paula J. Booth, Mark R. Howarth, Eamonn Reading

## Abstract

Proteins are dynamic molecules which sensitively adapt according to their environment. Hydrogen-Deuterium eXchange Mass Spectrometry (HDX-MS) provides unique insights into protein conformational processes. However, existing methodology cannot selectively enrich proteins post-labeling because D-to-H back exchange must be minimized by rapid processing at pH 2.3-3.0 and 0 °C, where affinity purification fails. Here, we create LemonCatcher, a protein superglue that spontaneously forms an amide bond to the LemonTag peptide under these harsh acidic and cold quench conditions, even at -20 °C. Engineering of a bead-coupled LemonCatcher purification system introduces fast and selective quench-capture HDX-MS (SelQueX) on LemonTagged fusion proteins. We demonstrate targeted measurement of protein dynamics in living bacterial cells, revealing ligand-induced conformational changes in maltose-binding protein. Moreover, probing a stalled membrane protein nascent-chain supports a role for the ribosome in maintaining partially unfolded folding intermediates. Thus, SelQueX makes possible selective characterization of protein structural dynamics within the complex cellular milieu.

## Introduction

Linking structure to function through protein dynamics is needed for a full mechanistic understanding of the molecular machines that drive life, health, and disease^1^. Hydrogen-Deuterium eXchange mass spectrometry (HDX-MS) provides unique insights into structural dynamics and is conceptually simple: a protein system is diluted into heavy water (deuterium oxide, D_2_O) and the amount of deuterium integrated into the protein backbone relates to dynamics and accessibility at that site, which is detected by mass spectrometry^2^ (**Fig. 1**). A major strength is that HDX-MS measures solution-phase dynamics, away from the immobilized states (from freezing and/or crystal packing) required for cryo-electron microscopy/tomography and X-ray crystallography^3^. Moreover, it can decipher structural dynamics without suffering from protein size and labelling complexity limitations, which can be the scourge of structural techniques such as NMR^4^. HDX-MS uniquely captures near residue-level transitions between closed conformations (typically in secondary structure) and higher-energy open conformations^5^.

**Figure 1.**
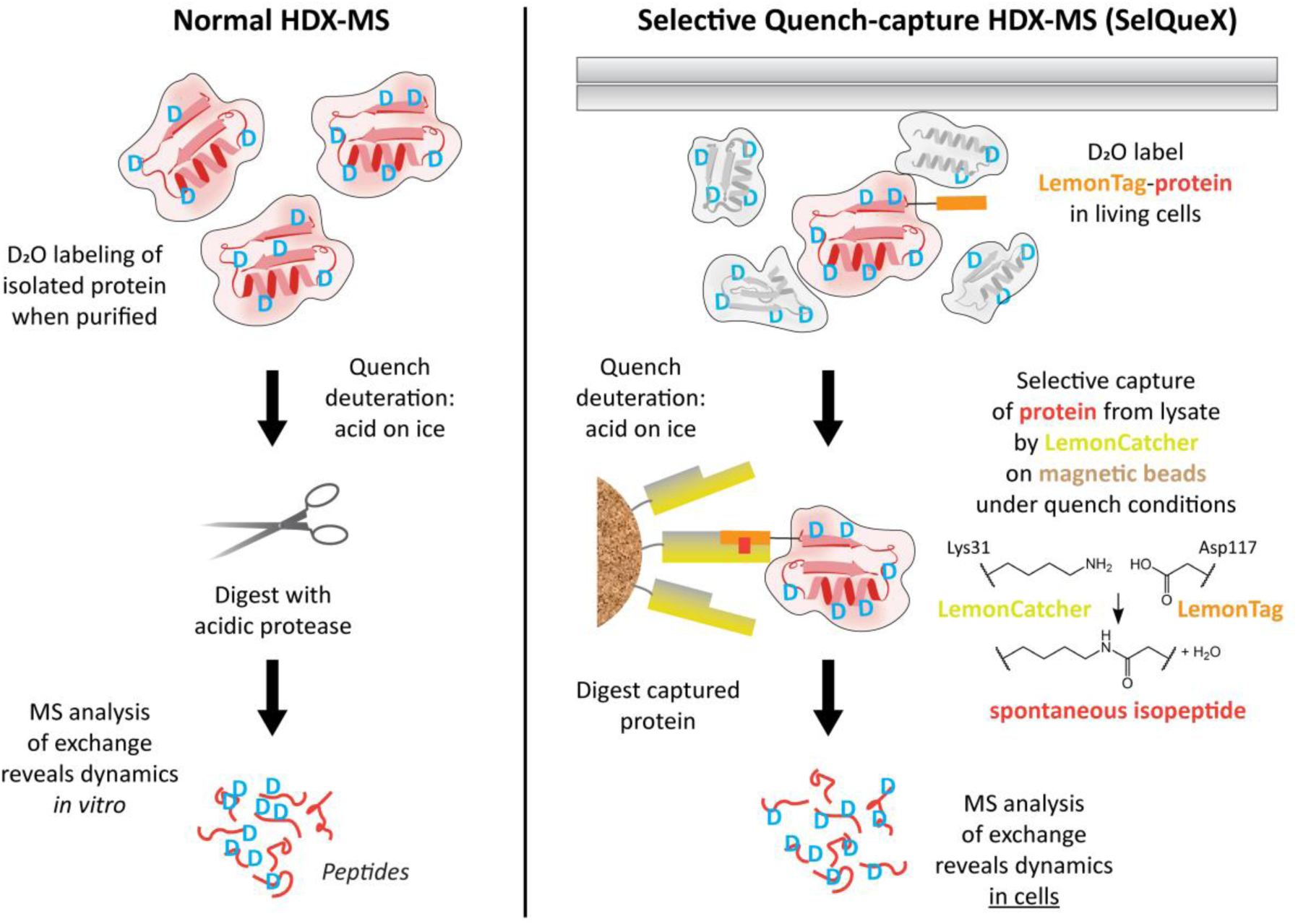
Schematic of Selective Quench-capture HDX-MS (SelQueX). Summary of SelQueX, acidic tag-capture methodology, versus the classical HDX-MS workflow enabled by engineering of SpyTag/SpyCatcher technology to achieve acidophilic “click biology”, through the creation of the LemonTag/LemonCatcher system.

HDX-MS has been demonstrated on proteins within cells at both intact-^6,7^ and peptide-level^8^, but it has never been possible to measure HDX in a selective, protein-targeted manner. Selective protein enrichment post-HDX must take place under the essential acidic and cold quench conditions that preserve the D-label (pH 2.3-3.0 and 0-4 °C)^9^, which disrupt conventional interaction technologies. Immobilized metal-affinity chromatography (IMAC) is only active over a pH range between 6 and 9^10^, barnase/barstar is inactive below pH 4^11,12^, and most antibodies undergo conformational changes at low pH, leading to ligand dissociation^13^. Moreover, deuterium exchange goes both ways, every subsequent step to quenching must be performed quickly to detect D-label incorporation, before D-labeling is all lost to bulk solvent H_2_O. Here, we create an acidophilic protein superglue system which can rapidly purify a protein of interest (POI) following deuteration and quenching, a method we term **Sel**ective **Que**nch-capture HD**X**-MS (SelQueX). SelQueX reverses the classical HDX-MS experiment from “isolate then probe” to “probe then isolate”, ensuring that protein dynamics are captured in their most natural context (**Fig. 1**).

Our strategy was to devise a general pull-down assay using a tag that provides minimal disruption, is independent of the POI, and which could easily be introduced as a fusion tag; akin to poly-histidine or Glutathione S-transferases (GST) tags used routinely for recombinant protein purification. The streptavidin-biotin interaction has been implemented for biotinylated protein depletion under HDX quench conditions but never effectively for purification within a purified *in vitro* system^14^. Additionally, streptavidin pull-down would introduce an additional biotinylation step of an AviTagged POI fusion by biotin ligase, which would require characterization for each construct as this reaction may often not reach completion, even in strains with biotin ligase BirA overexpressed (AVB101, Avidity)^15^. Moreover, biotin-binding partners, streptavidin or avidin (and their derivatives), unlike covalent capture strategies, rely on maintenance of weaker non-covalent interactions and are not good fusion partners^16^. In this work we re-design our protein superglue^17^, where a spontaneous isopeptide bond forms between a specific Asp on SpyTag and Lys on SpyCatcher (**Fig. 1**), to operate efficiently under HDX quench conditions. Harnessing this new pattern of reactivity, we established a robust, broadly applicable approach for enriching target proteins post-HDX. We apply this “click biology”^18^ approach both in living bacterial cells and through selective measurement of a nascent membrane-protein chain emerging from the ribosome, even though the nascent chain constitutes only <2% of the mass of the megadalton complex.

## Results

### Creating acidophilic LemonTag/LemonCatcher peptide-protein superglue

First, we explored whether any existing SpyTag/SpyCatcher pair permutations (with established naming convention of original, 002 or 003)^19^ could react in the acid and cold conditions required to limit D-to-H back exchange. We determined the efficiency of SpyCatcher reaction with maltose-binding protein (MBP) recombinantly tagged with SpyTag (SpyTag-MBP) by SDS-PAGE analysis of the covalent complex (**Fig. 2a**). The SpyTag/SpyCatcher system has a history of improvement for reaction at neutral pH^20,21^ but not within extreme conditions of HDX quench. Intriguingly, the highest, albeit moderate, reconstitution level under HDX quench conditions was observed for SpyCatcher003 in combination with SpyTag002 (**Fig. 2a-b**), which we selected for further optimization. Next, we screened various HDX quench reaction conditions to identify beneficial buffers and additives for the protein/peptide reaction. Low levels (5-15%) of glycerol or sucrose enhanced product formation, although their benefit vanished at higher concentrations (**Fig. S1**). Moreover, the addition of sodium chloride beyond its concentration in PBS buffer (137 mM) decreases the reactivity between both binding partners. These initial conditions with low protein concentration could only provide a system with a reconstitution efficiency of 6 ± 1% (mean ± SD, *n* = 3), even when the reaction progressed longer than the 15-minute reaction time desired (120 min at 1 µM) (**Fig. 2b**). Therefore, further engineering was required to establish a robust solution for acidic tag-capture.

**Figure 2.**
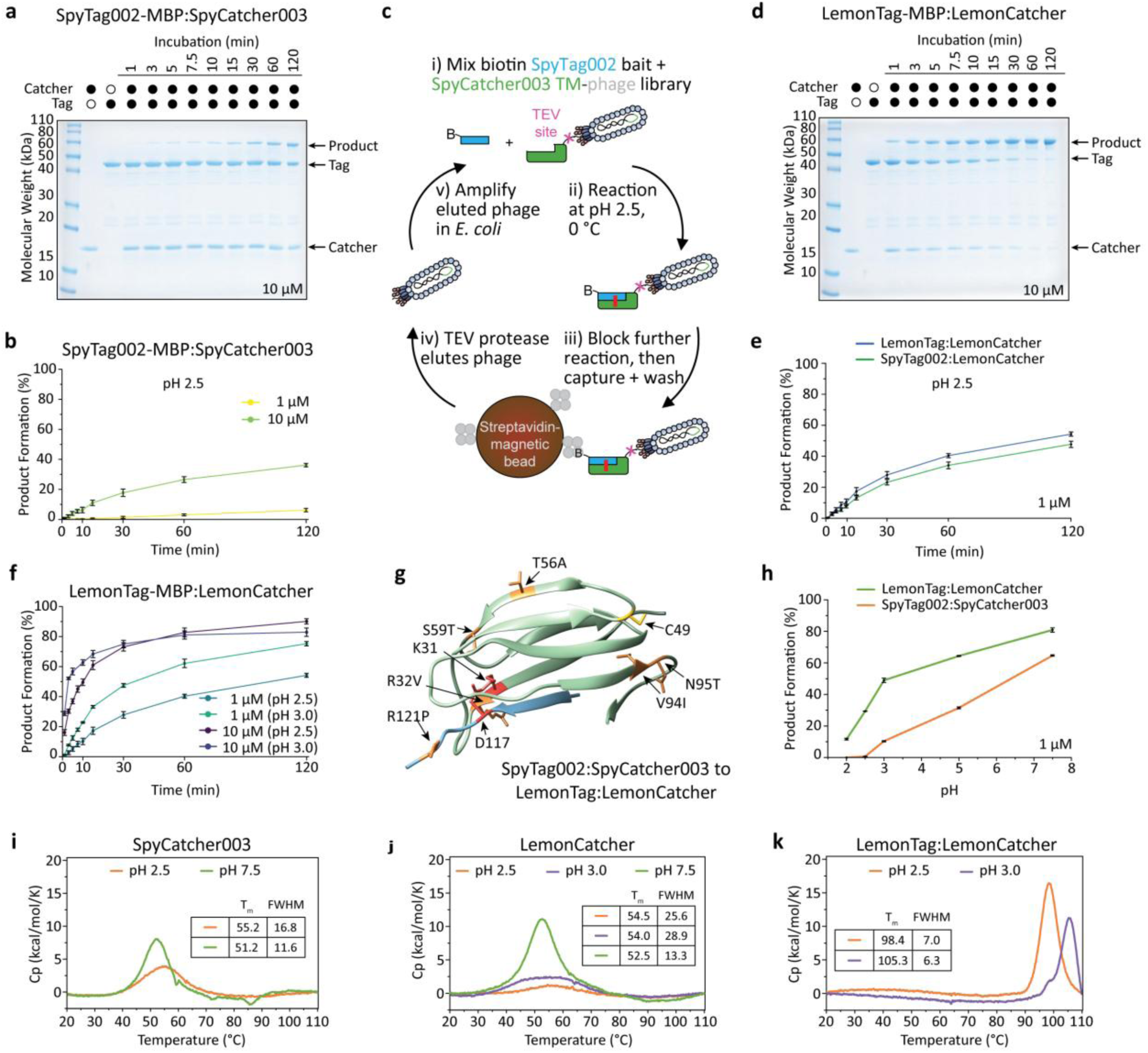
Creation of acidophilic LemonTag/LemonCatcher protein superglue system. (**a**) Representative SDS-PAGE with Coomassie staining for the SpyTag/SpyCatcher reactivity assay. SpyTag002-MBP and SpyCatcher003 were purified by IMAC and size exclusion chromatography. Reconstitution conditions were 10 µM of each protein for the indicated time at 0 °C in quenched PBS sample buffer (pH 2.5) + 2.5 mM TCEP to prevent S49C-mediated disulfide bond formation of Catcher dimers. (**b**) Reaction efficiency comparison of SpyTag002-MBP:SpyCatcher003, pH 2.5 at 1 and 10 µM equimolar concentration. Data are the mean reconstitution product, while error bars indicate 1 s.d. (independent measurements n = 3). (**c**) Schematic of M13 phage displaying on the pIII protein a library based on SpyCatcher003 triple mutant (TM: R32V/S59T/V94I) (not to scale). B represents biotin, gray dots represent streptavidin, and the red line represents an isopeptide bond. (**d**) Representative SDS-PAGE with Coomassie staining for LemonTag/LemonCatcher reactivity assay, prepared as in (a). (**e**) Reaction efficiency comparison of LemonTag/LemonCatcher, pH 2.5 versus SpyTag002/LemonCatcher at 1 µM equimolar concentration. Data are the mean reconstitution product, while error bars indicate 1 s.d. (independent measurements n = 3). (**f**) Reaction efficiency comparison of LemonTag/LemonCatcher, pH 2.5 and 3.0, at 1 and 10 µM equimolar concentration. Data are the mean reconstitution product, while error bars indicate 1 s.d. (independent measurements n = 3). (**g**) Structure of SpyCatcher (green) bound to SpyTag (blue) (PDB: 4MLI^24^) highlighting the mutations made to create LemonTag/LemonCatcher, along with the unique Cys49 residue for surface coupling. Amidation between Lys31 on Catcher with Asp117 on Tag is highlighted in red. (**h**) Reaction efficiency comparison across pH 2.0-7.5. Data are the mean reconstitution product, while error bars indicate 1 s.d. (independent measurements n = 3, with representative gels presented in **Fig. S10**). (**i**) DSC of SpyCatcher003, (**j**) LemonCatcher (**k**) and preformed LemonTag:LemonCatcher complexes within quenched or unquenched PBS sample buffer + 5 mM TCEP at pH 2.5, 3.0 or pH 7.5. Within each graph, the melting temperature (T_m_,) and Full Width Half Maximum (FWHM) in °C is tabulated.

To enhance reactivity, we focused on modifying SpyTag002 and SpyCatcher003 via rational mutations or deletions. At neutral pH, most proteins will have a mixture of positive and negative charges on their surface. At pH 2-3 there may be almost no negative charges remaining, leading to positive-positive electrostatic repulsion reducing protein stability^22^. Initially, we targeted positively-charged amino acids on both binding partners (**Fig. S2** and **Table S1**), since the high degree of protonation of carboxylates from Asp/Glu (typical pKa ∼ 4.4^23^) at such an acidic pH might lead to destabilization of the Catcher and its interaction to the Tag through electrostatic repulsion between the positively-charged sidechains. Mutating Arg32 next to the reactive Lys31 in SpyCatcher003 led to enhanced reactivity with SpyTag002 (**Fig. S2a**). Our first attempt was R32T substitution, since T is well tolerated in β-strands, and this mutation increased reactivity to ∼20-25% within 120 minutes. However, the reactivity was improved further by 52 ± 4% (mean change ± SD, *n* = 3), compared to R32T, by introduction of a hydrophobic residue here (R32V) (**Fig. S2c**). Although this mutation was successful, targeting other amino acids likely to be charged at pH 2-3 within SpyCatcher003 (H26N, K28I, R37N, R47T, K52E, H62Q, K72E, and H112Q) and SpyTag002 (K120E, R121T, K120E-R121T) did not improve reactivity (**Fig. S2b-d**). In addition, deletion of the positively charged KRYK C-terminus of SpyTag002 led to a striking reduction in amidation efficiency (**Fig. S2e**). These data support that a strategy beyond rational deletion or substitution of positively-charged residues would be required for substantial acid-adaptation of reactivity.

Therefore, we established a directed evolution approach, harnessing the exceptional tolerance of bacteriophage to such extreme pH^25^. We generated a phage library based on SpyCatcher003 and panned for covalent bond formation under acidic and cold conditions (**Fig. 2c** and **S3**). From this library we discovered two extra mutations which improved reactivity, S59T and V94I (by 39 ± 3% and 21 ± 6% respectively (mean change ± SD, *n* = 3); **Fig. S4**). Combining these mutants into a triple mutant (TM) R32V/S59T/V94I construct majorly improved the reactivity (**Fig. S5**). Using SpyCatcher003 R32V/S59T/V94I as a template, we then performed phage display under multiple rounds of selection in HDX quench conditions, which provided another two beneficial mutations (T56A and N95T). When combined with the original construct, this set of substitutions (R32V/T56A/S59T/V94I/N95T) conferred the highest reactivity and was designated LemonCatcher (**Fig. 2d**, **S5f** and **S6**). We then revisited the optimal Tag for our pairing with LemonCatcher; R121P mutation of SpyTag002 (Pro121 being from the original SpyTag^17^) markedly improved reactivity (20 ± 8%; mean change ± SD, *n* = 3) (**Fig. 2e** and **S5f**). Our final selection of SpyTag002 R121P was termed LemonTag. In combination, we had achieved a protein superglue pair highly adapted to HDX quench conditions which we named LemonTag/LemonCatcher (**Fig. 2d-g**), since this pair reacted well in lemon juice which has a matching pH of 2-3^26^ (**Fig. S7**, with amino acid sequences in **Fig. S8**).

### LemonTag/LemonCatcher reactivity and stability

We succeeded in advancing SpyTag/SpyCatcher from negligible reactivity under HDX quench conditions to achieving >60% rapid covalent capture (in <5 minutes) of a recombinantly Lemon-tagged protein (**Fig. 2b** *versus* **2f**). We validated spontaneous covalent isopeptide formation between LemonCatcher and LemonTag under these conditions by intact protein electrospray-ionization mass spectrometry (**Fig. S9**). Further assessment of LemonTag/LemonCatcher shows reactivity including both acidic and more neutral pH settings; interestingly reaction is even faster at pH 7.5 than the SpyCatcher003/SpyTag002 benchmark (**Fig. 2h** and **S10**). LemonTag/LemonCatcher reactivity improved from pH 2.0 to 3.0 (**Fig. 2f** and **2h**), the ideal pH range to quench HDX reactions^9^.

Differential scanning calorimetry (DSC) revealed that the melting temperatures for apo LemonCatcher compared to SpyCatcher003 are similar at both pH 2.5 and 7.5 (*T*_m_ values of 51.2-56.1 °C) (**Fig. 2i-j** and **S11**). Nonetheless, the full width at half maximum (FWHM) values are wider for LemonCatcher within acid conditions, which suggests comparatively less cooperative protein unfolding at low pH. As seen for SpyCatcher003^20^, reaction of LemonCatcher with its Tag drastically improved thermal stability: up to 98.4 °C (LemonTag:LemonCatcher) and 98.9 °C (SpyTag002:LemonCatcher) at pH 2.5, and 105.3 °C (LemonTag:LemonCatcher) at pH 3.0 (**Fig. 2k** and **S11**). Such stabilization upon Tag reaction is thought to be important thermodynamically in driving isopeptide bond formation towards completion^17,20^. The LemonTag/LemonCatcher reaction tolerated different quench components compatible with HDX-MS, being largely unaltered in various acidified buffers (i.e. glycine, phosphate), denaturant (up to 4M Urea) and mild detergents, e.g. n-dodecyl-β-D-maltoside (DDM) but demonstrated significantly reduced efficiency in harsher n-octyl-β-D-glucoside (OG) and abolished reactivity for Fos-Choline 12 (**Fig. S12**). Strikingly, we found that the protein superglue can even retain reactivity despite acidic, subzero temperature conditions of -20 °C, enabled by the presence of an antifreeze buffer modifier^27^ (ethylene glycol, 37% w/w) (**Fig. S13a-c**). Although the reaction is slowed under these conditions, coupled reaction and back exchange times align with the SelQueX protocol when performed on ice (**Fig. S13d**). Taken together, these results demonstrate that the LemonTag/LemonCatcher reaction system is compatible with a wide range of quench-and-capture conditions, granting versatility in workflow design and allowing fine-tuning for needs of particular systems.

### Establishing selective quench tag-capture HDX-MS (SelQueX)

Creation of a selective quench tag-capture method requires the LemonCatcher to be linked to a suitable pull-down device, so that the LemonTagged POI can be isolated from the complex sample background. Doing so will decrease non-targeted peptide co-elution and potential protease and column fouling, which hampers protein identification and analysis^28^. Our LemonCatcher construct contains a unique cysteine on the opposite side of the protein to the reactive center (C49; **Fig. 2g**) and thus offers surface coupling capability via disulfide bridge formation. Different bead systems were trialed using a non-reactive LemonTag(PA)-MBP construct (containing M115P D117A mutation) as a control during workflow development, to act as a reporter for non-selective enrichment and false-positives (**Table S2** and see **Methods**); D117A abolishes spontaneous isopeptide bond formation with Lys31 in the Catcher, whereas M115P is deleterious to non-covalent binding with Catcher (**Fig. S14**).

Following careful optimization, we established that a POI, recombinantly tagged with LemonTag, could be selectively pulled down by streptavidin-magnetic beads coupled with biotin-modified LemonCatcher for both purified and cell lysate conditions (**Fig. 3a** and see **Methods**). Biotin-HPDP (HPDP = N-[6-(biotinamido)hexyl]-3’-(2-pyridyldithio)propionamide) was used to react with the free sulfhydryl (-SH) groups on the unique cysteine within LemonCatcher (C49) to form a cleavable disulfide bond, allowing reversible biotinylation useful in elution after affinity purification (**Fig. S15a**). Biotinylation did not affect LemonTag/LemonCatcher reactivity (**Fig. S15b**). This strategy was inspired by the work of the Guttman laboratory who used a similar cleavable biotin-streptavidin system to enrich antibody complexes within pre-quench (i.e. non-acidic and time-limiting) HDX-MS conditions^29^. Our workflow required a pH of 3.0 to reliably capture LemonTag-MBP using these LemonCatcher beads, which may result from excessive electrostatic repulsion of the protein backbones at pH <2.5^28^. Importantly, pH 3.0 caused negligible increase in back exchange compared to pH 2.5, as evaluated using peptide standards (**Fig. S13d**).

**Figure 3.**
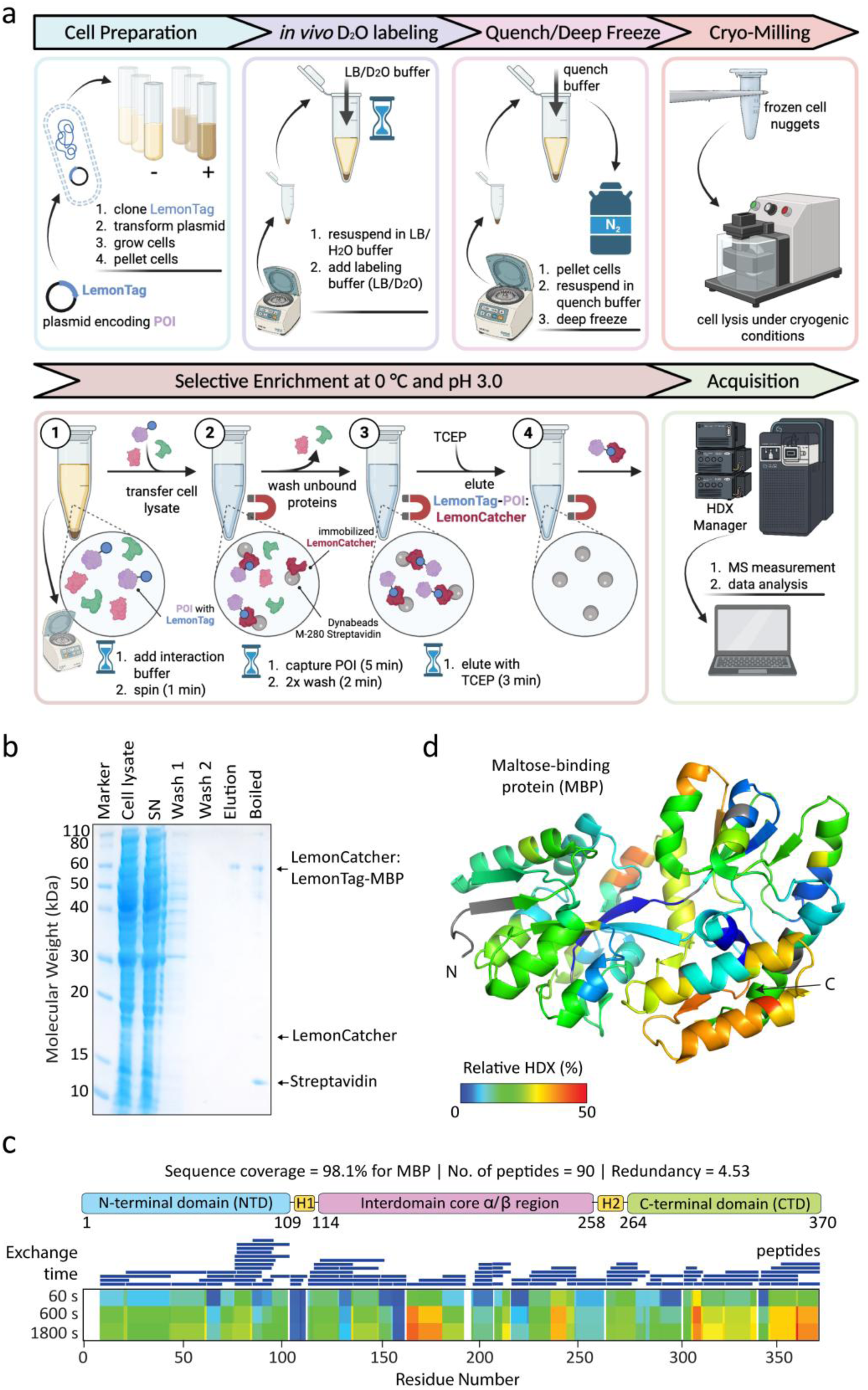
Cellular HDX-MS using SelQueX. (**a**) SelQueX workflow (created with BioRender.com). (**b**) Representative SDS-PAGE with Coomassie staining for cellular SelQueX on POI (LemonTag-MBP) captured from E. coli. SN is the supernatant after centrifugation of cell lysate. Boiled samples lanes denote the remaining bound protein species on the resin after elution, removed after boiling in SDS-containing loading buffer for 5 minutes (see **Methods**). Reproducibility of the workflow is demonstrated in replicate analysis in **Fig. S16b**. (**c**) Peptide coverage map -*after filtering based on deuterated spectral quality - achieved for SelQueX on LemonTag-MBP. (**d**) Relative fractional uptake of deuterium mapped onto the structure of apo-MBP (PDB: 1PEB*^35^*). Relative HDX color bar key depicts range for both plot in (c) and on protein structure in (d)*.

Whilst the bead-captured POI could potentially be digested while anchored to beads, we devised a strategy for elution of the entire complex (LemonTag-POI:LemonCatcher) with tris(2-carboxyethyl)phosphine (TCEP), which is tolerated at up to 0.5 M by pepsin^30^. Release of the purified complex permits digestion by immobilized proteases (such as pepsin, nepenthesin and fungal protease type XIII^31–33)^, which improves digestion efficiency, reproducibility, throughput, and decreases background from digestion of scaffold streptavidin as well as autolytic products prevalent with solution digests^34^. For elution, we found that a condition of 0.5 M TCEP/4 M urea/pH 3.0 (buffered with formic acid) was potent enough to elute >50% of LemonTag-POI:LemonCatcher complex in 3 minutes. These results show highly effective purification of the POI away from an *E. coli* lysate in 10 minutes (>90% from SDS-PAGE; time combines all bead binding, washing, and elution steps) (**Fig. 3b** and **S16**). The back exchange increase was only ∼16% (from 32 ± 2% for *in vitro* HDX-MS to 48 ± 10% for cellular quench tag-capture workflow) (**Table S3**). SelQueX achieves high level peptide identification coverage: 79-98%, depending on the sensitivity of the MS instrumentation (**Fig. 3c** and **S17**). Importantly, performing the same procedure with the non-reactive LemonTag(PA)-MBP control yielded no assignable peptides, consistent with the absence of a corresponding covalent LemonCatcher complex band in SDS-PAGE analysis on both purified and cell lysate conditions (**Fig. S16a**). These data validate that the SelQueX method enables truly selective capture from cells under quench conditions.

### Measuring structural dynamics on a cellular-confined protein

We next proved SelQueX applicability to achieve in-cell structural dynamics insight using LemonTag-MBP expressed in the *E. coli* cytosol. MBP’s natural function is in the transport and chemotaxis of maltodextrins, including maltose, but is also used in molecular biology as an effective solubility tag for proteins in the bacterial cytosol^36^. Cells with overexpressed LemonTag-MBP were incubated in Luria Broth/D_2_O buffer for 1, 10 or 30 minutes. Labeling proteins *in vivo* by HDX requires an additional appropriate lysis strategy that prevents extensive back exchange, therefore cell lysis under cryogenic milling conditions was performed^8^ (**Fig. 3a** and see **Methods**). Cellular deuterium uptake obtained for cytosolic LemonTag-MBP within living cells revealed areas with time-dependent exchange. Unstructured regions exchange rapidly, while well-ordered hydrogen-bonded regions exchange more slowly, characteristic of a folded protein with differences in secondary structure and dynamics (**Fig. 3c-d**). The protein core surrounding the maltose-binding pocket showed the least exchange, whereas the “spring-loaded” hinge or balancing interface (residues 175-184), C-terminal helix (residues 350-370) and loop regions spanning residues 304-314 and 238-245 had the highest exchange.

We next compared LemonTag-MBP HDX profile changes between maltose-fed and unfed cells (i.e. differential HDX analysis, ΔHDX) on independent biological replicates (*n* = 3, direct HDX-MS assessment of different bacterial colony isolates). Maltose was added during cell growth to stimulate maltose import into the cytoplasm by the *E. coli* maltose transport system^37^ (**Fig. 4a**), with maltose levels (1 mM) maintained in the buffers during cellular deuteration. Significant changes in backbone HDX (p ≤ 0.05; ΔHDX significance cut-off of 0.40 Da) were caused by maltose ligand introduction within multiple peptides encompassing residue sections 8-61, 77-103, 163-192, and 340-370, with some isolated peptides demonstrating significance in other areas of the protein (**Fig. 4b-e**). Many of the areas and directionality of dynamical perturbation on cellular-contained LemonTag-MBP corroborate previous *in vitro* maltose-bound MBP dynamics determined by classical HDX-MS^38^, with decreased HDX observed in regions close by the maltose ligand binding pocket where packing is altered as the protein closes^39^. However, distinctly pronounced decreased HDX was observed within the balancing interface and C-terminal helix in our cellular measurements (**Fig. 4d**). There were also allosteric sites in the N-terminal domain (peptides including residues 77-103 and 263-282) which became less protected to exchange within cytoplasmic MBP in the presence of maltose, suggestive of ligand-evoked increase in backbone flexibility upon closure. Control experiments further supported cellular ligand engagement. No significant HDX changes were observed upon maltose addition during cellular deuteration to maltose-unfed cells, where the maltose import machinery is not induced. In contrast, maltose-fed cells displayed weaker but similar perturbations in the absence of extracellular maltose during cellular deuteration, consistent with intracellular maltose depletion over time (**Fig. S18**). Overall, this is consistent with backbone perturbations putative of a closed, maltose-bound form of MBP, being observed in living bacterial cells by the SelQueX method.

**Figure 4.**
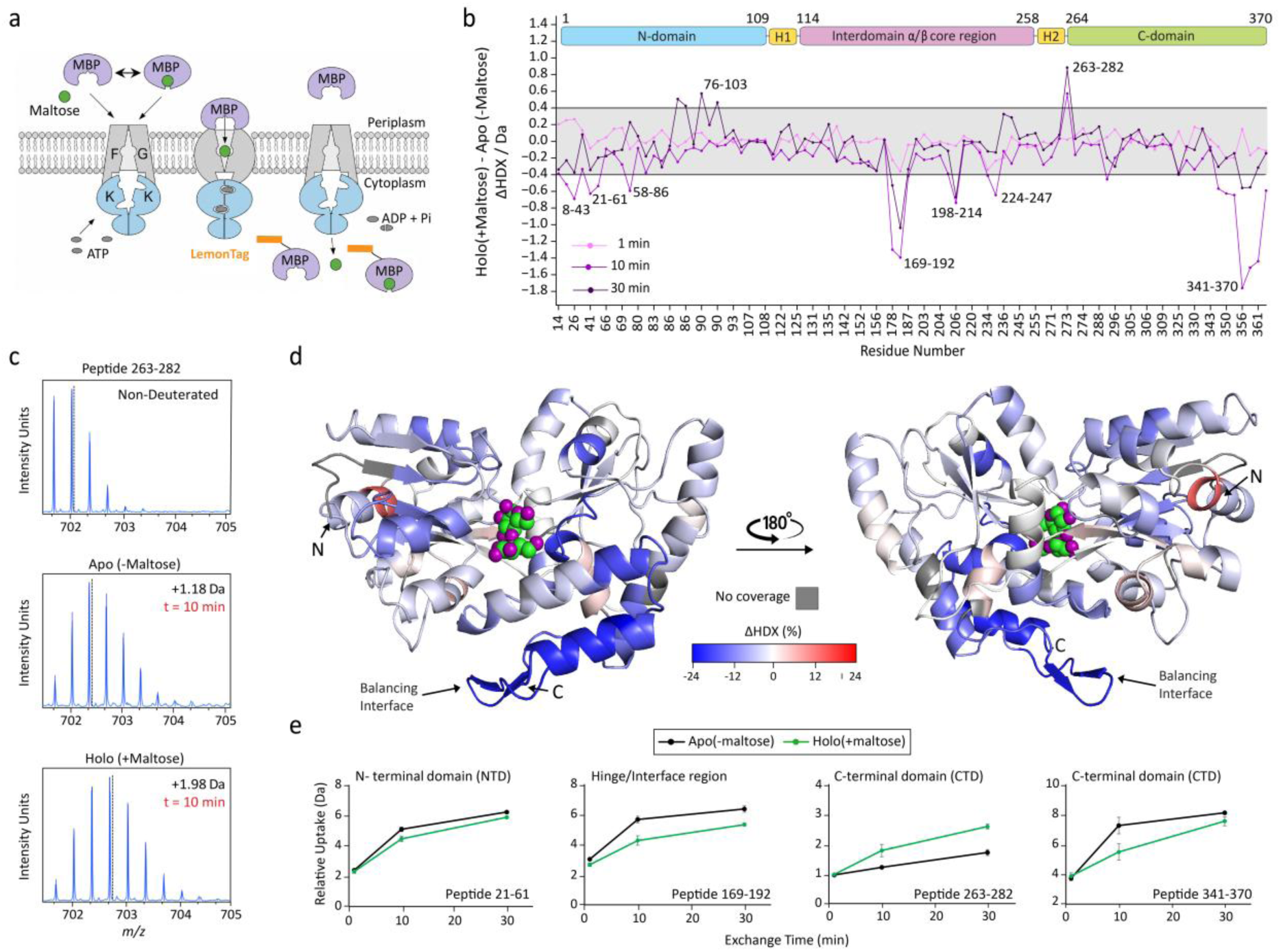
SelQueX characterization of dynamic shifts in cytoplasmic-contained MBP driven by a naturally imported cellular ligand. *(a) Schematic of the experiment; maltose is introduced to the bacterial population which triggers MalFGK_2_ maltose/maltodextrin ABC transporter complex expression to enable maltose uptake into the cytoplasm, where LemonTag-MBP has been expressed. (b) Differential plot from SelQueX illustrates the difference in HDX (ΔHDX) between maltose-fed cells and maltose-absent cell populations across the time points studied. The dashed lines indicate the threshold of significance for ΔHDX (p ≤ 0.05; ΔHDX significance cut-off of 0.40 Da; n_biological_ = 3). The MBP domains are illustrated in the top bars and residues comprising a region with significant differences in HDX are indicated. Peptides are arranged from the N- to C-terminus according to their peptide center residue. (c) Representative m/z spectrum for peptide 263-282 under non-deuterating conditions and deuterating conditions. The centroid is represented by the dotted line, and the mass change of the deuterated samples is written in Daltons. (d) Regions manifesting decreased HDX in holo-MBP versus apo-MBP are superimposed and colored blue, while increased HDX is colored red. Based on the liganded closed structure (PDB: 3MBP*^40^*); ligand shown in space fill; regions with no coverage are colored gray. %ΔHDX was corrected using a maximally deuterated control (MaxD). (e) Representative uptake plots for four peptides in different domains of MBP. Uptake plots are the average deuterium uptake, and error bars indicate ± 1 standard deviation (n_biological_ = 3)*.

### Targeted membrane protein nascent chain detection within the megadalton ribosome

SelQueX should possess the ability to extract structure-dynamics information of Lemontagged POIs, whether that is embedded in a cellular proteome or in a large bio-macromolecular complex. To evaluate this ability, we challenged SelQueX to selectively capture HDX information on a membrane protein nascent chain emerging from the megadalton ribosome into detergent micelles. HDX-MS has recently emerged as a key tool in interpreting co-translational nascent chain folding intermediates that are trapped within ribosome nascent chain (RNC) complexes^41,42^. On the other hand, HDX-MS insight into membrane protein RNC complexes has yet to be achieved. Existing protocols to explicitly determine nascent chain information are hampered by spectral crowding from auxiliary ribosomal protein components, as well as prospective proteolysis, chromatography fouling, and signal suppression by non-proteinaceous species such as RNA^43,44^. GlpG is an intramembrane serine protease, essential for colonization by extraintestinal pathogenic *Escherichia coli*^45^. We applied our previous work on GlpG RNC complexes^46^ and recombinantly inserted an N-terminal LemonTag to a GlpG co-translational folding intermediate; which has the N-terminal cytosolic domain (CytD) and subsequent four transmembrane helices (TM) translated by the ribosomes, tethered by a stall sequence containing the *E. coli* SecM arrest peptide (construct denoted as GlpG(4TM); **Fig. 5a-c** and **S19a**). By adapting SelQueX to our RNC construct, we found excellent nascent chain capture, achieving 28 high-quality peptides covering 94% of the emerging protein chain (**Fig. 5b-c** and **S19b**), determining HDX across the nascent chain. It is worth noting that our methodology could also be used to fix the RNC, or any POI, to a solid support before or during deuteration (as LemonCatcher/LemonTag is highly reactive across pH 2.5-7.5, **Fig. 2h**) but this procedural arrangement increases the chance of non-native conformational stasis.

**Figure 5.**
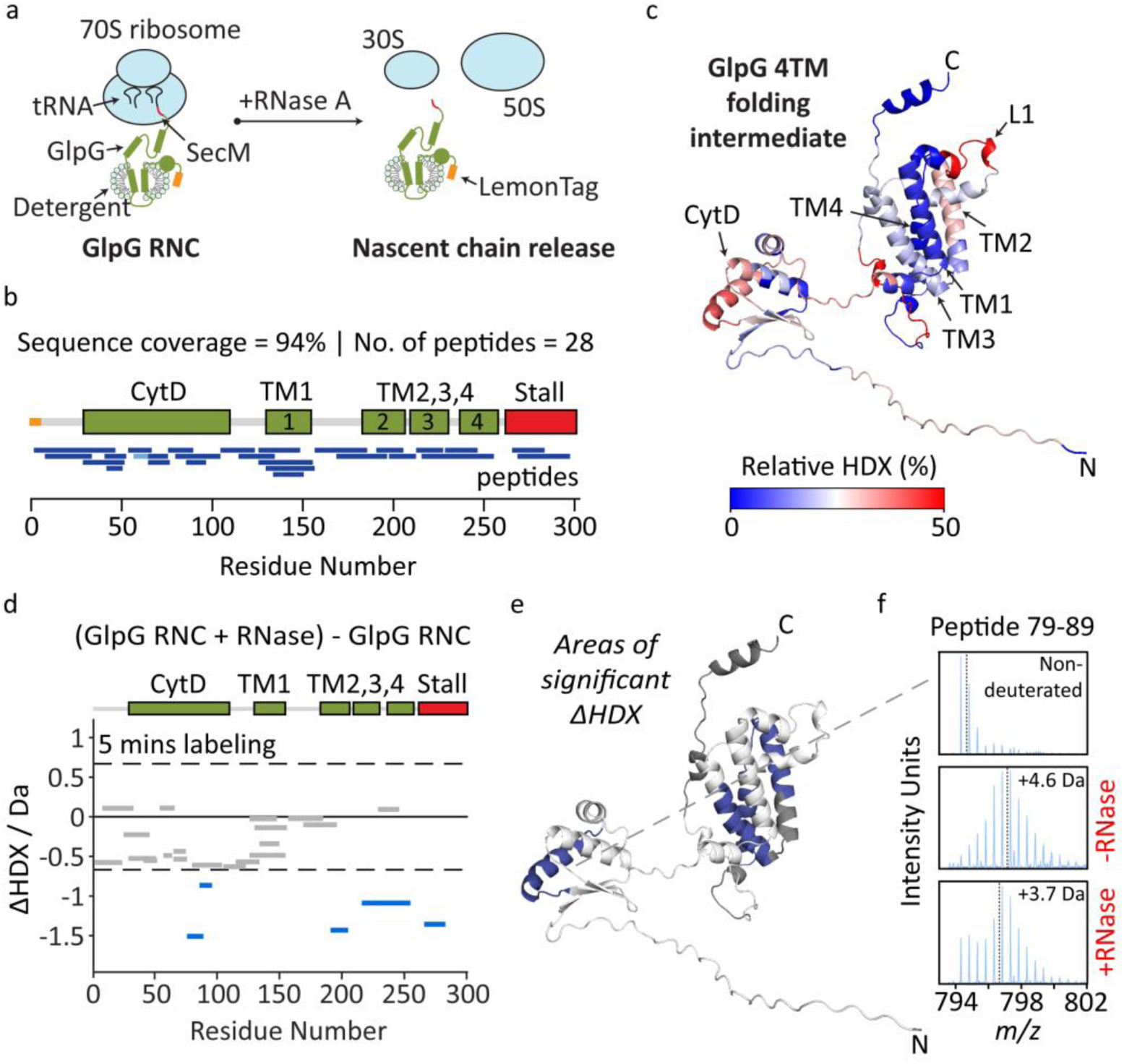
SelQueX on an emerging membrane protein nascent chain within a ribosome nascent chain complex (RNC). (**a**) Adapted SelQueX workflow applied to the Lemontagged GlpG 4TMH RNC construct. (**b**) Peptide coverage map after visual inspection of the deuterated peptides for GlpG 4TMH RNC. (**c**) Relative fraction uptake of deuterium onto the AlphaFold3^49^ model of GlpG 4TMH. (**d**) Woods plot displaying ΔHDX for GlpG 4TMH RNC +/- RNase for the 5-minute pulsed timepoint. Blue signifies a decrease in HDX between states. Significance was defined as ΔHDX = p ≤ 0.05; ΔHDX significance cut-off of 0.67 Da; n_biological_ = 3, and is represented by the dashed lines on the plot. (**e**) Areas of significant ΔHDX onto the structure of GlpG 4TMH. Regions in gray represent no coverage. (**f**) A representative spectrum for peptide 79-89 under non-deuterated and deuterated states is shown. Centroid is shown by a dotted line and the mass change shown in Daltons.

Building upon this, we next compared pulse HDX profiles (5 minutes deuteration time) of the folding intermediate in both ribosomes tethered and released forms (**Fig. 5d-e**). Prior to labelling, the RNC sample was incubated with either standard protein buffer or RNase, the latter resulting in digestion of the ribosomal transfer RNA (tRNA) and subsequent release of the nascent chain (**Fig. 5a** and **S19a**). In the presence of RNase, we observed that TM2-4 and regions of CytD exhibited a decrease in HDX compared to the nascent chain embedded in the ribosome. This change in exchange may suggest that the ribosome is involved in interactions with the nascent chain during co-translational folding, influencing GlpG’s energy landscape to exist in more unfolded states, potentially preventing misfolding or aggregation as observed within soluble nascent chain folding intermediates^47,48^. Indeed, mixed-modal HDX spectra in regions of the CytD domain indicates possible aggregate populations following ribosomal release by RNase treatment (**Fig. 5f** and **S20**). Such multi-modal isotopic distributions can hallmark of heterogeneous, non-interconverting subpopulations during aggregation, where some molecules become trapped in exchange-protected aggregate states^2^. Overall, we show that SelQueX can be applied to large bio-macromolecular complexes to unlock previously unattainable samples to be interrogated by HDX-MS, enabling future work for the determination of an entire folding pathway of a membrane protein chain as it emerges from the ribosome.

## Discussion

Recent advances in AI-driven structure prediction and design, including AlphaFold^49^ and RFdiffusion^50^, have transformed structural biology, yet remain largely rooted in static structural representations. Consequently, we often understand very little about structural transitions and dynamics of proteins, where characterizing structural dynamics experimentally within cellular systems offers the key to understanding how molecules carry out their biological functions. More broadly, there is increasing recognition that structural biology must move beyond isolated biomolecules towards empirical measurements of protein structure-function relationships within cellular environments^1^. Techniques such as cryo-electron tomography^51^ and in-cell NMR^52^ have pioneered this transition, providing valuable insights into molecular organisation and behaviour *in situ*. Nevertheless, cryo-ET primarily captures ensemble snapshots of frozen states and remains challenging for resolving dynamic conformational transitions at high temporal resolution, while in-cell NMR often requires specialised isotopic labelling and is restricted by molecular size and spectral complexity. Further refined experimental methods capable of measuring protein dynamics in their native context will therefore be needed for developing the next generation of dynamics-aware molecular models^1^.

Here, we have demonstrated that SelQueX, enabled by creation of a simple and highly stable acidophilic protein–peptide “superglue”, provides selective and readily implemented access to protein structural dynamics amongst biological complexity. Using this approach, we resolved ligand-induced changes in maltose-binding protein structural dynamics directly in living bacterial cells, establishing targeted HDX-MS measurements across multiple levels of biological complexity, including highly heterogeneous purified systems such as membrane protein ribosome nascent chain complexes (RNCs). To our knowledge, this represents the first validated HDX-MS detection of ligand engagement for a protein within the interior of living cells, as well as on a membrane protein RNC. Whereas previous in-cell HDX-MS studies monitored protein aggregation/unfolding^6,7^ or peptide binding to an outer membrane protein^8^, we directly resolved maltose ligand-induced changes on cytoplasmic expressed MBP, including dynamical stabilisation of the binding pocket, and allosteric regulation of the balancing interface and C-terminal helix that govern the equilibrium between open and closed conformations^39,53–56^. Moreover, selective probing of a nascent membrane protein provides evidence consistent with ribosome-mediated stabilisation of more dynamic, partially folded, cytosolic and transmembrane structural elements, as established for nascent multidomain soluble proteins^47,48^. Given the central role of HDX-MS in modern structural biology and drug discovery^57–59^, extending such measurements to living cells enables direct assessment of target engagement and its conformational consequences within the native cellular environment.

One of the criteria for “click biology” is reactivity under a broad range of conditions; there has been extensive protein engineering to achieve activity in thermophilic zones but much less exploration of protein functionality in these acidophilic and psychrophilic zones^18,60^. Peak reactivity of the different SpyTag/SpyCatcher pairs is close to neutral pH^17,20,61^. One would anticipate minimal reactivity of SpyTag/SpyCatcher at pH 2-3: the mechanism of spontaneous isopeptide bond formation depends on Lys31 being deprotonated^62^ (-NH_3_^+^ is non-nucleophilic) and the normal pKa of lysine is ∼10.0^23^. It is established that burial of polar sidechains can modify pKa values by a few units^63,64^, but the extent is surprising in enabling efficient LemonTag/LemonCatcher amidation at pH 2.5. Going from SpyTag003 to LemonTag removes three residues that would be positively charged at pH 2.5, so reducing electrostatic repulsion to LemonCatcher. S59T and V94I were previously identified from phage selection of SpySwitch^65^, so their benefit may be general in enhancing Catcher stability and not restricted to low pH. R32V may generate favorable non-polar contacts in docking with V114 and V116 on LemonTag. To gain further insights into LemonTag/LemonCatcher reactivity AlphaFold 3^49^ was used to model the complex and compare to SpyTag/SpyCatcher (**Fig. S21**). Future biophysical investigations may probe the contributions of the mutations to spontaneous isopeptide formation under such challenge quench conditions.

A disadvantage is that the benchmarked workflow presented here requires LemonCatcher elution with the POI. Although LemonCatcher did not considerably interfere with POI peptide identification and HDX analysis, which we attribute to the small size of the Catcher (<120 residues, ∼12.5 kDa), development of an elution strategy which avoids LemonCatcher presence could be beneficial. Strikingly, we discover that the protein superglue even works within acidic -20 °C conditions with antifreeze^66^, at which temperature D-label loss is almost eliminated^27^ (**Fig. S13**). Although LemonTag/LemonCatcher reaction is slowed here, it offers a starting position to increase reaction kinetics to achieve a complete subzero, back exchange-absent method. We predict that the simplicity of a protein fusion Tag/Catcher system (LemonTag is only 14 amino acids long) affords sufficient flexibility for multiple future selective quench-capture workflows, e.g. utilizing on-bead digestion, LemonTag-decorated beads, or fluidic HDX-MS devices^67,68^. Ultimately, SelQueX will be most powerful when combined with recent breakthroughs in nano-scaled HDX-MS^69^, decoupled platforms^70^, automated HDX analysis^71,72^ and subzero chromatography technologies^27^ which, in turn, increase sensitivity, enhance throughput, and will rectify additional back exchange experienced during work-up by eradicating back exchange during LC-MS. It is exciting to anticipate how integration of SelQueX with these emerging advances could aid in higher throughput HDX-MS structural biology^73,74^ on protein dynamics without disrupting biological complexity and pave the way to studying single cell and endogenous levels of proteins within their physiological environments.

## Acknowledgements

E.R, D.H., and M.E, were supported by a UK Research and Innovation (UKRI) Future Leaders Fellowship (MR/S015426/1 and MR/X009580/1) to E.R.. A.H.K. and M.R.H. were funded by the Biotechnology and Biological Sciences Research Council (BB/S007369/1). P.J.B. acknowledges funding from a Wellcome Investigator Award (214259/Z/18/Z). V.C. is funded by a personal Wellcome Early-Career Award (310356/Z/24/Z). W.S. acknowledges funding from the UKRI Future Leaders Fellowship (UKRI2045) and EPSRC (EP/W021609/1) equipment grant. D.Z is funded by a Chinese Scholarship Council (CSC) Scholarship. We thank Ryan Brady for assistance with analysis of the Tag/Catcher reconstitution assay and David Staunton for the DSC analysis at the Molecular Biophysics Suite at the University of Oxford.

## Author contributions

D.H, A.H.K, M.R.H., and E.R. designed the project; D.H., M.E., B.R.L., V.C., and D.Z. performed mass spectrometry experiments and analysis; D.H., A.H.K., P.H. and H.H. performed biophysical experiments and reactivity analysis; B.R.L. performed all work on ribosome nascent chain complexes; D.H., M.E., A.H.K., D.Z., and E.R. cloned, purified, and characterized all other protein constructs; W.S., V.C., P.J.B., M.R.H. and E.R. supervised and/or financially supported the project; D.H., M.E., A.H.K., M.R.H. and E.R. wrote the manuscript with input from the other authors.

## Materials and methods

### Plasmid constructs

pET28a-SpyTag-MBP (Addgene plasmid ID 35050)^17^, pET28a-SpyTag002-MBP (GenBank MF974389, Addgene plasmid ID 102831)^61^, and pET28a-SpyTag003-MBP (GenBank MN433888, Addgene plasmid ID 133450)^20^ were used as initial SpyTag constructs. pET28a AviTag-SpyTag002-MBP has been described^61^. pET28a-LemonTag-MBP (GenBank deposition in progress, Addgene deposition in progress) and pET28a-LemonTag(PA)-MBP (GenBank deposition in progress) were generated from pET28a-SpyTag002-MBP by site-directed mutagenesis. pDEST14-SpyCatcher (GenBank JQ478411, Addgene plasmid ID 35044)^17^, pDEST14-SpyCatcher002 (GenBank MF974388, Addgene plasmid ID 102827)^61^, and pDEST14-SpyCatcher003 (GenBank MN433887, Addgene plasmid ID 133447)^20^ were used as initial SpyCatcher constructs. pDEST14-LemonCatcher (GenBank deposition in progress, Addgene deposition in progress) was generated from pDEST14-SpyCatcher003 by site-directed mutagenesis. pET28a-MBP-sTEV (GenBank MZ365307, Addgene plasmid ID 171782)^75^ encodes an MBP fusion to superTEV protease - a variant with high stability that is active without added reducing agent. pGEX-2T-GST-BirA for site-specific biotinylation^76^ was provided by Chris O’Callaghan at the University of Oxford pET28a-His_6_-GlpG(4TM)-SecM^46^ was used as initial construct for constructing pET28a-LemonTag-His_6_-GlpG(4TM)-SecM (GenBank deposition in progress, Addgene deposition in progress).

### Site-directed mutagenesis

Site-directed mutations were performed by standard PCR using Q5 High-Fidelity 2× Master Mix (New England Biolabs) using the primers provided in **Table S1**. All mutations were validated by Sanger sequencing (Eurofins Genomics). Residue numbering is based on PDB 2X5P^77^.

### Expression and purification of SpyCatcher variants and LemonCatcher

pDEST14 vector containing Catcher variants was transformed into chemically competent *E. coli* C41 (DE3) cells (a gift from Anthony Watts, University of Oxford). Single colonies were picked into 100 mL LB medium containing 100 mg/mL ampicillin and grown for 16-18 h at 37 °C and 220 rpm. 1 L LB medium with appropriate antibiotic was inoculated with 7 mL of the saturated overnight culture and grown at 37 °C and 220 rpm. At OD_600_ = 0.5-0.6, protein overexpression was induced by addition of 0.42 mM Isopropyl b-D-1-thiogalactopyranoside (IPTG) and incubated for 4-5 h at 30 °C and 220 rpm. Cells were harvested and washed with TBS (50 mM Tris-Cl, pH 7.5, 150 mM NaCl). Cells were resuspended and lysed by sonication in Ni-NTA buffer A (50 mM Tris-HCl, 300 mM NaCl, 20 mM imidazole, pH 7.5) containing protease inhibitors (Pierce protease inhibitor tablet, EDTA-free, Thermo Scientific) and 1 mM phenylmethylsulfonyl fluoride (PMSF). Cell lysate was centrifuged for 35 min at 30,000 g. Supernatant was purified with super Ni-NTA affinity resin (ProteinArk) by equilibration/wash with buffer A and elution with buffer B (buffer A + 250 mM imidazole). Purified protein was buffer-exchanged to PBS sample buffer (PBS + 10% (v/v) glycerol, pH 7.5) using a PD-10 desalting column (Cytiva). Some samples were further purified by size exclusion chromatography using a Superdex 200 Increase 10/300 GL column (Cytiva). Protein concentration was determined by absorbance measurements at 280 nm using the extinction coefficient (e = 17,420 M^-1^ cm^-1^) predicted from ExPASy ProtParam.

### Expression and purification of SpyTag-MBP variants and LemonTag-MBP

pET28a vector containing LemonTag-MBP (or other sequences along the development pathway) was used. Protein expression and purification were performed as described for LemonCatcher (*see section above*) using the appropriate antibiotic (30 mg/mL kanamycin) for the overnight and main culture.

### Expression and purification of GlpG(4TM)-RNC

The GlpG RNC was prepared with the same construct design and high-salt purification protocol as previously described^46^. Briefly, pET28 containing GlpG with an N-terminal His_6_-tag and LemonTag, and a C-terminal arrest enhanced SecM stalling sequence (FSTPVWIWWWPRIRAPP) after transmembrane (TM) helix 4 (GlpG-4TM) was transformed into *E. coli* BL21-AI^TM^ cells (Thermo Fisher). 5 mL of an overnight LB culture was added to 4 x 1 L of pre-warmed LB containing 50 µg/mL kanamycin and grown at 37 °C. When an OD_600_ of 1.8 was reached, cultures were induced with 1 mM IPTG and 0.1% arabinose, and were cooled to 30 °C. After 1.5 hours cells were washed and harvested by centrifugation at 5,000 g.

Pellets were resuspended in 20-30 mL of Lysis Buffer (50 mM HEPES-KOH (pH 7.5), 1 M KOAc, 15 mM Mg(OAc)_2_, 5% (v/v) glycerol, 5 mM EDTA, 2 mM 2-mercaptoethanol, 1 mM PMSF, 250 μg/mL chloramphenicol, and cOmplete EDTA-free protease inhibitor tablet (Roche), and then frozen cell nuggets were prepared by dripping a concentrated cell suspension dropwise into a wide container of liquid nitrogen. Nuggets were lysed in a Spex 6875 Freezer/Mill High-Capacity Cryogenic Grinder for 15 cycles and 15 cycles per second. The resulting lysate dust was thawed and diluted in ∼100 mL Lysis Buffer and RNase-free DNase (Roche). Cell debris was removed by centrifugation at 20,000 g for 30 minutes, 4 °C. The membranes were then pelleted at 150,000 g for 45 minutes, 4 °C, and the crude membrane pellet resuspended at 40 mg/mL in Solubilization Buffer (50 mM HEPES-KOH, pH 7.5, 500 mM KOAc, 15 mM Mg(OAc)_2_, 2 mM 2-mercaptoethanol, 1 mM PMSF, 20 mM imidazole) aided by homogenization. 1% (w/v) DDM was added to the membranes, and the mixture was left to solubilize for 2 hours at 4 °C. Insoluble material was removed by ultra-centrifugation at 100,000 g for 20 minutes, 4 °C.

The supernatant was then filtered through a 0.45 μm filter (Thermo Fisher Scientific), and GlpG-TM4 RNCs were purified using an AKTA Pure purification system. Sample was loaded onto a 1 mL HiTrap Nickel column equilibrated in Wash Buffer (50 mM HEPES-KOH pH 7.5, 500 mM KOAc, 15 mM Mg(OAc)_2_, 5% (v/v) glycerol, 2 mM 2-mercaptoethanol, 1 mM PMSF, 20 mM imidazole, and 0.05% (w/v) DDM). The column was washed with 20 column volumes of Wash Buffer, 10 column volumes of Wash Buffer with 50 mM imidazole and eluted using 5 column volumes of Wash Buffer with 500 mM imidazole. GlpG-4TM RNCs were directly loaded onto a 16/60 HiPrep Sephacryl S-500 HR size-exclusion column, equilibrated in Tico Buffer (10 mM HEPES-KOH, pH 7.5, 30 mM NH_4_Cl, 15 mM Mg(OAc)_2_, 5% (v/v) glycerol, 1 mM EDTA, 2 mM 2-mercaptoethanol, 1 mM PMSF, and 0.05% (w/v) DDM). The left-hand size of the elution peak was taken as this has been shown to contain the highest abundance of 70S ribosomes^78^.

Ribosomes were concentrated and the quality of sample was assessed. The absorbance ratio at 260/280 nm was monitored to assess the 70S content and its homogeneity; expected range was between 1.9-2.0, and nascent chain integrity was confirmed by Western Blot. Samples were concentrated and RNC concentration was determined by its absorbance at 260 nm. 1 A_260_ = 24 pmol/mL^79^. GlpG-4TM RNCs were flash frozen and stored at -70 °C.

### Tag/Catcher reconstitution assay

Both binding partners, i.e. LemonCatcher and LemonTag-MBP, were diluted into sample buffer (PBS + 10% (w/v) glycerol, pH 7.5) at their desired concentrations before quenching with 1:1 (v/v) quench buffer 3.0 (100 mM NaH_2_PO_4_, 10 mM TCEP, pH 2.72) to reach pH 3.0, or 1:1 (v/v) quench buffer 2.5 (100 mM NaH_2_PO_4_, 10 mM TCEP, pH 2.3) to reach pH 2.5. TCEP (final, 5mM) addition prevented S49C-mediated disulfide bond formation of Catcher dimers during Tag/Catcher reconstitution assays. pH measurements were conducted on ice using a four-point calibration (pH solutions of 1.68, 4.01, 7.01 and 10.01) with a HI-11310 pH Edge Electrode (Hanna instruments). Quenched protein binding partners were then reacted for defined time-points before being stopped by adding 1× Laemmli sample buffer (62.5 mM Tris-HCl, pH 6.8, 10% (v/v) glycerol, 2% (w/v) SDS, 0.01% (w/v) bromophenol blue) and samples were heated for 5 min at 95 °C. Samples were loaded on a 12% (w/v) Bis-Tris protein gel (NuPAGE, Thermo Scientific) and proteins were separated by electrophoresis. Proteins were stained with Quick Coomassie Stain (Generon). The rate of amidation, i.e. isopeptide bond formation, was determined based on protein band intensities analyzed by an in-house written software.

### Generation of LemonCatcher variants by error-prone PCR

The phage display vector was based on pBAD-DsbA(ss)-HA tag-SpyDock2.0 C49S-pIII (GenBank ON131078)^65^. Gibson assembly was used to generate pBAD-DsbA(ss)-HA tag-SpyCatcher003-pIII and pBAD-DsbA(ss)-HA tag-SpyCatcher003 EA-pIII (where the E77A mutation prevents isopeptide bond formation). For library creation, the vector backbone was amplified using KOD polymerase (MilliporeSigma) with oligonucleotide primers flanking SpyCatcher003 (forward primer: 5’-GGATCCAGTGGTAGCGAAAACC-3’; reverse primer: 5’-CATGGCGCCCTGATCTCG-3’). Error-prone PCR was performed on SpyCatcher003 R32V S59T V94I (forward primer: 5’-CGAGATCAGGGCGCCATG-3’; reverse primer: 5’-GGTTTTCGCTACCACTGGATCC-3’) using GeneMorph II Random Mutagenesis Kit (Agilent). Both PCR-amplified fragments were assembled by Gibson assembly using NEBuilder HiFi DNA Assembly Master Mix (New England Biolabs). Assembly reactions were run on an agarose gel. The DNA was extracted by using Monarch DNA Gel Extraction Kit (New England Biolabs) and eluted in nuclease-free water. TG1 phage display electrocompetent *E. coli* cells (Lucigen) were transformed with 250 ng of library DNA (8 aliquots of 25 mL in total). Electroporation was performed in 0.2 mm cuvettes with a Micro-Pulser (Bio-Rad) using the EC2 program. Each electroporation was immediately recovered in 1 mL recovery medium (Lucigen) and incubated for 1 h at 37 °C and 200 rpm. Recovered cells were plated onto two bioassay dishes (245 mm × 245 mm, Nunc) with LB agar containing 0.8% (w/v) glucose and 100 mg/mL carbenicillin and incubated overnight at 30 °C. Cells were carefully collected in 2×YT medium containing 0.8% (w/v) glucose and 100 mg/mL carbenicillin. Collected cells were centrifuged and stored in 2×YT medium containing 20% (v/v) glycerol at -80 °C.

### Phage production and purification

100 mL 2×YT medium containing 2% (w/v) glucose, 0.2% (w/v) glycerol and 100 µg/mL carbenicillin were inoculated from an overnight culture of the library (*E. coli* TG1). Cells were grown at 37 °C and 200 rpm until OD_600_ reached 0.5. Cultures were infected with M13KO7 helper phage (New England Biolabs) at 20× multiplicity of infection and incubated for 45 min at 37 °C and 70 rpm. Infected cells were harvested at 3,000 g and 4 °C for 10 min, subsequently resuspended in the same volume of 2×YT medium containing 0.2% (w/v) L-arabinose, 0.2% (w/v) glycerol, and 100 µg/mL carbenicillin, and then incubated for 30 min at 18 °C and 200 rpm. 100 µg/mL kanamycin was added, and the culture was incubated for 16 h at 18 °C and 200 rpm. Cells were pelleted at 4,000 g and 4 °C for 15 min, and phage was precipitated from supernatant by incubation with 4% (w/v) polyethylene glycol (approx. 8000 g/mol) (PEG8000, Thermo Fisher) and 0.5 M NaCl for 2 h on ice. Phage were pelleted at 15,000 g and 4 °C for 45 min, resuspended in PBS (137 mM NaCl, 2.7 mM KCl, 10 mM Na_2_HPO_4_, 1.8 mM KH_2_PO_4_), pH 7.5, and centrifuged again at 15,000 g and 4 °C to remove insoluble material. Phage precipitation was repeated twice, and purified phage were stored in PBS pH 7.5 + 20% (w/v) glycerol at -80 °C. Purified phage were titered relative to a serial dilution of M13KO7 by qPCR with 2× SensiMix SYBR Hi-ROX (Bioline) master mix (forward primer: 5’-ACTGATTACGGTGCTGCTATCG-3’; reverse primer: 5’-TATCACCGTCACCGACTTGAGC-3’). qPCR was performed on an ABI7500 FAST real-time PCR machine (Thermo Fisher).

### Phage survival assay

Six samples of 25 µL phage buffer (0.66× PBS + 20% (w/v) glycerol) were mixed 1:1 with quench buffer (25 µL of 100 mM NaH_2_PO_4_, pH 2.4) to bring the solutions to pH 2.5. Six control samples at pH 7.5 were made using 50 µL of phage buffer. 10^8^ plaque forming units (pfu) of M13K07 helper phage were added to each sample by adding 1 µL of stock M13K07 solution at a concentration of 10^11^ pfu/mL, with pipetting and vortexing to mix. The samples and corresponding controls were incubated on ice for 15 min, 30 min, 1 hrs, 2 hrs, 4 hrs or 8 hrs. The reactions were stopped by addition of 50 µL of neutralization buffer (1 M Tris-HCl, pH 7.6) and incubated for 5 mins to bring the reaction pH to 7.5. Each 100 µL reaction (all samples and controls) was added to 1 mL of *E. coli* TG1 cell culture at an OD600 of 0.5, and was incubated at 37 °C, 70 rpm for 45 mins. Serial dilutions of 1:10-1:10^6^ were prepared using YT medium, and 100 µL of each dilution was plated on kanamycin plates and incubated at 37 °C overnight. Colonies were counted and concentrations of phage in pfu per mL were calculated.

### Phage display

AviTag-SpyTag002-MBP equipped with a TEV protease cleavage site was enzymatically biotinylated as performed previously^61^ and used as bait to react with LemonCatcher phage library. To simulate HDX-MS quench conditions, i.e. pH 2.5 and 0 °C, the reaction was performed in 1:1 (v/v) mix of reaction buffer (PBS pH 7.5, 3% (w/v) BSA, 0.05% (v/v) Tween-20) and quench buffer (100 mM NaH_2_PO_4_, pH 2.3). Both bait and phage were quenched (brought to pH 2.5 and 0 °C) before mixing. In the first panning round, 3.0 µM bait was mixed with 1 × 10^12^ phage and reacted for 6 h at 1,000 rpm and 4 °C. The reaction was stopped by adding a large excess of unbiotinylated SpyTag002-MBP (60 µM final). Three further panning rounds were performed by reacting (i) 0.5 µM bait and 2 × 10^11^ phage for 1 h; (ii) 0.25 µM bait and 1 × 10^11^ phage for 30 min; and (iii) 0.1 µM bait and 1 × 10^11^ phage for 10 min in the presence of *E. coli* C41(DE3) cell lysate (4 mg/mL total protein concentration). Cell lysate provides diverse competing proteins to select for sequences with specific Tag/Catcher interaction. Phage were purified from unreacted bait by precipitation with 4% (w/v) PEG8000 (Thermo Fisher) and 0.5 M NaCl for 1 h on ice. Phage were pelleted at 15,000 g and 4 °C for 15 min and resuspended in 100 µL blocking buffer (PBS pH 7.5 containing 3% (w/v) BSA and 0.1% (v/v) Tween-20). The phage bound to biotinylated AviTag-SpyTag002-MBP were captured by addition of 25 µL Dynabeads Biotin Binder (Thermo Scientific) in blocking buffer and incubating at 500 rpm and 4 °C for 1 h. The beads were captured using a MagRack 6 (Cytiva) and washed with 150 µL blocking buffer. Weakly bound phage was removed by one wash with 150 µL 200 mM glycine-HCl pH 2.2, one wash with 150 µL 100 mM triethylamine pH 11.0, 3 washes with 150 µL TBS + 0.05% (v/v) Tween-20, and a last wash with 150 µL PBS pH 7.5 + 0.1% (w/v) BSA. The supernatant was removed, and phage were eluted from beads by addition of 50 µL 50 µM MBP-sTEV in PBS pH 7.5, 10% (w/v) glycerol, and 0.5 mM ethylenediamine tetraacetic acid (EDTA). TEV protease digestion was performed at 34 °C and 1,000 rpm for 2 h. Eluted phage were rescued by infection of 1 mL *E. coli* TG1 cell culture at 37 °C and 70 rpm for 45 min. 10 µL of the culture were used for plating a serial dilution on LB agar plates containing 100 µg/mL carbenicillin to quantify the eluted phage. The remaining culture was transferred into 25 mL 2×YT supplemented with 100 µg/mL carbenicillin and grown at 37 °C and 200 rpm for 16 h. Cells were harvested and, after addition of 20% (w/v) glycerol, cells were flash-frozen in liquid nitrogen and stored at -80 °C.

### Differential scanning calorimetry (DSC)

Synthetic SpyTag002 peptide (GVPTIVMVDAYKRYK) and synthetic LemonTag peptide (GVPTIVMVDAYKPYK), each containing an additional N-terminal glycine compared to the core sequence, were solid-phase synthesized by Insight Biotechnology at >95% purity. Experiments were performed with 20 mM SpyCatcher003, SpyCatcher003 triple mutant, and LemonCatcher at pH 7.5, 3.0 or 2.5. Buffers and solution conditions for the different pH were as follows: sample buffer (PBS + 10% (w/v) glycerol) for pH 7.5; 1:1 (v/v) mixture of sample buffer and quench buffer 3.0 (100 mM NaH_2_PO_4_, pH 2.72) for pH 3.0; and 1:1 (v/v) mixture of sample buffer and quench buffer 2.5 (100 mM NaH_2_PO_4_, pH 2.3) for pH 2.5. To form the covalent complex at acidic pH, LemonCatcher was pre-reacted with 40 mM SpyTag002 (pH 2.5) or 40 mM LemonTag (pH 2.5 and 3.0) under gentle agitation using a rotation mixer (Cole-Parmer) at 4 °C for 16-18 hrs, followed by dialysis with three changes of either sample buffer at pH 7.5, 3.0, or 2.5. Thermal transitions were monitored from 20 to 110 °C at a scan rate of 3 °C/min at 3 atm on a MicroCal PEAQ-DSC (Malvern). Data was processed with MicroCal PEAQ-DSC analysis software version 1.22. The appropriate blank buffer was subtracted from the experimental sample and corrected for concentration and volume, followed by baseline subtraction. The observed transition was fitted to a two-state model, to obtain the melting temperature *T*_m_ and Full Width Half Maximum using MicroCal PEAQ-DSC analysis software (version 1.22) and Origin 2019b (OriginLab).

### Intact protein electrospray-ionization mass spectrometry

Both purified LemonCatcher and LemonTag peptide (GVPTIVMVDAYKPYK) were diluted to a final concentration of 4 µM in sample buffer (1 x PBS, 10% (v/v) glycerol, pH 7.5) and quenched by 1:1 mixture in quench buffer (100 mM NaH_2_PO_4_, pH 2.3). LemonCatcher and LemonTag peptide were then mixed in a 1:1 ratio and incubated for complex formation on ice for 1 hour. The complex was injected onto an ultraperformance liquid chromatography (UPLC) system (nanoACQUITY, Waters, Wilmslow, UK) coupled to an electrospray ionization quadrupole time-of-flight (ESI-Q-ToF) mass spectrometer (Xevo G2-XS, Waters, Wilmslow, UK). The UPLC system was equipped with a Vanguard pre-column (BEH C4, 130 Å, 1.7 µm, 1.0 mm x 100 mm; Waters) kept at 20 °C. The complex was trapped and washed with solvent A [0.23% (v/v) formic acid in H_2_O, pH 2.5) at 200 µL/min for 1 min. The complex was eluted with a 1 min linear gradient from 8-90% solvent B (0.23% (v/v) formic acid in acetonitrile) at 40 µL/min. The measurement of the complex was performed in positive ion mode between 50 and 2,000 m/z. Spectra were deconvoluted with MaxEnt1 software (Waters, UK) under standard settings.

### Biotinylation of LemonCatcher

LemonCatcher was biotinylated by addition of 10 µL EZ Link^TM^ HPDP-Biotin (N-[6-(Biotinamido)hexyl]-3′-(2′-pyridyldithio)propionamide at 4 mM in dimethylsulfoxide (DMSO, Thermo Scientific) to 50 µL of LemonCatcher with a final concentration of 667 µM for two hours at room temperature. The reaction was desalted with a micro bio-spin column, Zeba Spin Desalting Columns, 7K MWCO (Thermo Scientific), pre-equilibrated in wash buffer (PBS containing 0.01% (v/v) Tween-20, pH 7.4) according to the manufacturer’s protocol. The concentration of biotinylated LemonCatcher was determined by absorbance measurements at 280 nm using the predicted extinction coefficient (□ = 17,420 M^-1^ cm^-1^) from ExPASy ProtParam.

### LemonCatcher immobilization to beads

#### Dynabeads M-280 Streptavidin

Dynabeads M-280 Streptavidin (Thermo Fisher Scientific, Oslo, Norway, REF. 11206D) were quickly vortexed and transferred to a new tube (100 mL bead suspension, ∼ 1 mg beads; binding capacity: 0.1 mg protein/mL). Beads were washed twice with 1 mL wash buffer (PBS containing 0.01% (v/v) Tween-20, pH 7.4). Beads were resuspended in wash buffer at the initial volume. Biotinylated LemonCatcher was prepared in 500 µL of wash buffer and added to the beads (with the binding capacity of 0.1 mg protein/mL). The suspension was incubated for one hour under gentle agitation (end-over-end mixing) at room temperature. The supernatant was removed and coated beads were washed four times in 1 mL wash buffer. Beads were resuspended in wash buffer using the initial volume of bead suspension and stored at 4 °C for <1 week before use.

#### CarboxyLink Agarose, Dynabeads™ MyOne Tosyl-activated and Dynabeads™ M-270 Epoxy Beads

150 mL CarboxyLink or Dynabead coupling gel (Thermo Scientific) were transferred into a Pierce spin column (Thermo Scientific) and washed 3× with 500 mL MilliQ H_2_O (centrifugation for 1 min at 500 g). 500 mL of 5 mg/mL sulfosuccinimidyl 6-[3′-(2-pyridyldithio)propionamido]hexanoate (Sulfo-LC-SPDP; prepared fresh in MilliQ H_2_O; Sigma-Aldrich) were added and incubated for 1 h with shaking at 1,500 rpm and 25 °C. Activated beads were washed 3 time with 500 mL MilliQ H_2_O before being equilibrated with 3 × 500 mL equilibration buffer [PBS, 10% (w/v) glycerol, 0.1% (v/v) Triton X-100, pH 7.5). 250 mL LemonCatcher (362.8 mM; PBS, 10% (w/v) glycerol, pH 7.5) were provided with 0.1% (v/v) Triton X-100 and 1 mM EDTA, before being added to activated beads and incubated at 1,500 rpm and 25 °C for 24 hrs. Beads were washed 3 times with 500 mL equilibration buffer. 500 mL of L-cysteine (1 mg/mL in equilibration buffer) were added and incubated for 16-18 hrs at 1,500 rpm and 25°C, to block free sulfhydryl groups. Excess L-cysteine was removed by washing 3 times with 500 mL equilibration buffer. Beads were washed 5 times with 500 mL 1:1 (v/v) sample buffer [PBS, 10% (w/v) glycerol, pH 7.5] and quench buffer (100 mM NaH_2_PO_4_, pH 2.3) to remove non-specifically bound proteins. Beads were equilibrated by washing 3 times with 500 mL sample buffer, before adding 250 mL sample buffer for storage at 4 °C.

### Testing bead capture efficiency and specificity across LemonCatcher immobilized bead types

Both LemonCatcher beads and sample (purified LemonTag-MBP and LemonTag(PA)-MBP, cell lysate of *E. coli* C41(DE3) with overexpressed LemonTag-MBP and LemonTag(PA)-MBP, or *E. coli* C41(DE3) cell lysate provided with LemonTag-MBP and LemonTag(PA)-MBP) were quenched by 1:1 (v/v) addition of quench buffer (100 mM NaH_2_PO_4_, pH 2.72; final pH 3.0). Quenched LemonCatcher beads and quenched sample were reacted for 15 min with shaking at 1,500 rpm and 4 °C (5 mL of bead suspension pre-quench in 100 mL final reaction volume). Reacted beads were spin-filtered through pre-wet Spin-X centrifuge tube filters (Costar) (centrifugations performed for 15 s at 15,000 g) or separated by magnet (DynaMag™ Spin Magnet), depending on non-magnetic or magnetic bead type, and washed.

Initial attempts using LemonCatcher beads coupled to CarboxyLink agarose resin, Dynabeads MyOne Tosyl-activated or Dynabeads M-270 Epoxy via a sulfosuccinimidyl 6-[3′-(2-pyridyldithio)propionamido]hexanoate (Sulfo-LC-SPDP) linker led to substantial co-enrichment of unwanted proteins and increased risk of false positives (**Table S2**). Attempts to eradicate this background with glycine and bovine serum albumin (BSA) blocking and subsequent washes did not yield sufficient improvements; especially due to back exchange time-constraints restricting protocol time to <15 minutes.

The biotinylated-LemonCatcher coupled to Dynabeads M-280 Streptavidin was found to produce the most robust bead-based selective tag-capture workflow. Optimization of the SelQueX method required further testing of wash and elution step compositions and timings; speed and temperature being the dominant factors to mitigate as much back exchange as possible during work-up, with all steps being performed on wet-ice (0 °C). We found that two-times 1-minute wash steps with the addition of high salt, low denaturant and non-ionic detergent (1 M NaCl, 2 M Urea and 0.1% (w/v) DDM (ANAGRADE, CliniSciences)) improved purity. Notable, compatibility of the entire workflow with DDM detergent is important for translating the method to membrane proteins, since DDM is considered the “gold standard” for solubilization and stability^80^. The entire complex (LemonTag-MBP:LemonCatcher) was eluted by reduction with TCEP. 100 mL elution buffer (0.1-0.5 mM TCEP in 1:1 (v/v) PBS sample and quench buffer) were added to the beads separated by magnet, incubated for 1-3 min on ice with vortexing every 30 seconds. Eluant was collected and assessed by SDS-PAGE with Coomassie staining and/or liquid chromatography-mass spectrometry (LC-MS) analysis.

#### Consideration of streptavidin-biotin interaction at acidic pH

Although streptavidin undergoes substantial structural perturbations at pH < 3.0, these do not disrupt pre-existing interactions with biotinylated LemonCatcher^81^. Once biotin is bound, it stabilizes the streptavidin tetramer by strengthening inter-subunit contacts and by rigidifying flexible surface loops. As a result, the streptavidin-biotinylated LemonCatcher complex remains intact throughout the acidic capture and wash steps (as observed by others^29^) and dissociates only upon boiling in Laemmli loading buffer (**Fig. 3b**, boiled lane). In contrast, streptavidin is poorly suited as a primary capture reagent under these conditions because capture requires binding to apo-streptavidin, which is structurally perturbed at low pH^81^. Acid-induced rearrangements of the Asp61-Ser69 loop, disruption of inter-subunit hydrogen bonds, and conformational changes within the empty biotin-binding pocket reduce tetramer stability and are expected to slow association with a singly biotinylated target. Thus, while biotin-bound streptavidin remains sufficiently stable to retain captured proteins during quench and wash steps, apo-streptavidin is unlikely to support efficient target capture under the same acidic conditions. While multivalent biotinylation could improve capture through avidity effects^14^, it would require multiple efficiently labelled and accessible biotin sites, increasing tag complexity, labelling heterogeneity, and the risk of perturbing protein function. As such, this approach is impractical for a broadly applicable recombinant tagging strategy.

### SelQueX on LemonTag-MBP expressed within *E. coli* cells

LemonTag-MBP was expressed as described above. Briefly, a previously constructed pET28a plasmid was transformed into OverExpress^TM^ C41(DE3) *E. coli* cells (Sigma Aldrich) and overexpressed as previously described^21^. Overexpression was induced by adding 0.42 mM IPTG at OD_600_ of 0.5-0.6 at 37 °C in the presence of maltose at a final concentration of 0.4 % (w/v) (2 mL of maltose 20 % (w/v) was added to 100 mL of the growing cells). When the culture without maltose treatment was prepared, the same amount of H_2_O was added to the sample without maltose. Cells were then harvested at 4,500 g at 4 °C for 30 minutes, washed with 50 mL LB/H_2_O buffer, pH 7.2 (2.5 g/L LB in 50 mM NaPi and 150 mM NaCl, pH 7.2) and centrifuged at 4,500 g at 4 °C for 15 minutes. Cells were subsequently resuspended in LB/H_2_O buffer, pH 7.2 to reach the OD_600_ = 40-50; buffer was supplemented with PMSF (with a final concentration 0.5 mM) and EDTA-free protease cocktail (cOmplete™, Mini, EDTA-free Protease Inhibitor Cocktail; added at ¼ tablet to 1-3 mL lysis) to prevent endogenous proteases from breaking down and degrading the target protein after cell lysis.

#### Non-deuterated samples

200 µL of cell suspension were transferred to a new tube and 800 µL of LB/H_2_O buffer were added and vortexed. 500 µL of quench buffer (600 mM NaH_2_PO4, pH 2.5) was added to the sample and centrifuged at 4,500 g at 0 °C for 1 minute. The cell pellet was then resuspended in 100 µL of bead interaction buffer (containing 1:1 (v/v) of 100 mM NaH_2_PO_4_ with pH 2.7:PBS with pH 7.4, 0.1% (w/v) DDM). The resuspended sample was added to a 2 mL Eppendorf Safe-Lock tubes with a single 5 mm stainless steel grinding ball (Retsch) precooled using liquid N_2_ to prepare the cell nuggets. The tubes were loosely capped to allow excess liquid N_2_ to fully evaporate and held in a cryogenic box overnight at -80 °C, and the caps were tightened the next morning, as recommended by LaCava *et al.*^82^. Warning: this step is necessary to prevent excessive pressure from any remnant liquid N_2_ causing the tube to explode during storage of milling, whereas not closing the tubes after the liquid N_2_ has dissipated may result in the accumulation of frost on the cell material adding excess water weight.

Labelling proteins *in vivo* by HDX provides an appropriate lysis strategy that prevents extensive back exchange, as was previously demonstrated successfully by the Sosnick laboratory^8^. Due to the extremely low temperatures during lysis (-196°C, liquid nitrogen) and milled cell-powder storage (-80°C) then there is practically no back exchange during both stages, if the samples are kept in storage for <4 weeks before analysis^70^. After the cell suspension is quenched (by 2:1 mixture of cell suspension to quench buffer), the cells are immediately pelleted and resuspended in complete aqueous quench before being flash-frozen in preparation for cryo-milling, effectively eliminating “in-exchange” artefacts by removing remaining D_2_O content during SelQueX work-up.

#### Deuterated samples

These samples were processed in the same manner as for the non-deuterated above, except that 200 µL of cell suspension were transferred to a new tube and 800 µL of LB/D_2_O (100 atom % D, AcroSeal™, Thermo Scientific Chemicals) buffer, pH 7.2 (25g/L LB in 50mM NaPi and 150 mM NaCl in D_2_O, pD 7.2 (pH_read_ = 6.8), with or without 1 mM maltose) was added and vortexed. To measure pD, a standard pH meter was calibrated with conventional aqueous pH buffers (four-point calibration, including standards with pH 1.68, 4.01, 7.01 and 10.01), we took the reading in the D₂O buffer and added a correction factor of +0.40 (pD = pH_read_ + 0.40)^2^. We performed cellular HDX on *E. coli* cells with overexpressed LemonTag-MBP -grown with or without 1 mM maltose present - across 1-, 5- and 30-minute HDX time points at 22 °C. All the experiments and steps were performed and repeated in at least triplicate from individual transformants/colonies (representing biological replicates) for both overexpressed samples with and without maltose.

Frozen cell pellets were disrupted under cryogenic conditions using a Retsch CryoMill (Retsch GmbH, 42781 Haan, Germany), *Model:* 20.749.0001 (100-240 V, 50/60 Hz with 5-30 Hz vibrational frequency). The chamber was pre-cooled for 5 minutes with liquid nitrogen (LN₂) to -196 °C before milling begins using the automated pre-cool function. Cell disruption was performed for 5 cycles of 3 minutes at 25 s^-1^ (run) and 1.5 minutes at 5 s^-1^ (break).

For the complex pull-down step, 40 µL of LemonCatcher immobilized Dynabeads M-280 Streptavidin (“LemonCatcher beads”) bead suspension was transferred to a new tube and the supernatant was removed. 200 µL of interaction buffer pre-chilled to 0 °C (on wet ice) (containing 1:1 (v/v) of 100 mM NaH_2_PO_4_, pH 2.7:PBS pH 7.4, 0.1% (w/v) DDM - at the final pH 3.0) was added to the frozen cell dust and rapidly vortexed to thaw and centrifuged at 16,000 g at 0 °C for 1 minute. 200 µL of the supernatant was transferred to the LemonCatcher beads and incubated under shaking at 0 °C (on wet ice) for 5 minutes, with light vortexing every 30 seconds to prevent bead clumping. The supernatant was removed and beads were washed twice with 200 mL wash buffer (containing 50 mM NaH_2_PO_4_, 2 M urea, 1 M NaCl, 0.1% (w/v) DDM with pH 2.7). 100 mL of elution buffer (containing 500 mM TCEP in 4 M Urea, 0.2% (v/v) formic acid and 0.1% (w/v) DDM with pH 3.0) was added to the beads and incubated on ice for 3 minutes under gentle agitation every 10 seconds. Fortunately, urea slows the base-catalyzed amide hydrogen exchange pathway and shifts the optimal quench pH upward, resulting in only minimal back-exchange differences between pH 2.5 and 3.0 in urea-TCEP buffers^30^. The supernatant containing the eluate of the LemonTag-MBP:LemonCatcher was transferred to a pre-cooled tube and used for downstream SDS-PAGE or HDX-MS analysis (samples were stored at -80 °C for <4 weeks before further processing with LC-MS).

### Hydrogen-deuterium exchange mass spectrometry analysis

#### In-cell LemonTag-MBP measurements

Samples (eluted LemonTag-MBP:LemonCatcher complex) were injected as is or diluted with Solvent A (0.23% (v/v) formic acid in H_2_O (Optima™ LC/MS Grade, Fisher Chemical™)) at a 2:3 ratio (v/v) of Solvent A:sample to decrease the TCEP and buffer component concentrations. Samples were injected into an UPLC system (nanoACQUITY, Waters, Wilmslow, UK) coupled to an electrospray ionization quadrupole time-of-flight (ESI-Q-ToF) mass spectrometer (Synapt XS, Synapt G2-Si, or Xevo G2-XS, Waters, Wilmslow, UK).

The HDX manager of the nanoACQUITY system was equipped with a Vanguard column (BEH C18, 130 Å, 1.7 μm, 2.1 mm × 5 mm; Waters) and an Acquity UPLC column (BEH C18, 130 Å, 1.7 μm, 1.0 mm × 100 mm; Waters) for peptide trapping and separation, respectively. Protein digestion was performed online with the UPLC system using an in-house packed protease column (immobilized pepsin agarose resin, Thermo Scientific) at 20 °C. The generated peptides were trapped and washed with solvent A (0.23% (v/v) formic acid in H_2_O, pH 2.5) at 200 mL/min for 3 or 4 min (depending on experiment). Subsequently, peptides were separated by applying a 7.5 min linear gradient from 8 to 35% solvent B (0.23% (v/v) formic acid in acetonitrile) at 40 mL/min on the Xevo G2-XS mass spectrometer. Peptides were separated by applying a 7-min linear gradient from 8 to 35% solvent B (0.23% (v/v) formic acid in acetonitrile) at 85 mL/min on the Synapt G2Si or Synapt XS mass spectrometers. Peptides were measured in positive ion mode between 50 and 2,000 m/z on the Xevo G2-XS mass spectrometer. Peptides were measured in positive ion mode between 50 and 2,000 m/z on the Synapt G2Si or Synapt XS mass spectrometers applying ion mobility separation. For comparison, purified LemonCatcher and LemonTag-MBP were separately measured in triplicates under the same buffer conditions and instrument settings.

#### HDX-MS of RNCs

30 µL of GlpG-4TM RNC (0.4-1.0 µM) was incubated with 1 µL Tico Buffer or RNase (1/10 dilution from 10 mg/mL stock) for 5 minutes at room temperature. Samples were then diluted in Tico Buffer made up with MilliQ water or D_2_O (100 atom % D, AcroSeal™, Thermo Scientific Chemicals) and incubated at 25 °C for 5 minutes. The reaction was quenched 1:1 with interaction buffer to bring the final pH to 3.0, and 40 µL of LemonCatcher beads (see ‘LemonCatcher immobilization to beads’) was added. The sample was left on ice for 5 minutes and vortexed every 30 s. The supernatant was removed, and the beads were washed twice with wash buffer. 60 µL elution buffer was added to the beads and incubated on ice for 3 minutes, with gentle agitation every 10 seconds. Supernatant was transferred to a pre-cooled tube and flash-frozen by plunging into liquid nitrogen and stored at -70 °C, before injection into the mass spectrometer within 4 weeks.

Samples were thawed and injected into a nanoAcquity UPLC system (Waters) coupled to a Synapt G2-Si mass spectrometer (Waters, UK). 60 µL were injected into a 50 µL sample loop before being injected onto an online Enzymate™ pepsin digestion column (Waters) equilibrated in Solvent A (0.23% (v/v) formic acid in water) at 200 µL/min. The peptic fragments were trapped onto an Acquity Vanguard pre-column (BEH C18, 130 Å, 1.7 μm, 2.1 mm × 5 mm; Waters) for 3 minutes and washed with Solvent A at 200 µL/min. Peptides were eluted by applying an 8-35% linear gradient of Solvent B (0.23% formic acid in acetonitrile) at 40 µL/min and separated in an Acquity UPLC column BEH C18, 130 Å, 1.7 μm, 1.0 mm × 100 mm; Waters). The trap and UPLC columns were both maintained at 0 °C. The eluted peptides were ionized by electrospray measured in positive ion mode between 50 and 2,000 *m/z* on the Synapt G2-Si mass spectrometer in ion mobility mode. Leucine enkephalin was used for lockmass accuracy correction and the mass spectrometer calibrated with sodium iodide. Each sample was measured under identical conditions in at least triplicate, and a blank run of pepsin wash (1.5M guanidinium hydrochloride, 4% (v/v) acetonitrile, 0.8% (v/v) formic acid), was performed between each sample to prevent significant peptide carryover.

### HDX-MS data processing

#### In-cell LemonTag-MBP measurements

Protein identification and peptide filtering were performed with ProteinLynx Global Server 3.0 (PLGS) and DynamX 3.0, respectively (Waters, Wilmslow, UK). PLGS workflow parameters for peptide identification were as follows: peptide tolerance: automatic; fragment tolerance: automatic; min fragment ion matches per peptide: 2; minimum fragment ion matches per protein: 7; minimum peptide matches per protein: 3; maximum protein mass 250,000 Da; primary digest reagent: nonspecific; and false discovery rate: 100%. PLGS output files were then consulted for further validation by DynamX. DynamX parameters for peptide filtering were as follows^83^: (for Xevo G2-XS measurements) minimum intensity: 1,481; minimum sequence length: 5; maximum sequence length: 25; minimum products per amino acid: 0.11; minimum consecutive products: 1; minimum PLGS score: 6.62; maximum MH+ error (ppm): 5 | (for Synapt G2-Si or Synapt G2-XS measurements): peptides identified in 3 out of 4 runs with minimum products: 2, minimum products per amino acid: 0.2; minimum consecutive products: 1; maximum MH+ error (ppm): 7.. All the spectra were visually examined and only those with a suitable signal to noise ratio were used for analysis.

The amount of relative deuterium uptake for each peptide was determined using DynamX (v. 3.0) and only corrected for back exchange when specified. The relative fractional uptake (RFU) was calculated from the following equation, where *Y* is the deuterium uptake for peptide *a* at incubation time (*t*), and *D* is the percentage of deuterium in the final labelling solution: RFU_a_ = Y_a,t_ / MaxUptake_a_ × *D*_max_. *D*_max_ was collected using the protocol established by Peterle *et al.*^84^.

#### RNC measurements

Peptide were identified from high-definition MS^E^ analyses of non-deuterated GlpG(4TM)-RNC samples, using PLGS. PLGS workflow parameters are the same as described in the HDX-MS data processing section. PLGS output files were then consolidated in DynamX 3.0 for further validation, using previously published parameters for HDX-MS of soluble RNCs^41^: maximum intensity 1000; minimum sequence length 5; maximum sequence length 40; minimum products per amino acid 0.05; minimum consecutive products 1. Each peptide had to be found in 2/3 references files, and all spectra were visually examined, with only those with a suitable signal to noise ratio being used for analysis.

### HDX-MS quantification and statistical analysis

For in-cell HDX-MS analysis on LemonTag-MBP a significance level cut-off for experiments on biological replicates (*n* = 3) was set at p ≤ 0.05; ΔHDX of 0.40 Da. For HDX-MS analysis on LomeTag-GlpG(4TM)-RNCs a significance level cut-off for experiments on biological replicates (*n* = 3) was set at p ≤ 0.05; ΔHDX of 0.67 Da. This deuterium difference was greater than the confidence intervals calculated based on the pooled standard deviation values^85^. The HDX-MS data and the HDX-MS summary tables generated in this study have been deposited to ProteomeXchange (deposition currently private) and provided in **Table S3** and the HDX uptake plots as **Supporting Data** as per consensus guidelines.

### Protein structure analysis

Protein structures were displayed in PyMOL version 2.0.6 (DeLano Scientific), based on PDB 4MLI^86^. The model of LemonTag:LemonCatcher was created using AlphaFold 3 using the AlphaFold Server^49^ and the electrostatic maps visualized in ChimeraX^87^ The final AlphaFold model used had the flexible termini of the Tag and Catcher truncated. The isopeptide bond was not recapitulated in the AlphaFold model.

## References

1. Beck, M., Covino, R., Hänelt, I. & Müller-McNicoll, M. Understanding the cell: Future views of structural biology. Cell 187, 545–562 (2024).

2. Masson, G. R. et al. Recommendations for performing, interpreting and reporting hydrogen deuterium exchange mass spectrometry (HDX-MS) experiments. Nat. Methods 16, 595–602 (2019).

3. Engen, J. R. & Komives, E. A. Complementarity of Hydrogen/Deuterium Exchange Mass Spectrometry and Cryo-Electron Microscopy. Trends Biochem. Sci. 45, 906–918 (2020).

4. Platzer, G., Mayer, M., McConnell, D. B. & Konrat, R. NMR-driven structure-based drug discovery by unveiling molecular interactions. Commun. Chem. 8, 167 (2025).

5. Hamuro, Y. Interpretation of Hydrogen/Deuterium Exchange Mass Spectrometry. J. Am. Soc. Mass Spectrom. 35, 819–828 (2024).

6. Kaldmäe, M. et al. High intracellular stability of the spidroin N-terminal domain in spite of abundant amyloidogenic segments revealed by in-cell hydrogen/deuterium exchange mass spectrometry. FEBS J. 287, 2823–2833 (2020).

7. Ghaemmaghami, S. & Oas, T. G. Quantitative protein stability measurement in vivo. Nat. Struct. Biol. 8, 879–882 (2001).

8. Lin, X., Zmyslowski, A. M., Gagnon, I. A., Nakamoto, R. K. & Sosnick, T. R. Development of in vivo HDX-MS with applications to a TonB-dependent transporter and other proteins. Protein Science 31, e4402–e4402 (2022).

9. James, E. I., Murphree, T. A., Vorauer, C., Engen, J. R. & Guttman, M. Advances in Hydrogen/Deuterium Exchange Mass Spectrometry and the Pursuit of Challenging Biological Systems. Chem. Rev. 122, 7562–7623 (2022).

10. Cheung, R. C. F., Wong, J. H. & Ng, T. B. Immobilized metal ion affinity chromatography: a review on its applications. Appl. Microbiol. Biotechnol. 96, 1411– 1420 (2012).

11. Hartley, R. W. [38] - Barnase–Barstar Interaction. in Methods in Enzymology (ed. Nicholson, A. W.) vol. 341 599–611 (Academic Press, 2001).

12. Aghayeva, U. F., Nikitin, M. P., Lukash, S. V & Deyev, S. M. Denaturation-Resistant Bifunctional Colloidal Superstructures Assembled via the Proteinaceous Barnase– Barstar Interface. ACS Nano 7, 950–961 (2013).

13. Ejima, D. et al. Effects of acid exposure on the conformation, stability, and aggregation of monoclonal antibodies. Proteins: Structure, Function, and Bioinformatics 66, 954– 962 (2007).

14. Jensen, P. F., Jorgensen, T. J., Koefoed, K., Nygaard, F. & Sen, J. W. Affinity capture of biotinylated proteins at acidic conditions to facilitate hydrogen/deuterium exchange mass spectrometry analysis of multimeric protein complexes. Anal Chem 85, 7052–7059 (2013).

15. Li, Y. & Sousa, R. Expression and purification of E. coli BirA biotin ligase for in vitro biotinylation. Protein Expr. Purif. 82, 162–167 (2012).

16. Fairhead, M. & Howarth, M. Site-Specific Biotinylation of Purified Proteins Using BirA. in Site-Specific Protein Labeling: Methods and Protocols (eds. Gautier, A. & Hinner, M. J.) 171–184 (Springer New York, New York, NY, 2015). doi:10.1007/978-1-4939-2272-7_12.

17. Zakeri, B. et al. Peptide tag forming a rapid covalent bond to a protein, through engineering a bacterial adhesin. Proceedings of the National Academy of Sciences 109, E690–E697 (2012).

18. Howarth, M. R. Click biology highlights the opportunities from reliable biological reactions. Nat. Chem. Biol. 21, 991–1005 (2025).

19. Keeble, A. H. & Howarth, M. Power to the protein: enhancing and combining activities using the Spy toolbox. Chem. Sci. 11, 7281–7291 (2020).

20. Keeble, A. H. et al. Approaching infinite affinity through engineering of peptide–protein interaction. Proceedings of the National Academy of Sciences 116, 26523–26533 (2019).

21. Keeble, A. H., et al. Evolving Accelerated Amidation by SpyTag/SpyCatcher to Analyze Membrane Dynamics. Angew. Chem. Int. Ed. 56, 16521–16525 (2017).

22. Fink, A. L., Calciano, L. J., Goto, Y., Kurotsu, T. & Palleros, D. R. Classification of Acid Denaturation of Proteins: Intermediates and Unfolded States. Biochemistry 33, 12504– 12511 (1994).

23. Grimsley, G. R., Scholtz, J. M. & Pace, C. N. A summary of the measured pK values of the ionizable groups in folded proteins. Protein Science 18, 247–251 (2009).

24. Li, L., Fierer, J. O., Rapoport, T. A. & Howarth, M. Structural Analysis and Optimization of the Covalent Association between SpyCatcher and a Peptide Tag. J. Mol. Biol. 426, 309–317 (2014).

25. Smith, G. P. Filamentous Fusion Phage: Novel Expression Vectors That Display Cloned Antigens on the Virion Surface. Science 228, 1315–1317 (1985).

26. Martínez-Nicolas, J. J. et al. Physico-Chemical Attributes of Lemon Fruits as Affected by Growing Substrate and Rootstock. Foods 11, (2022).

27. Hansen, A. B., Rouhi, O. & Rand, K. D. Developments in hydrogen/deuterium exchange mass spectrometry at sub-zero temperatures. TrAC Trends in Analytical Chemistry 194, 118539 (2026).

28. Masson, G. R. et al. Recommendations for performing, interpreting and reporting hydrogen deuterium exchange mass spectrometry (HDX-MS) experiments. Nat. Methods 16, 595–602 (2019).

29. Vorauer, C. et al. Direct Mapping of Polyclonal Epitopes in Serum by HDX-MS. Anal. Chem. 96, 16758–16767 (2024).

30. Hamuro, Y. & Coales, S. J. Optimization of Feasibility Stage for Hydrogen/Deuterium Exchange Mass Spectrometry. J Am Soc Mass Spectrom 29, 623–629 (2018).

31. Ahn, J., Cao, M.-J., Yu, Y. Q. & Engen, J. R. Accessing the reproducibility and specificity of pepsin and other aspartic proteases. Biochimica et Biophysica Acta (BBA) - Proteins and Proteomics 1834, 1222–1229 (2013).

32. Rey, M. et al. Nepenthesin from Monkey Cups for Hydrogen/Deuterium Exchange Mass Spectrometry*. Molecular & Cellular Proteomics 12, 464–472 (2013).

33. Hamuro, Y. & Zhang, T. High-Resolution HDX-MS of Cytochrome c Using Pepsin/Fungal Protease Type XIII Mixed Bed Column. J. Am. Soc. Mass Spectrom. 30, 227–234 (2019).

34. Ahn, J., Jung, M. C., Wyndham, K., Yu, Y. Q. & Engen, J. R. Pepsin Immobilized on High-Strength Hybrid Particles for Continuous Flow Online Digestion at 10 000 psi. Anal. Chem. 84, 7256–7262 (2012).

35. Telmer, P. G. & Shilton, B. H. Insights into the Conformational Equilibria of Maltose-binding Protein by Analysis of High Affinity Mutants*. Journal of Biological Chemistry 278, 34555–34567 (2003).

36. Sun, P., Tropea, J. E. & Waugh, D. S. Enhancing the Solubility of Recombinant Proteins in Escherichia coli by Using Hexahistidine-Tagged Maltose-Binding Protein as a Fusion Partner. in Heterologous Gene Expression in E.coli: Methods and Protocols (eds. Evans Thomas C., Jr. & Xu, M.-Q.) 259–274 (Humana Press, Totowa, NJ, 2011). doi:10.1007/978-1-61737-967-3_16.

37. Winfried, B. & Howard, S. Maltose/Maltodextrin System of Escherichia coli: Transport, Metabolism, and Regulation. Microbiology and Molecular Biology Reviews 62, 204– 229 (1998).

38. Liu, T., Limpikirati, P. & Vachet, R. W. Synergistic Structural Information from Covalent Labeling and Hydrogen–Deuterium Exchange Mass Spectrometry for Protein–Ligand Interactions. Anal. Chem. 91, 15248–15254 (2019).

39. Millet, O., Hudson, R. P. & Kay, L. E. The energetic cost of domain reorientation in maltose-binding protein as studied by NMR and fluorescence spectroscopy. Proceedings of the National Academy of Sciences 100, 12700–12705 (2003).

40. Quiocho, F. A., Spurlino, J. C. & Rodseth, L. E. Extensive features of tight oligosaccharide binding revealed in high-resolution structures of the maltodextrin transport/chemosensory receptor. Structure 5, 997–1015 (1997).

41. Pellowe, G. A. et al. The human ribosome modulates multidomain protein biogenesis by delaying cotranslational domain docking. Nat. Struct. Mol. Biol. 32, 2296–2307 (2025).

42. Wales, T. E. et al. Resolving chaperone-assisted protein folding on the ribosome at the peptide level. Nat. Struct. Mol. Biol. 31, 1888–1897 (2024).

43. Lin, X., Molina, A. V, Shangguan, J., Chen, R. & Sosnick, T. R. Practical Tips for the Application of HDX-MS to Membrane Proteins, Biomolecular Condensates, and Weak Protein Binders. J. Am. Soc. Mass Spectrom. https://doi.org/10.1021/jasms.5c00067 (2025) doi:10.1021/jasms.5c00067.

44. Roeselová, A. et al. Hydrogen/deuterium exchange mass spectrometry analysis of ribosome-nascent chain complexes to study protein biogenesis at the peptide level. Nat. Protoc. 21, 2271–2300 (2026).

45. Russell CW, Richards AC, Chang AS, Mulvey MA. The Rhomboid Protease GlpG Promotes the Persistence of Extraintestinal Pathogenic Escherichia coli within the Gut. Infect. Immun. 85, 10.1128/iai.00866-16 (2017).

46. Pellowe, G. A. et al. Capturing Membrane Protein Ribosome Nascent Chain Complexes in a Native-like Environment for Co-translational Studies. Biochemistry 59, 2764–2775 (2020).

47. Chan, S. H. S. et al. The ribosome stabilizes partially folded intermediates of a nascent multi-domain protein. Nat. Chem. 14, 1165–1173 (2022).

48. Masse, M. M. et al. Nascent chains derived from a foldable protein sequence interact with specific ribosomal surface sites near the exit tunnel. Sci. Rep. 14, 12324 (2024).

49. Abramson, J. et al. Accurate structure prediction of biomolecular interactions with AlphaFold 3. Nature 630, 493–500 (2024).

50. Watson, J. L. et al. De novo design of protein structure and function with RFdiffusion. Nature 620, 1089–1100 (2023).

51. Irobalieva, R. N., Martins, B. & Medalia, O. Cellular structural biology as revealed by cryo-electron tomography. J Cell Sci 129, 469–476 (2016).

52. Luchinat, E. & Banci, L. A Unique Tool for Cellular Structural Biology: In-cell NMR. J. Biol. Chem. 291, 3776–3784 (2016).

53. Bucher, D., Grant, B. J., Markwick, P. R. & McCammon, J. A. Accessing a Hidden Conformation of the Maltose Binding Protein Using Accelerated Molecular Dynamics. PLoS Comput. Biol. 7, e1002034- (2011).

54. Shen, Y. & Bax, A. Two-conformer equilibrium of maltose-binding protein in the absence of ligand from residual dipolar coupling analysis. Protein Science 35, e70425 (2026).

55. Tang, C., Schwieters, C. D. & Clore, G. M. Open-to-closed transition in apo maltose-binding protein observed by paramagnetic NMR. Nature 449, 1078–1082 (2007).

56. Walker, I. H., Hsieh, P. & Riggs, P. D. Mutations in maltose-binding protein that alter affinity and solubility properties. Appl. Microbiol. Biotechnol. 88, 187–197 (2010).

57. Masson, G. R., Jenkins, M. L. & Burke, J. E. An overview of hydrogen deuterium exchange mass spectrometry (HDX-MS) in drug discovery. Expert Opin Drug Discov 12, 981–994 (2017).

58. Krishnamurthy, S., Musgaard, M., Tehan, B. G., Jazayeri, A. & Liko, I. The evolving role of hydrogen/deuterium exchange mass spectrometry in early-stage drug discovery. Curr. Opin. Struct. Biol. 92, 103051 (2025).

59. F. Malta, C., et al. Pushing the limits of hydrogen/deuterium exchange mass spectrometry to study protein:fragment low affinity interactions. Commun. Chem. 8, 405 (2025).

60. Rekadwad, B. N. et al. Extremophiles: the species that evolve and survive under hostile conditions. 3 Biotech 13, 316 (2023).

61. Keeble, A. H., et al. Evolving Accelerated Amidation by SpyTag/SpyCatcher to Analyze Membrane Dynamics. Angew. Chem. Int. Ed. 56, 16521–16525 (2017).

62. Hagan, R. M., et al. NMR Spectroscopic and Theoretical Analysis of a Spontaneously Formed Lys–Asp Isopeptide Bond. Angew. Chem. Int. Ed. 49, 8421–8425 (2010).

63. Isom, D. G., Castañeda, C. A., Cannon, B. R. & García-Moreno E., B. Large shifts in pKa values of lysine residues buried inside a protein. Proceedings of the National Academy of Sciences 108, 5260–5265 (2011).

64. Pettinger, J., Jones, K. & Cheeseman, M. D. Lysine-Targeting Covalent Inhibitors. Angew. Chem. Int. Ed. 56, 15200–15209 (2017).

65. Vester, S. K. et al. SpySwitch enables pH- or heat-responsive capture and release for plug-and-display nanoassembly. Nat. Commun. 13, 3714 (2022).

66. Yue, H., Zhao, Y., Ma, X. & Gong, J. Ethylene glycol: properties, synthesis, and applications. Chem. Soc. Rev. 41, 4218–4244 (2012).

67. Svejdal, R. R., Dickinson, E. R., Sticker, D., Kutter, J. P. & Rand, K. D. Thiol-ene Microfluidic Chip for Performing Hydrogen/Deuterium Exchange of Proteins at Subsecond Time Scales. Anal. Chem. 91, 1309–1317 (2019).

68. Hammerschmid, D. et al. Droplet microfluidic hydrogen/deuterium exchange for investigating protein dynamics with millisecond precision. ChemRxiv https://doi.org/doi:10.26434/chemrxiv-2025-j9k30 (2025) doi:doi:10.26434/chemrxiv-2025-j9k30.

69. Raval, S. et al. Nano-scaled, fully automated hydrogen/deuterium exchange for analysis of macromolecular assemblies reaching the MDa scale. Molecular & Cellular Proteomics 101519 (2026) 10.1016/j.mcpro.2026.101519.

70. Watson, M. J. et al. Simple Platform for Automating Decoupled LC–MS Analysis of Hydrogen/Deuterium Exchange Samples. J. Am. Soc. Mass Spectrom. 32, 597–600 (2021).

71. Filandr, F. et al. Automating data analysis for hydrogen/deuterium exchange mass spectrometry using data-independent acquisition methodology. Nat. Commun. 15, 2200 (2024).

72. Lu, C., Wells, M. L., Reckers, A., McBride, S. K. & Glasgow, A. Site-resolved energetic information from HX–MS experiments. Nat. Chem. Biol. 22, 307–317 (2026).

73. Ferrari, Á. J. R. et al. Large-scale discovery, analysis and design of protein energy landscapes. Nature https://doi.org/10.1038/s41586-026-10465-z (2026) doi:10.1038/s41586-026-10465-z.

74. Abucayon, E. G. et al. An Ultra-High Throughput Hydrogen–Deuterium Exchange Workflow Using Acoustic Ejection Mass Spectrometry for Studying Peptide Solution Structural Conformation. Anal. Chem. https://doi.org/10.1021/acs.analchem.6c00487(2026) doi:10.1021/acs.analchem.6c00487.

75. Keeble, A. H. et al. DogCatcher allows loop-friendly protein-protein ligation. Cell Chem. Biol. 29, 339–350.e10 (2022).

76. Fairhead, M. & Howarth, M. Site-Specific Biotinylation of Purified Proteins Using BirA. in Site-Specific Protein Labeling: Methods and Protocols (eds. Gautier, A. & Hinner, M. J.) 171–184 (Springer New York, New York, NY, 2015). doi:10.1007/978-1-4939-2272-7_12.

77. Oke, M. et al. The Scottish Structural Proteomics Facility: targets, methods and outputs. J. Struct. Funct. Genomics 11, 167–180 (2010).

78. Becker, M. et al. The 70S ribosome modulates the ATPase activity of Escherichia coli YchF. RNA Biol. 9, 1288–1301 (2012).

79. Cassaignau, A. M. E. et al. A strategy for co-translational folding studies of ribosome-bound nascent chain complexes using NMR spectroscopy. Nat. Protoc. 11, 1492–1507 (2016).

80. Tulumello, D. V & Deber, C. M. Efficiency of detergents at maintaining membrane protein structures in their biologically relevant forms. Biochim Biophys Acta 1818, 1351–1358 (2012).

81. Katz, B. A. Binding of biotin to streptavidin stabilizes intersubunit salt bridges between Asp61 and His87 at low pH11Edited by I. A. Wilson. J. Mol. Biol. 274, 776–800 (1997).

82. LaCava, J., Jiang, H. & Rout, M. P. Protein Complex Affinity Capture from Cryomilled Mammalian Cells. JoVE e54518 (2016) doi:doi:10.3791/54518.

83. Sørensen, L. & Salbo, R. Optimized Workflow for Selecting Peptides for HDX-MS Data Analyses. J. Am. Soc. Mass Spectrom. 29, 2278–2281 (2018).

84. Peterle, D., Wales, T. E. & Engen, J. R. Simple and Fast Maximally Deuterated Control (maxD) Preparation for Hydrogen–Deuterium Exchange Mass Spectrometry Experiments. Anal. Chem. 94, 10142–10150 (2022).

85. Hageman, T. S. & Weis, D. D. Reliable Identification of Significant Differences in Differential Hydrogen Exchange-Mass Spectrometry Measurements Using a Hybrid Significance Testing Approach. Anal. Chem. 91, 8008–8016 (2019).

86. Li, L., Fierer, J. O., Rapoport, T. A. & Howarth, M. Structural Analysis and Optimization of the Covalent Association between SpyCatcher and a Peptide Tag. J. Mol. Biol. 426, 309–317 (2014).

87. Pettersen, E. F. et al. UCSF Chimera—A visualization system for exploratory research and analysis. J. Comput. Chem. 25, 1605–1612 (2004).

