## Supplementary Information for "LemonCatcher Acidic Pull-Down Enables Selective In-Cell Hydrogen-Deuterium Exchange Mass Spectrometry"

### Supplementary Figures

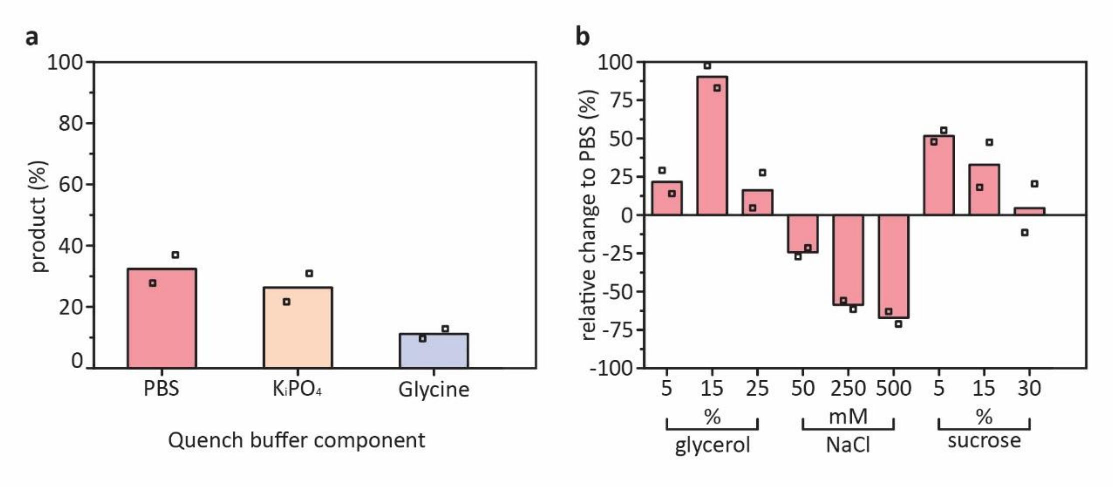


**Figure S1. Testing the influence of buffer conditions and additives on reaction efficiency of SpyTag002/SpyCatcher003 under HDX quench conditions.** *(**a**) Reconstitution product efficiency tested within commonly used HDX quench solutions that buffer at pH 2.5; performed* at *0 °C for 15 minutes using 5 µM each of SpyTag002-MBP and SpyCatcher003. PBS = quenched PBS buffer (PBS, pH 7.5 quenched to pH 2.5 by 1:1 (v/v) mixing with 100 mM $NaH_2PO_4$, pH 2.3). $KiPO_4$ = quenched potassium phosphate buffer (50 mM potassium phosphate pH 7.0 quenched to pH 2.5 by 1:1 (v/v) mixing with 100 mM $KH_2PO_4$, pH 1.8). Glycine = 50 mM Glycine pH 2.5. (**b**) Influence of additives on reconstitution product efficiency relative to average reconstitution product achieved in PBS condition. Reactions were performed as in (a) except for adding the indicated amount of additive. All reactions were performed in duplicate, with each data point shown as a square and the bar representing the mean. The additional % (v/v) of glycerol, NaCl, or % (w/v) of sucrose with PBS pH 2.5 is marked.*

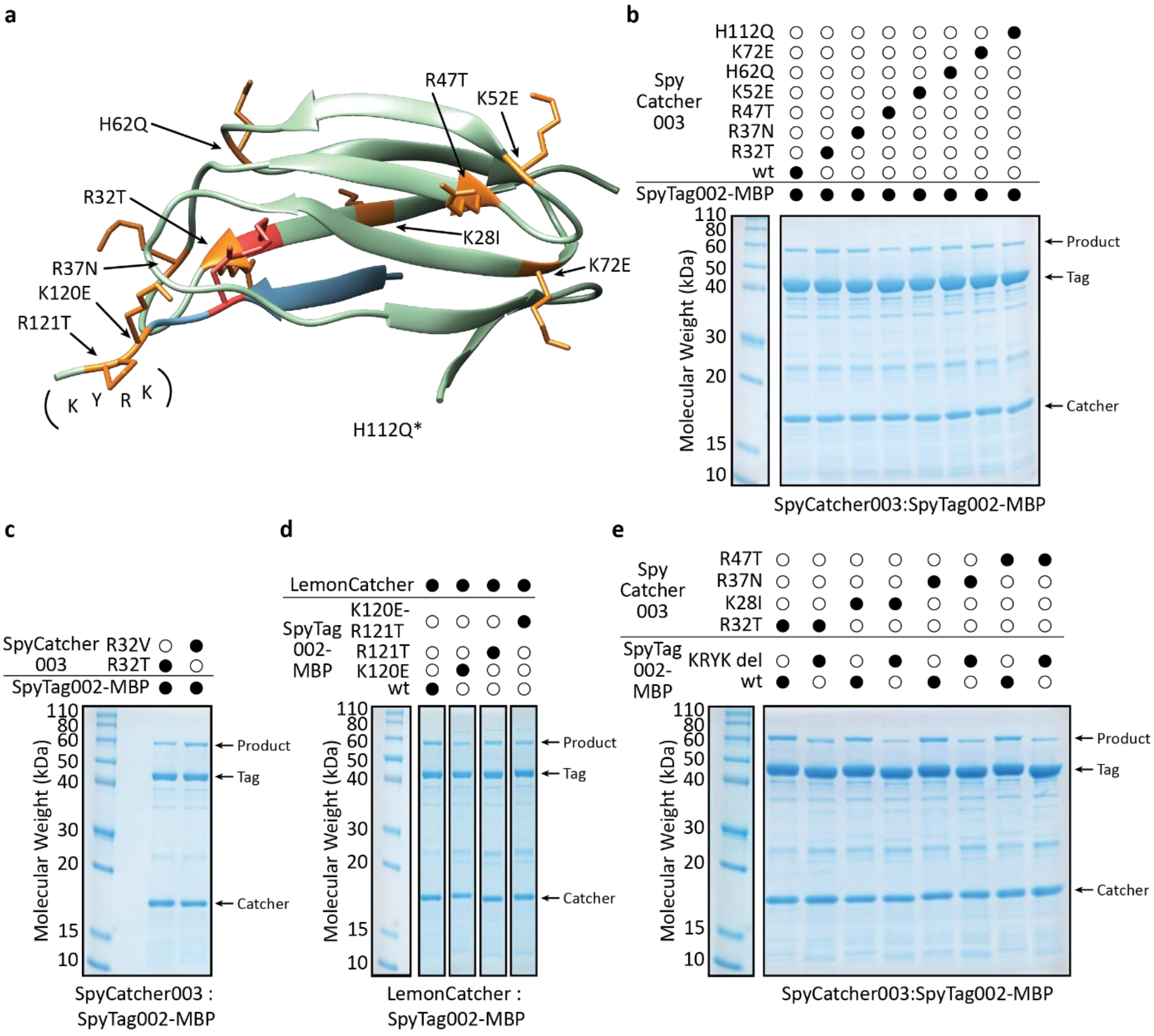


**Figure S2. Initial strategy to decrease positively charged residues to enhance Tag/Catcher reactivity in acidic HDX quench conditions.** *(**a**) Schematic of Catcher (green) and Tag (blue) based on SpyTag: SpyCatcher PDB 4MLI*[1]*. Single-point mutations trialed are highlighted, with side-chains shown in orange in stick format. *H112Q is contained within an unresolved part of the structure. (KYRK) defines the* $^{120}$*KYRK*$^{123}$ *SpyTag002 deletion mutant. (**b, c, d,** and **e**) SpyTag002-MBP and SpyCatcher003 variants were each present at 5 µM, with reaction for 15 minutes at 0 °C in quenched PBS sample buffer + 5 mM TCEP (pH 2.5), before SDS-PAGE with Coomassie staining under reducing conditions. Probable SlyD band detected at ~24 kDa, a common contaminant during IMAC purification due to its high percentage of histidine's (10.2%) which could not be completely removed after size exclusion chromatography due to its similar size to the Catcher*[2]*. Samples were only purified by IMAC before analysis.*

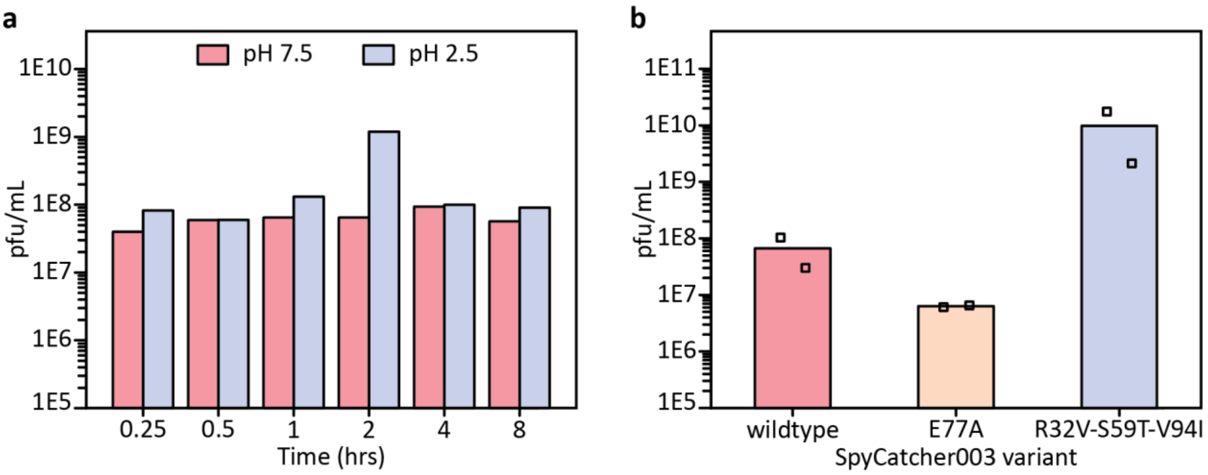


c

| Clone | Mutations relative to R32V/S59T/V94I SpyCatcher003 |
|---|---|
| L1R2C5 | D16H D22E E108V |
| L1R2C6 | E108G |
| L1R3C4 | E108D |
| L2R3C10 | D97V |
| L1R4C4 | V76D D97E E108G |
| L1R4C7 | Y69H D97E G104D |
| L1R4C11 | T56A N95T |
| L1R4C13 | N95S |

| | Additions to R32V/S59T/V94I SpyCatcher003 tested | | | | |
|---|---|---|---|---|---|
| Mutation | T56A | V76I | N95T | D97V | E108G |
| Rxn effect | ↑ | – | ↑ | – | – |

**Figure S3. Phage-display workflow for directed evolution within HDX quench-like conditions.** ***(a)*** *Phage remains infective over hours at pH 2.5: M13K07 helper phage was incubated at pH 2.5 or 7.5 at 4 °C for the indicated time, before titering of surviving phage by E. coli infection.* ***(b)*** *Model selection for phage panning. Phage displaying wildtype, E77A or R32V-S59T-V94I SpyCatcher003 was incubated with biotinylated SpyTag002-MBP and pulled down on streptavidin-beads, before quantifying as plaque-forming units (pfu) (mean with each of duplicate values marked). E77A stops isopeptide bond formation but does not prevent the initial non-covalent Tag/Catcher association, so this assay was used to validate enrichment of phage displaying covalently reactive Catcher.* ***(c)*** *Amino acid sequences of selected clones from the SpyCatcher003 (R32V-S59T-V94I) library selections. The final point mutations tested for the reconstitution reaction (rxn) were selected based on: (i) prioritizing residues which presented more than once within the fourth round which had cell lysate conditions, and (ii) mutations increasing hydrophobicity, since this approach proved successful in the generation of SpyCatcher003 (R32V-S59T-V94I).*

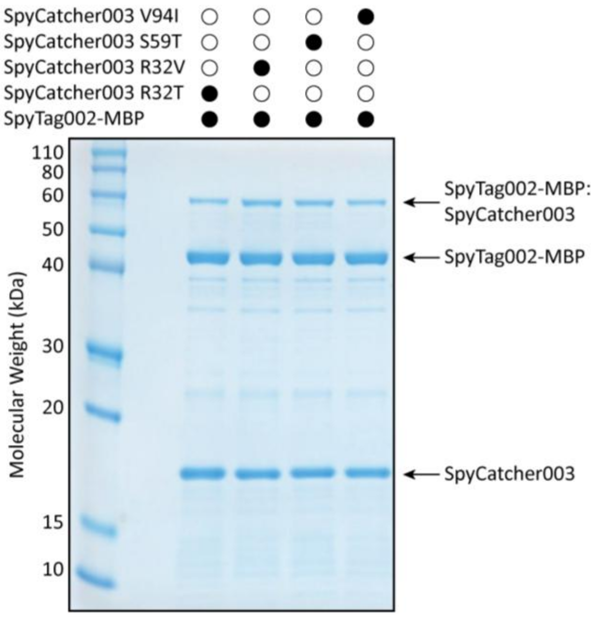


**Figure S4. Reactivity of the individual single-point mutants leading to R32V/S59T/V94I SpyCatcher003, which was used as a template for directed evolution.** *SpyTag002-MBP and SpyCatcher003 variants were each present at 1 µM. The reconstitution conditions were 15 minutes at 0 °C within quenched PBS sample buffer + 5 mM TCEP (pH 2.5). Representative SDS-PAGE with Coomassie staining from three independent measurements. Samples were only purified by IMAC before analysis.*

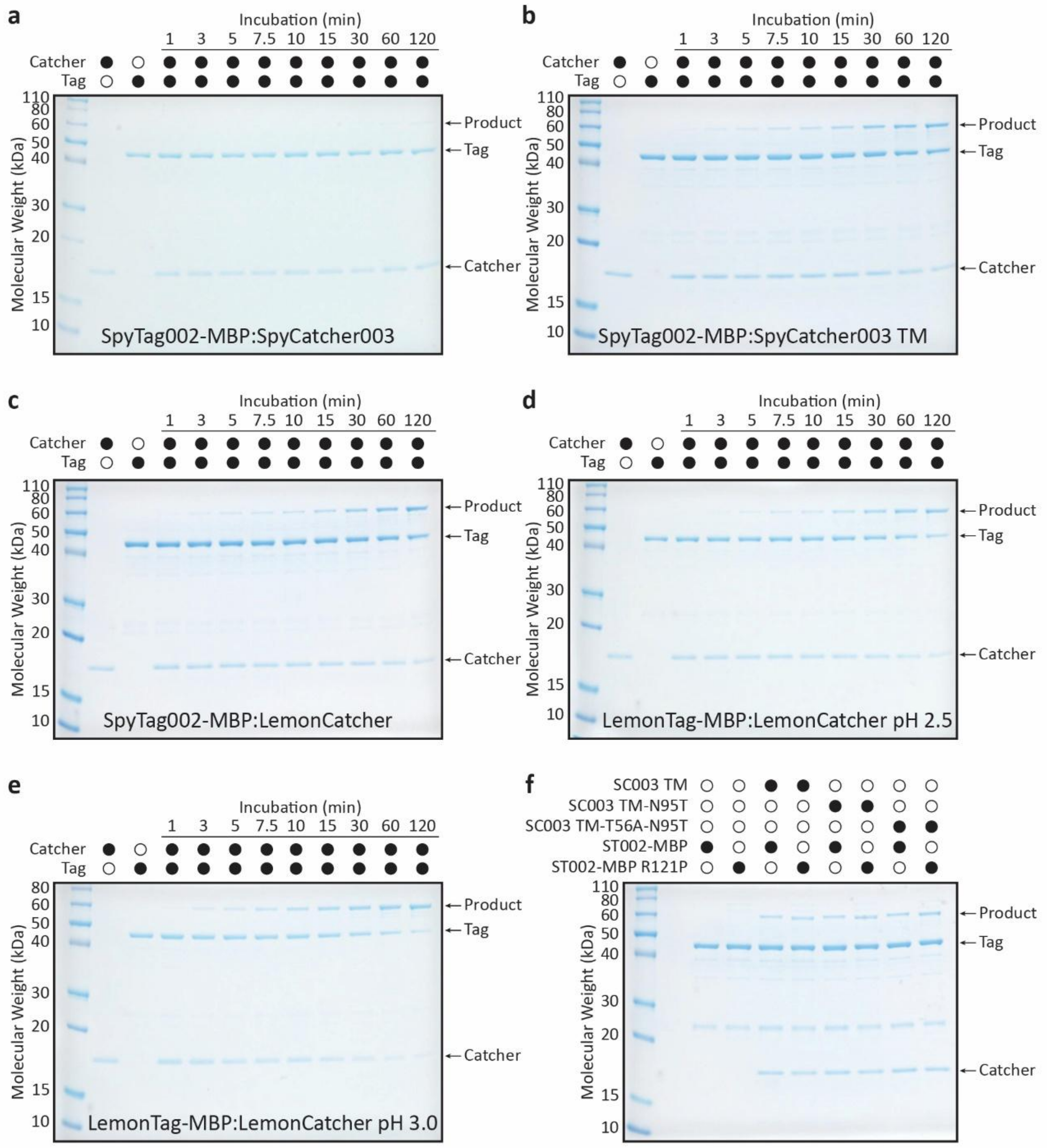


**Figure S5. Reaction rates of different Tag/Catcher pairs along the development pathway of the LemonTag/LemonCatcher pair.** *Representative SDS-PAGE gels with Coomassie staining for data presented in Figure 1 (**a**), Figure 3 (**b**), and Figure 5 (**c-e**). Both SpyTag-MBP and SpyCatcher003 variants were present at 1 µM and the reconstitution conditions were 15 minutes at 0 °C within quenched PBS sample buffer + 5 mM TCEP (pH 2.5) (**a-d** and **f**) or 3.0 (**e**). (**f**) Reactivity of the individual single-point mutants selected from phage display of the R32V/S59T/V94I (denoted here as triple mutant = TM) SpyCatcher003 construct used as a template for directed evolution. Tag and Catcher were each present at 1 µM and the reconstitution conditions were 15 minutes at 0 °C within quenched PBS sample buffer + 5 mM TCEP (pH 2.5). Representative gels of three independent measurements. SpyTag-MBP and SpyCatcher variants were purified by IMAC and size exclusion chromatography, except for in **Figure S2f** which was IMAC only.*

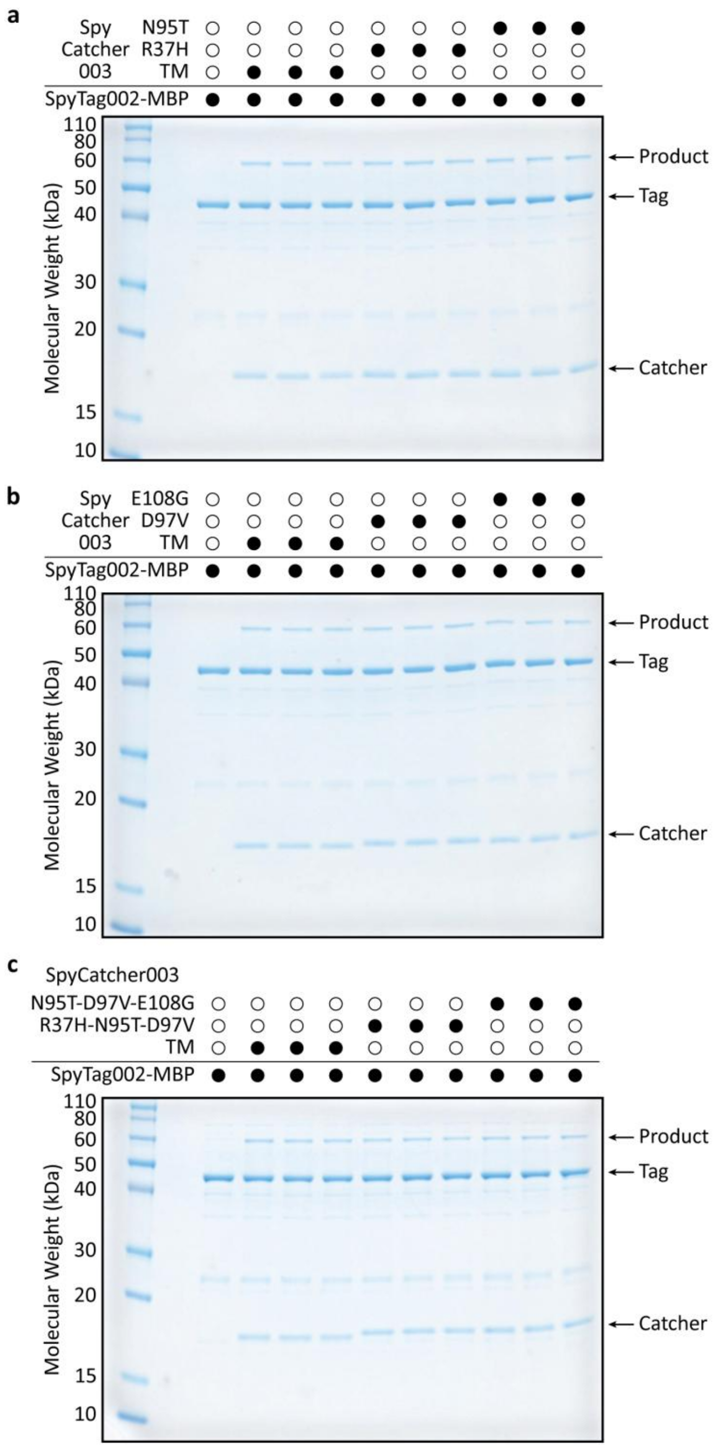


**Figure S6. Reactivity of the individual single-point mutants selected from phage display based on the R32V/S59T/V94I mutant.** *SpyTag002-MBP and SpyCatcher003 variants were each present at 1 µM. Reconstitution conditions were 15 minutes at 0 °C within quenched PBS sample buffer + 5 mM TCEP (pH 2.5), before SDS-PAGE with Coomassie staining. TM = triple mutant (R32V/S59T/V94I). SpyTag002-MBP and SpyCatcher003 variants were purified by IMAC and size exclusion chromatography.*

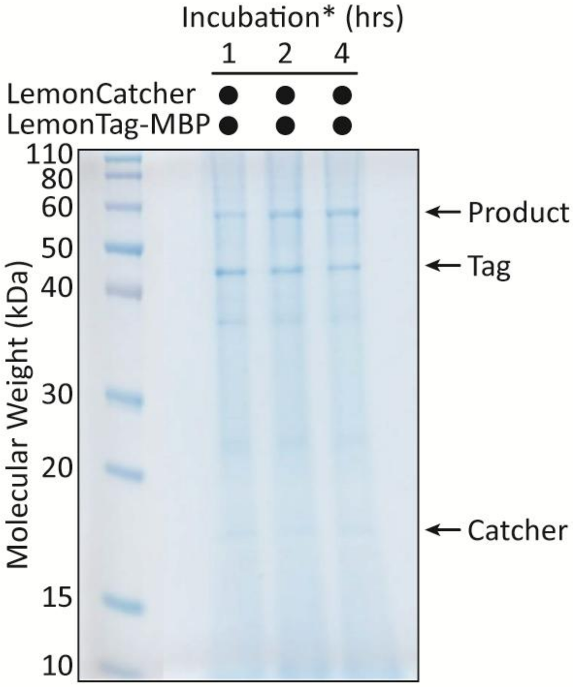


**Figure S7. LemonTag and LemonCatcher retain reactivity in lemon juice.** *Both Tag and Catcher were buffer exchanged into lemon juice (Packed in Belgium for Sainsbury's Supermarkets Ltd – Item code: 1180992) using a Micro Bio-Spin 6 column (Bio-Rad) before reaction at 1 µM each for the indicated time. Lemon juice was supplemented with 2.5 mM TCEP to prevent S49C mediated disulfide bond formation of Catcher dimers.*

```
                  1         10         20         30         40         50         60
                  |         |          |          |          |          |          |
SpyCatcher        VDTLSGLSSEQGQSGDMTIEEDSATHIKFSKRDEDGKELAGATMELRDSSGKTISTWISD
SpyCatcher002     VTTLSGLSGEQGPSGDMTTEEDSATHIKFSKRDEDGRELAGATMELRDSSGKTISTWISD
SpyCatcher003     VTTLSGLSGEQGPSGDMTTEEDSATHIKFSKRDEDGRELAGATMELRDSSGKTISTWISD
LemonCatcher      VTTLSGLSGEQGPSGDMTTEEDSATHIKFSKVDEDGRELAGATMELRDCSGKTISAWITD

                            70         80         90         100        110
                            |          |          |          |          |
SpyCatcher        GQVKDFYLYPGKYTFVETAAPDGYEVATAITFTVNEQGQVTVNGKATKGDAHI*
SpyCatcher002     GHVKDFYLYPGKYTFVETAAPDGYEVATAITFTVNEQGQVTVNGEATKGDAHT*
SpyCatcher003     GHVKDFYLYPGKYTFVETAAPDGYEVATPIEFTVNEDGQVTVDGEATEGDAHT*
LemonCatcher      GHVKDFYLYPGKYTFVETAAPDGYEVATPIEFTITEDGQVTVDGEATEGDAHT*

                     111       120
                     |         |
SpyTag            ---AHIVMVDAYKPTK
SpyTag002         --VPTIVMVDAYKRYK
SpyTag003         RGVPHIVMVDAYKRYK
LemonTag          --VPTIVMVDAYKPYK
```

Mutations arising from creation of SpyTag002/SpyCatcher002
Mutations arising from creation of SpyTag003/SpyCatcher003
Mutations arising from creation of LemonTag/LemonCatcher

**Figure S8: Amino acid sequence alignment of LemonTag/LemonCatcher with their preceding variants.** *Numbering is based on PDB 2X5P.*

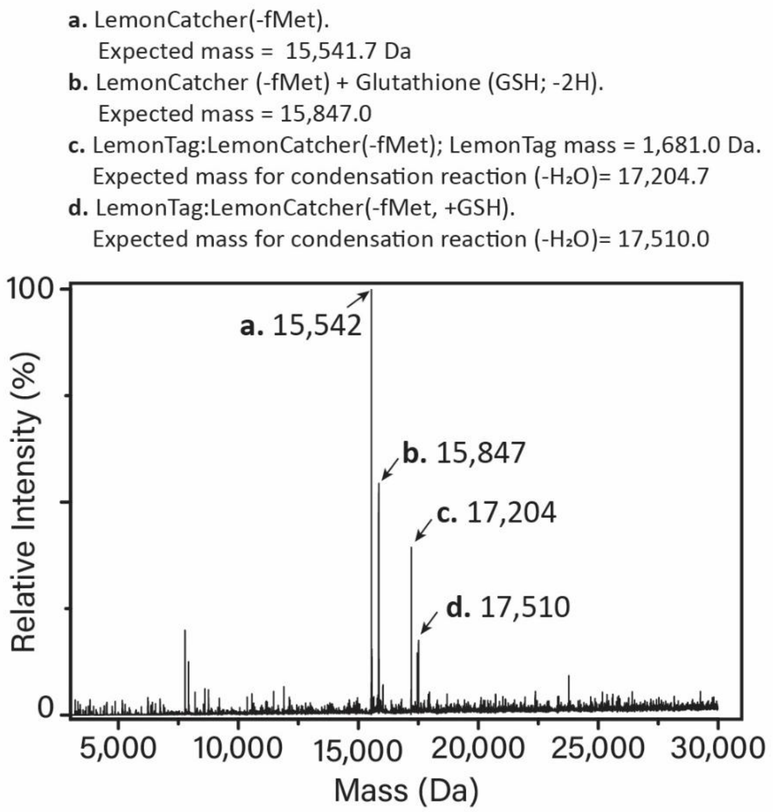


**Figure S9. Mass-transformed intact denatured mass spectrometry analysis of LemonTag:LemonCatcher complex.** *The starting fMet residue of the LemonCatcher constructs is removed efficiently in E. coli because the second amino acid is a serine amino acid which has a small sidechain[3]. Its expected mass = 15542 Da. The measured mass = 15,542 Da. There was a population which is modified through glutathionylation on the unique, free Cys49 residue – this was confirmed by TCEP treatment removing the modification and the mass difference (305 Da) matching the additional mass of glutathione (-2H; 307.3 – 2 = 305.3 Da). Although the glutathionylation is removed during our reaction tests (which include TCEP) and when coupled to beads via biotinlyation, we find that the modification does not alter the spontaneous amidation reactivity substantially.*

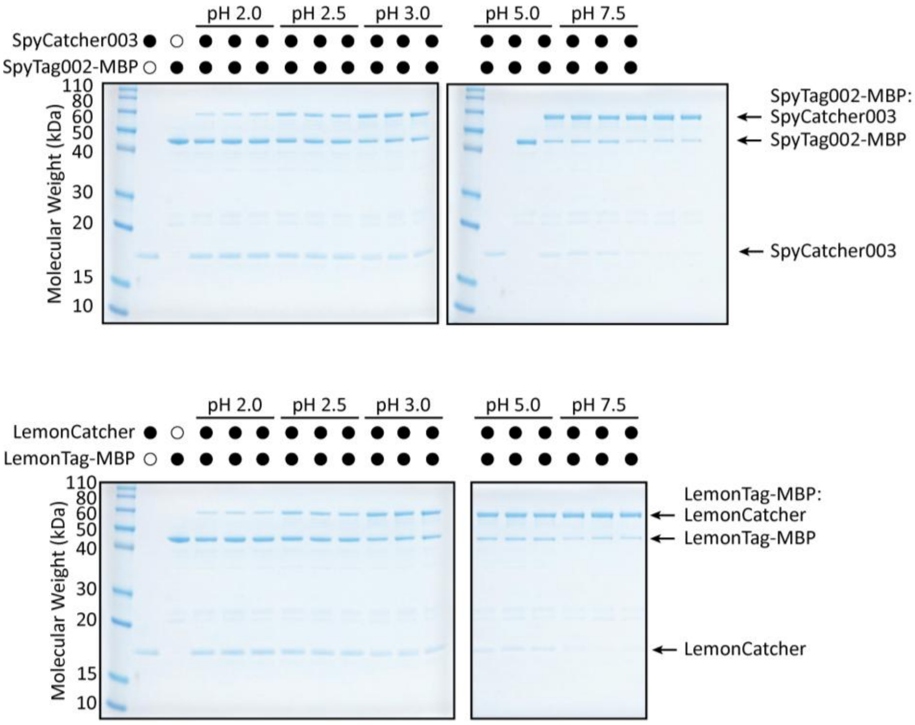


**Figure S10. Reactivity of SpyTag002:SpyCatcher003 and LemonTag:LemonCatcher at different pH values.** *(**Top**) SpyTag002-MBP and SpyCatcher003 were each present at 1 μM. Reconstitution conditions were 15 minutes at 0 °C in quenched or unquenched PBS sample buffer + 5 mM TCEP (pH as shown). (**Bottom**) LemonTag-MBP and LemonCatcher were each present at 1 μM. Reconstitution conditions were 15 minutes at 0 °C in quenched or unquenched PBS sample buffer + 5 mM TCEP (pH as shown). Three independent measurements are presented. Proteins were purified by IMAC and size exclusion chromatography.*

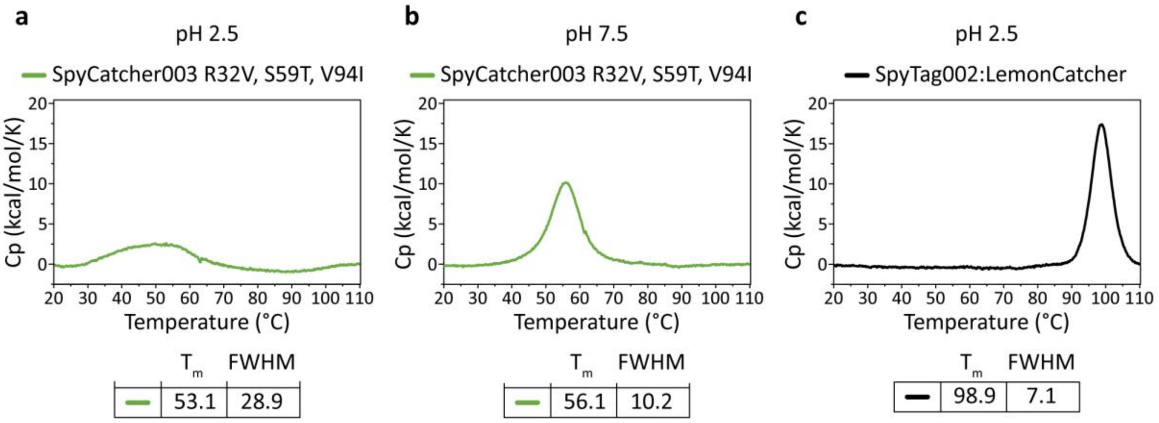


**Figure S11. DSC of Catcher variants at different pH values and reacted with Tag**. *DSC analysis of SpyCatcher003 R32V, S59T, V94I, in quenched or unquenched PBS sample buffer + 5 mM TCEP at pH 2.5 (**a**) or pH 7.5 (**b**) Below each graph, the melting temperature ($T_m$) and Full Width Half Maximum (FWHM) is tabulated. (**c**) DSC of preformed SpyTag002:LemonCatcher.*

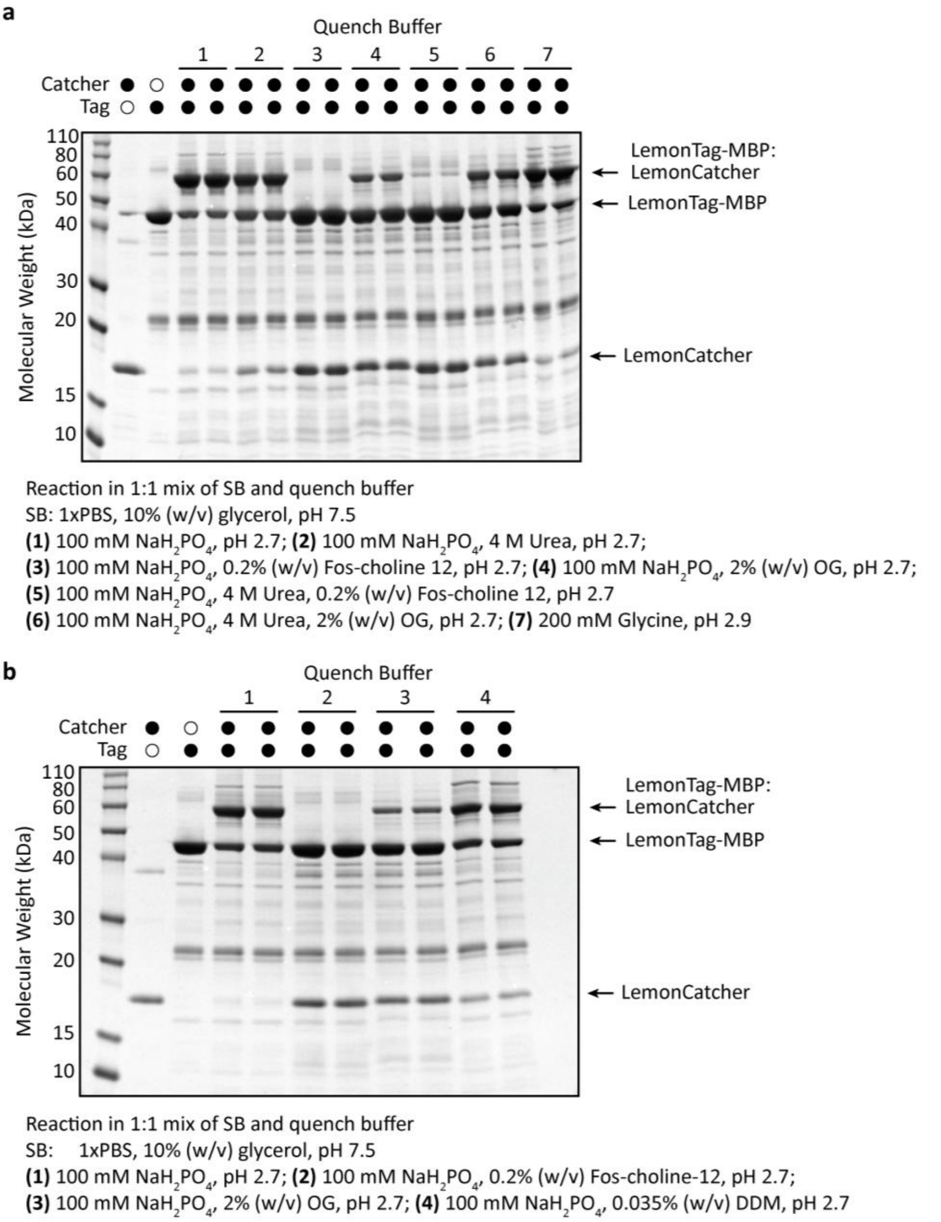


**Figure S12. LemonTag/LemonCatcher reactivity within different quench buffer conditions.** *LemonTag-MBP and LemonCatcher variants were each present at 10 µM and reacted at 0 °C, for 2 hours, in the different quenched PBS sample buffer (SB) mixtures indicated. Samples were only purified by IMAC before analysis.*

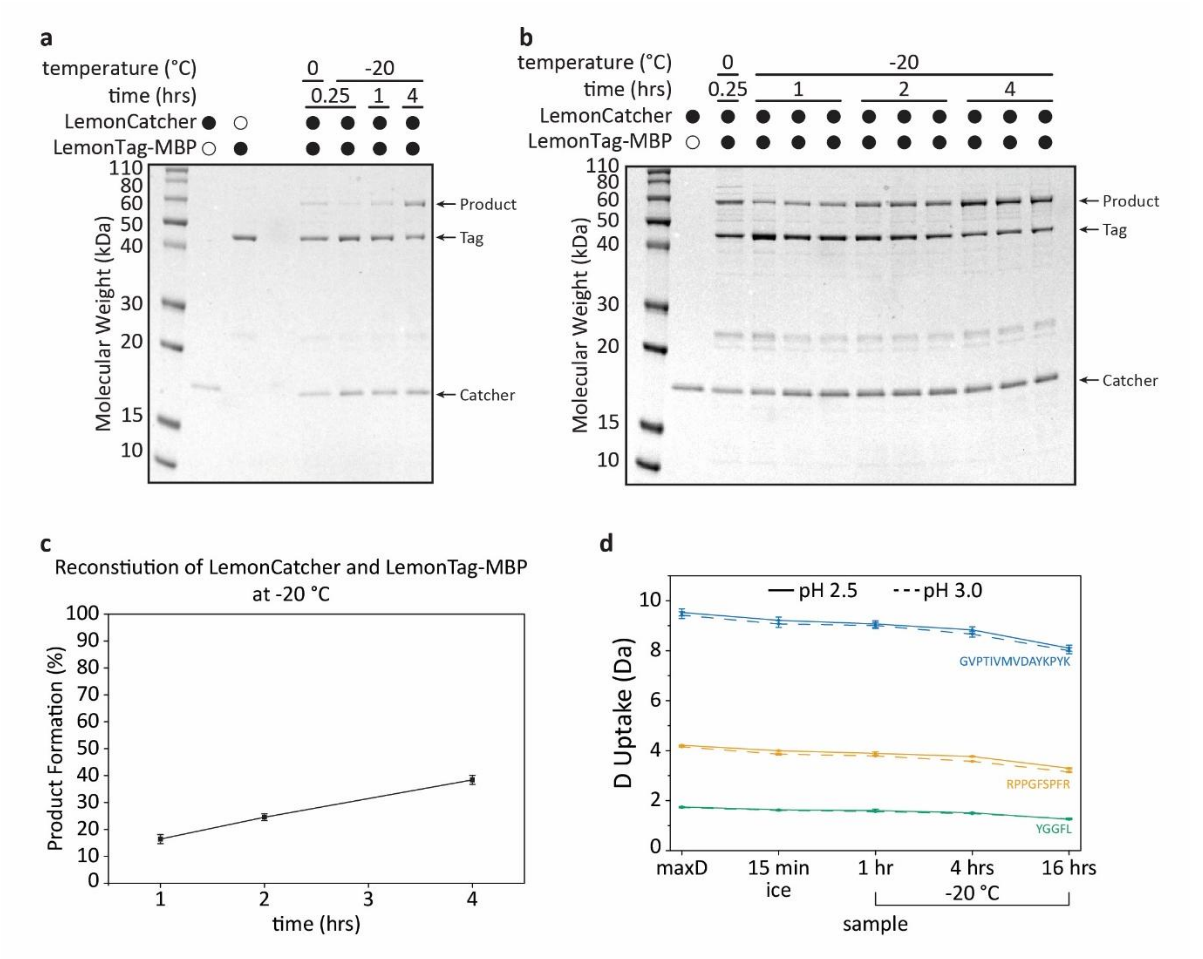


**Figure S13. Establishing LemonTag/LemonCatcher for covalent capture at -20 °C.** *LemonTag-MBP and LemonCatcher variants were each present at 1 µM and reacted at -20 °C, for the time points reported, in quenched PBS sample buffer, 40% (v/v) ethylene glycol + 5 mM TCEP at pH 2.5 (**a**) or pH 3.0 (**b**), before SDS-PAGE with Coomassie staining under reducing conditions. Addition of 40% (v/v) ethylene glycol prevents the solution from freezing at -20 °C and is used extensively in mass spectrometry because of its electrospray ionization compatibility[4]. Samples were only purified by IMAC before analysis. (**c**) Reaction speed at -20 °C, based on (b); data are mean ±1 s.d. (independent measurements n = 3). (**d**) Back exchange on model peptides within quenched PBS sample buffer at pH 2.5 (solid line) or pH 3.0 (dotted line) according to temperature and time. Data represent mean ±1 s.d. (independent measurements n = 3).*

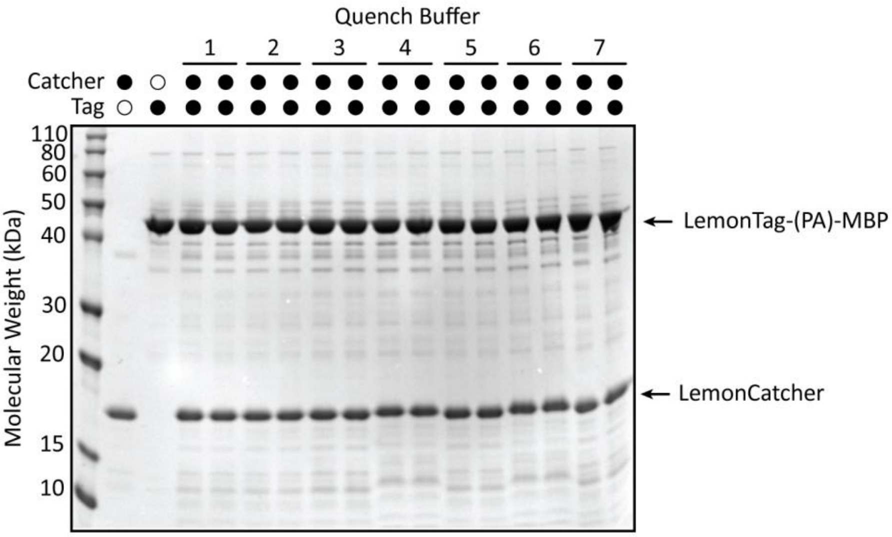


Reaction in 1:1 mix of SB and quench buffer
SB: PBS, 10% (w/v) glycerol, pH 7.5
**(1)** 100 mM $NaH_2PO_4$, pH 2.7; **(2)** 100 mM $NaH_2PO_4$, 4 M Urea, pH 2.7;
**(3)** 100 mM $NaH_2PO_4$, 0.2% (w/v) Fos-choline 12, pH 2.7; **(4)** 100 mM $NaH_2PO_4$, 2% (w/v) OG, pH 2.7;
**(5)** 100 mM $NaH_2PO_4$, 4 M Urea, 0.2% (w/v) Fos-choline 12, pH 2.7
**(6)** 100 mM $NaH_2PO_4$, 4 M Urea, 2% (w/v) OG, pH 2.7; **(7)** 200 mM Glycine, pH 2.9

**Figure S14. LemonTag(PA)/LemonCatcher did not react in any quench buffer conditions.** *LemonTag-MBP and LemonCatcher were each present at 10 µM and incubated together at 0 °C for 2 hours in the different quenched PBS sample buffer (SB) mixtures indicated. Samples were only purified by IMAC before analysis.*

**a**

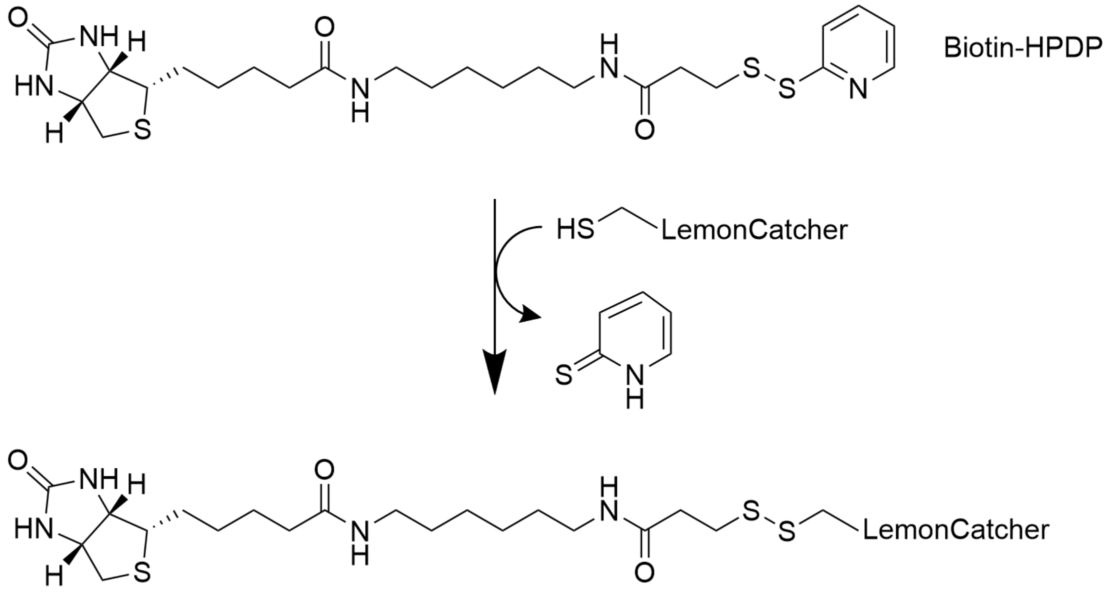


**b**

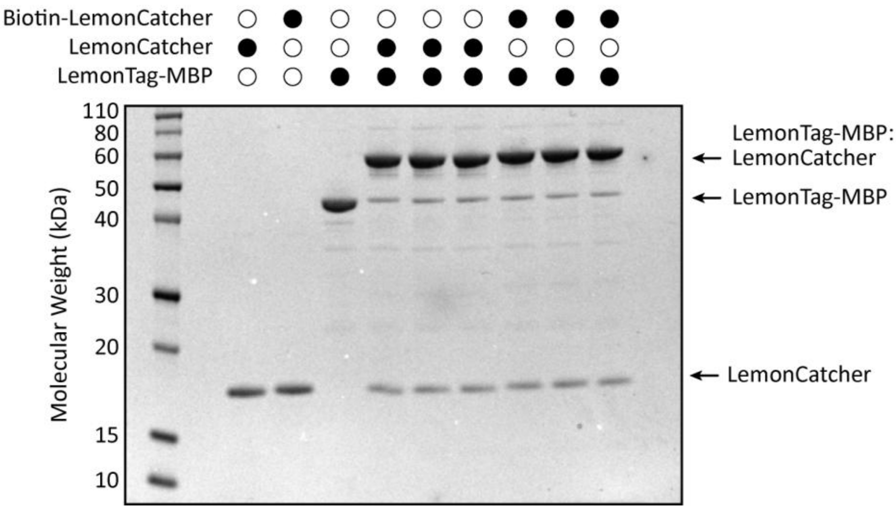


**Figure S15. LemonCatcher biotinylation and effect on reactivity.** ***(a)*** *Coupling of LemonCatcher to biotin-HPDP, to enable anchoring to streptavidin beads that can be rapidly released using reducing agent.* ***(b)*** *LemonTag reactivity was unaltered by biotinylating LemonCatcher. LemonTag-MBP and LemonCatcher ± biotinylation were each present at 10 µM and reacted at 0 °C, for 2 hours, in quenched PBS sample buffer, + 5 mM TCEP at pH 3.0. Proteins were purified by IMAC and size exclusion chromatography.*

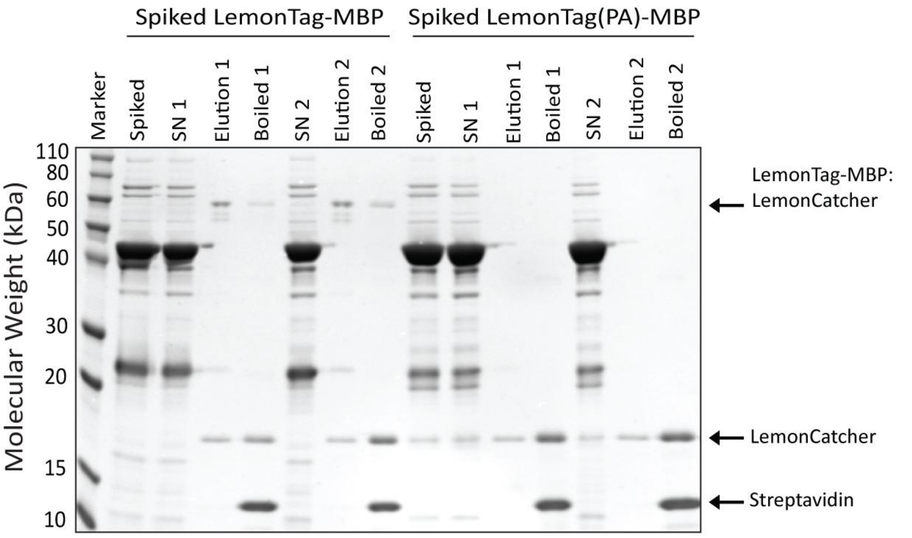


b

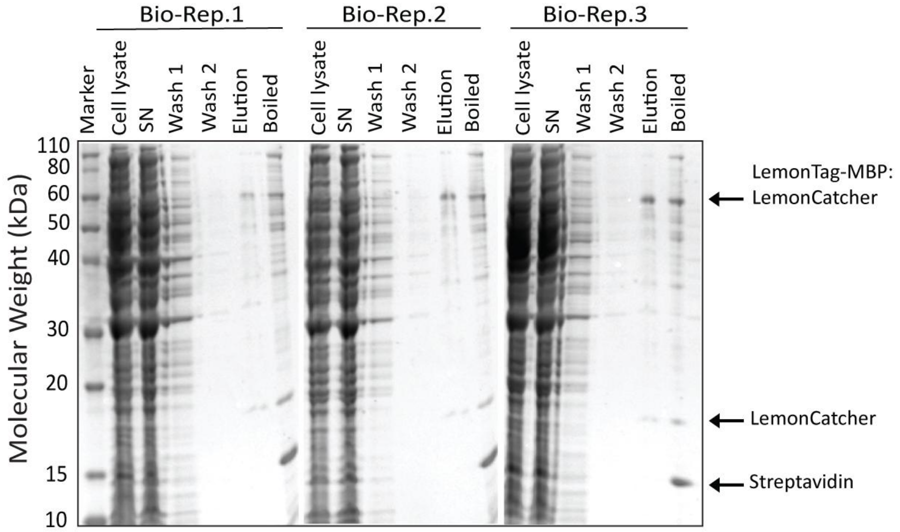


**Figure S16. SelQueX on purified LemonTag-MBP and replicate selective captures from cellular LemonTag-MBP.** *(**a**) Representative SDS-PAGE with Coomassie staining of quench-capture efficiency of POI (LemonTag-MBP) and LemonTag(PA)-MBP directly from cryo-milled cell lysate. Boiled samples lanes denote the remaining bound protein species after elution, removed after boiling in SDS loading buffer for 5 minutes (see **Methods**). (**b**) Reproducibility of the cellular SelQueX workflow related to **Fig. 3b**. Biological replicates (Bio-Rep.) 1-3 are shown.*

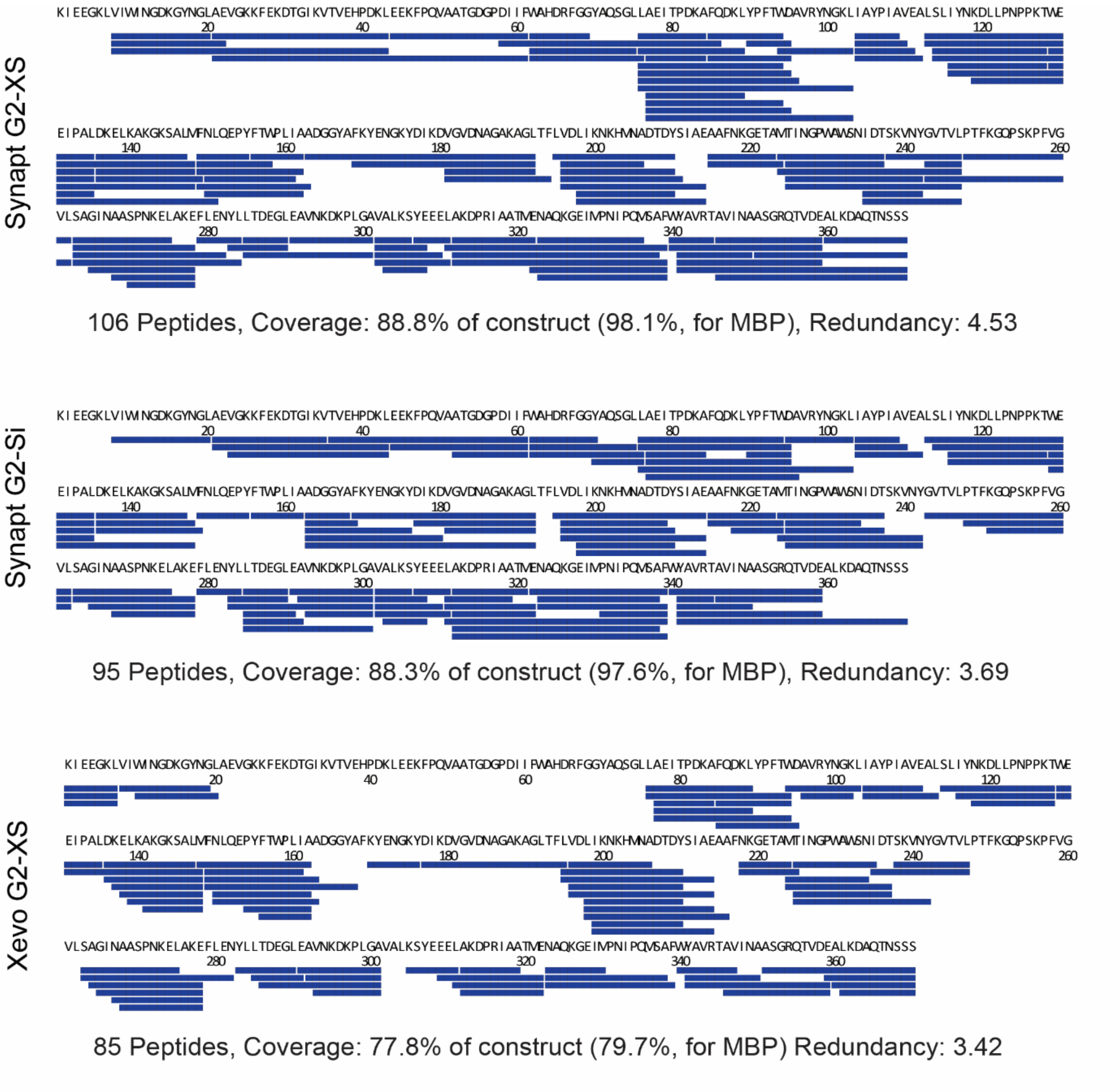


**Figure S17. Peptide coverage maps according to Mass Spectrometer used.** Peptide maps for cellular-contained LemonTagged MBP on Synapt G2-XS (Top), Synapt G2-Si (Middle) or Xevo G2-XS (Bottom) MS instruments. Only sequences corresponding to MBP are shown.

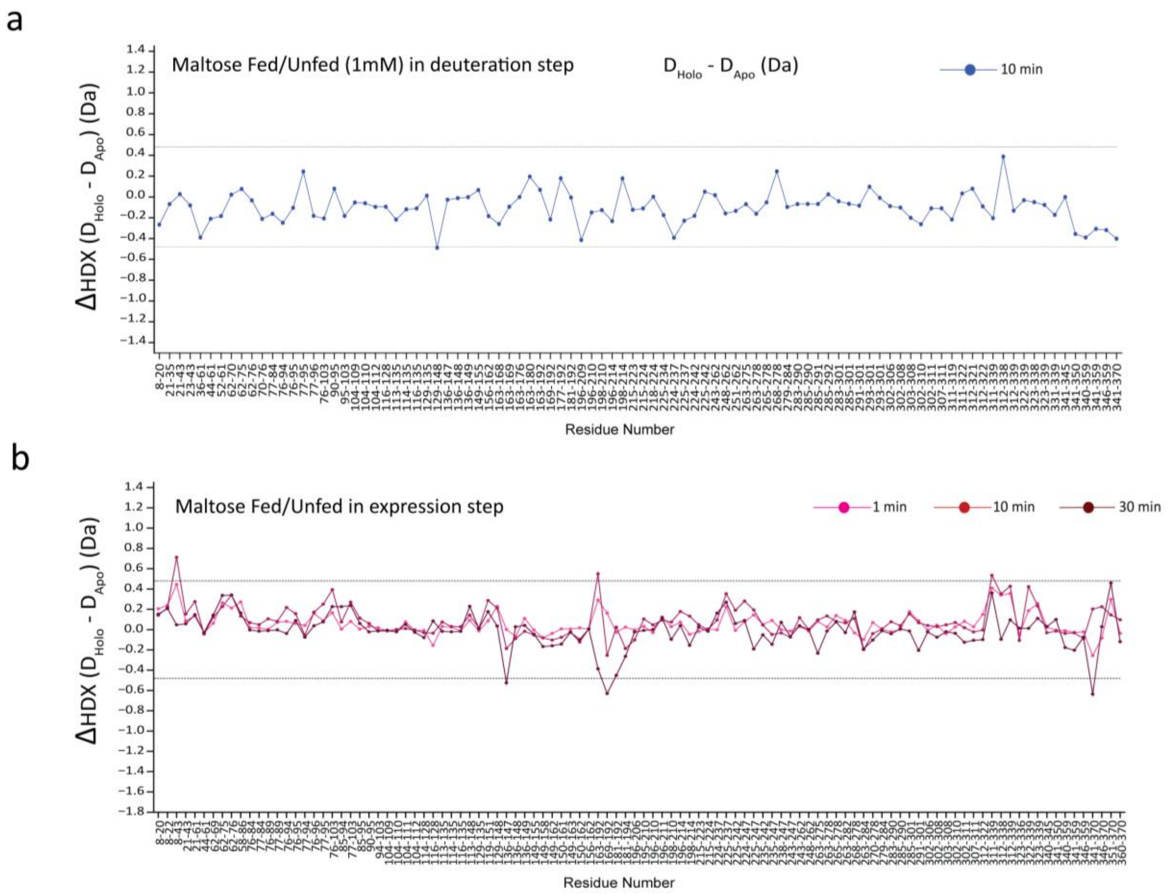


**Figure S18. Differential HDX-MS plots for (a) maltose un-fed, maltose during deuteration and (b) maltose-fed, maltose absent during deuteration.** *The dashed lines indicate the threshold of significance for ΔHDX ($p \leq 0.05$; ΔHDX significance cut-off of 0.40 Da; $n_{biological} = 3$). The MBP domains are illustrated in the top bars and residues comprising a region with significant differences in HDX are indicated by dotted lines. Peptides are arranged from the N- to C-terminus according to their peptide center residue.*

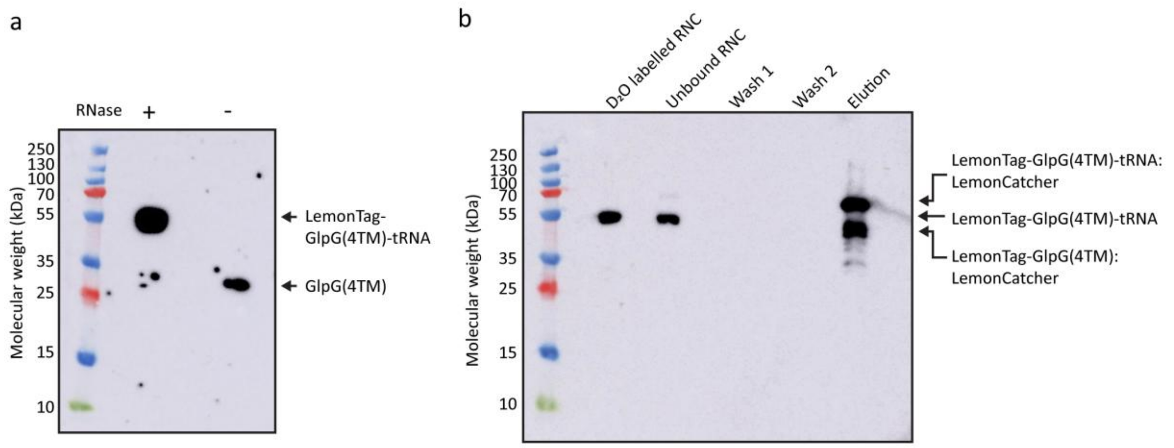


**Figure S19.** ***Western blot analysis on the GlpG(4TM) RNC sample preparation and during SelQuEX.*** *(**a**) Purified RNC GlpG(4TM) samples. The band at ~55 kDa represents GlpG-tRNA, and the lower band at ~30 kDa represents the GlpG nascent chain, with the tRNA having been digested by RNase A. (**b**) Western blot analysis on samples during the SelQueX workflow. We attribute the presence of tRNA-released product to spontaneous hydrolysis of peptidyl-tRNA ester bonds occurring during work-up. Samples were run on 12% (w/v) NuPAGE gels at neutral pH and with a sample dye at pH 5.7 [30% glycerol, 0.25 M Bis-Tris (pH 5.7), 0.8% DTT, 8% SDS, and bromophenol blue] to maintain the ester bond between the tRNA and the nascent chain. The samples were analyzed by Western blotting using anti-histidine-HRP (horse radish peroxidase) conjugate antibodies.*

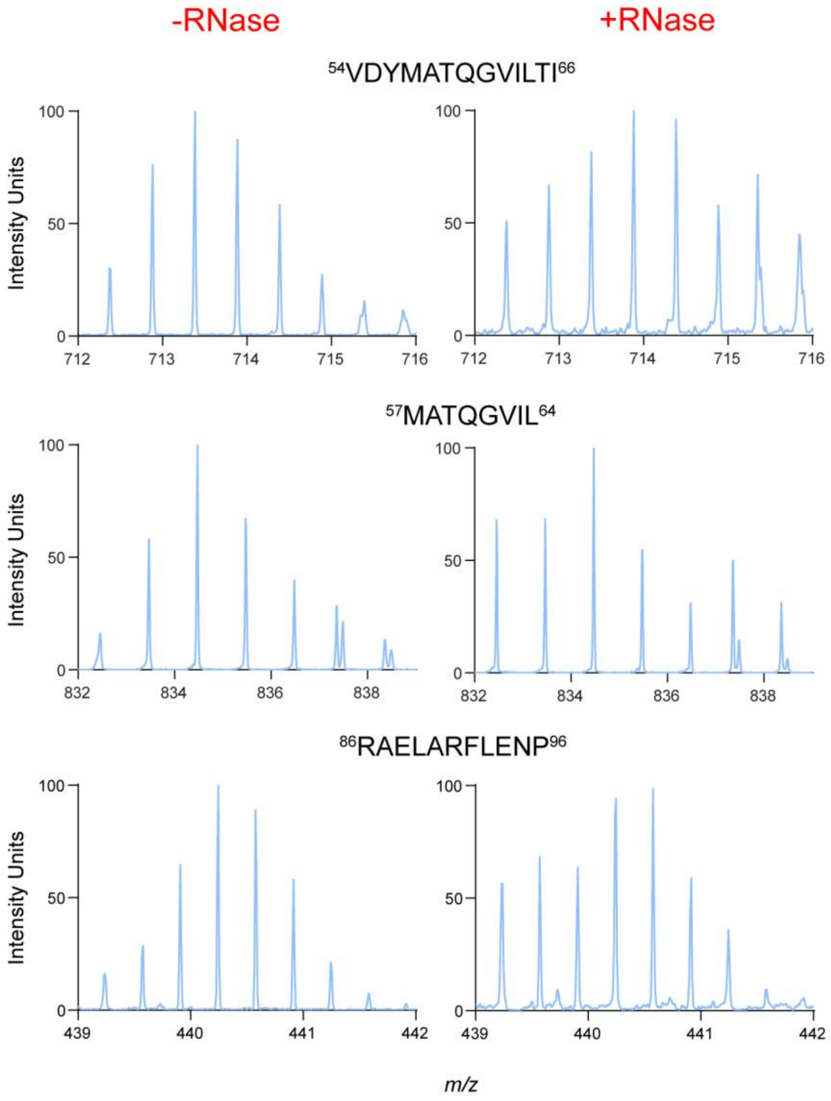


**Figure S20. Exemplar spectra of peptides in the CytD of GlpG.** *Sample that has been treated to RNase digestion show an increase in multimodal spectra, the lower spectral distributions can possibly be attributed to high exchange protection in aggregated populations.*

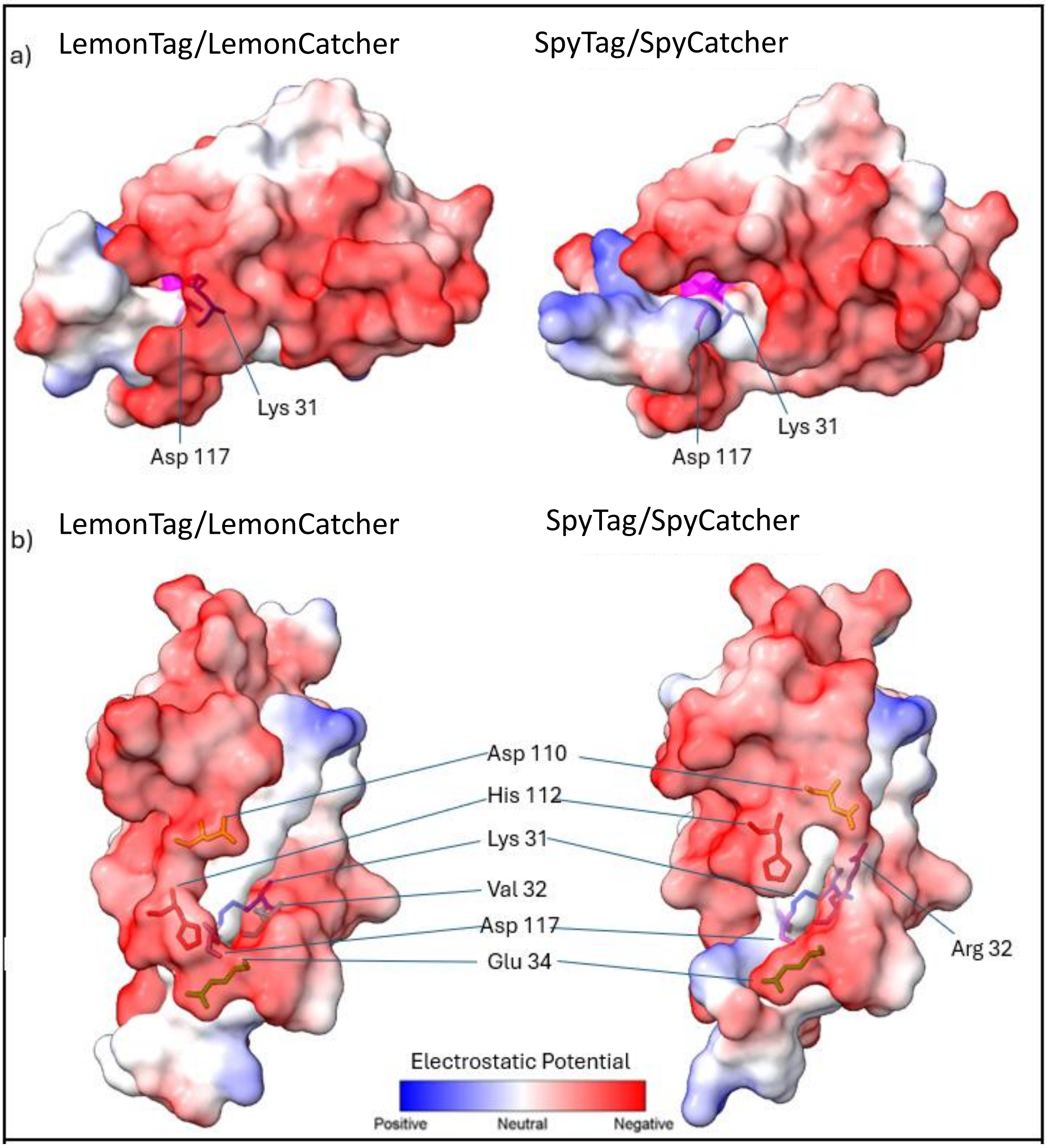


**Figure S21. Analysis of surface electrostatics of LemonTag/LemonCatcher.** *Top (**a**) and side (**b**) displays of electrostatic surface images of LemonTag:LemonCatcher and SpyTag/SpyCatcher. Red represents areas of high negative potential, with blue being positive electrostatic potential and white being neutral. Models were produced using AlphaFold 3*[5] *and visualized in ChimeraX*[6]*. The reactive residues Lys31 and Asp117 are marked, as well as residues with predicted orientation differences between SpyTag/SpyCatcher and LemonTag/LemonCatcher. Note that AlphaFold 3 assumes neutral pH.*

**Primers for SpyCatcher003 mutations (5' to 3')**

| Mutation | Forward primer | Reverse primer |
|---|---|---|
| **H26N** | TAGTGCTACCAACATTAAATTCTCAAAAC | TCTTCTTCAGTTGTCATATC |
| **K31A** | TAAATTCTCAGCGCGTGATGAGGACG | ATATGGGTAGCACTATCTTC |
| **K28I** | TACCCATATTATTTTCTCAAAACGTGATG | GCACTATCTTCTTCAGTTG |
| **R32T** | ATTCTCAAAAACCGATGAGGACGGCC | TTAATATGGGTAGCACTATC |
| **R32V** | ATTCTCAAAAGTGGATGAGGACGGCCG | TTAATATGGGTAGCACTATC |
| **R37N** | TGAGGACGGCAACGAGTTAGCTG | TCACGTTTTGAGAATTTAATATG |
| **R37H** | TGAGGACGGCCATGAGTTAGCTG | TCCACTTTTGAGAATTTAATATGGGTAGC |
| **R47T** | TATGGAGTTGACCGATTGCTCTGGTAAAAC | GTTGCACCAGCTAACTCAC |
| **K52E** | TTGCTCTGGTGAAACTATTAGTAC | TCACGCAACTCCATAGTTG |
| **T56A** | AACTATTAGTGCATGGATTACCG | TTACCAGAGCAATCACGC |
| **S59T** | TACATGGATTACCGATGGACATGTG | CTAATAGTTTTACCAGAGC |
| **H62Q** | TTCAGATGGACAGGTGAAGGATTTC | ATCCATGTACTAATAGTTTTACC |
| **K72E** | GTATCCAGGAGAATATACATTTGTC | AGGTAGAAATCCTTCACATG |
| **V76I** | ATATACATTTATTGAAACCGCAGC | TTTCCTGGATACAGGTAG |
| **V94I** | TGAATTTACAATTAATGAGGACG | ATTGGAGTTGCTACCTCATAAC |
| **N95T** | ATTTACAATTACCGAGGACGGTCAGGTTAC | TCAATTGGAGTTGCTACC |
| **D97V** | AATTAATGAGGTGGGTCAGGTTACTG | GTAAATTCAATTGGAGTTGC |
| **E108G** | TGAAGCAACTGGCGGTGACGCTC | CCATCTACAGTAACCTGAC |
| **H112Q** | AGGTGACGCTCAGACTGGATCCA | TCAGTTGCTTCACCATCTACAG |

**Primers for SpyTag002 mutations (5' to 3')**

| Mutation | Forward primer | Reverse primer |
|---|---|---|
| **KRYK deletion** | GGTAGTGGTGAAAGTGGTAAAATC | GTAGGCGTCCACCATCAC |
| **K120E** | GGACGCCTACGAACGTTACAAGG | ACCATCACGATAGTAGGC |
| **R121T** | CGCCTACAAGACCTACAAGGGTAG | TCCACCATCACGATAGTAG |
| **K120E/R121T** | GGACGCCTACGAAACCTACAAGGGTAGTGG | ACCATCACGATAGTAGGC |
| **R121P** | CGCCTACAAGCCGTACAAGGGTAG | TCCACCATCACGATAGTAG |

**Table S1. Overview of mutations in SpyCatcher003 and SpyTag002, including the corresponding forward and reverse primer for site-directed mutagenesis.** *Mutations in green improved reactivity, mutations in orange decreased reactivity, and mutations in white led to no measurable difference in reactivity. Red highlights the K31A mutant which prevents the spontaneous amidation reaction and acts as a negative control.*

| | **Efficiency of Binding to Tosylactivated Magnetic LemonCatcher DynaBeads** | | | | | |
|---|---|---|---|---|---|---|
| | PBS Quench Conditions | | Glycine Quench Conditions | | Glycine Quench Conditions using Glycine-Blocked Beads | |
| *Volume Bead Slurry (μL)* | *Reactive LemonTag* | *Non-Reactive (PA) LemonTag* | *Reactive LemonTag* | *Non-Reactive (PA) LemonTag* | *Reactive LemonTag* | *Non-Reactive (PA) LemonTag* |
| 10 | 36% | 38% | 23% | 26% | 22% | 26% |
| 5 | 22% | 17% | 8% | 15% | 10% | 11% |
| 2 | 10% | 5% | 8% | 11% | 3% | 5% |
| 0 | 0% | 0% | 0% | 0% | 0% | 0% |

**Table S2. Optimization of capture using LemonCatcher beads.** *ImageJ analysis was conducted on Coomassie-stained SDS-PAGE from three independent quench capture experiments. LemonTag capture by LemonCatcher beads was conducted for 15 min at 4 °C under quench conditions. LemonTag-MBP and beads were pre-quenched before conducting the capture reaction experiments.*

| LemonTag-MBP protein (Dataset) | | | | |
|---|---|---|---|---|
| **Data Set** | **Apo-LemonTag-MBP** | **Holo-LemonTag-MBP Maltose-bound protein** | **Apo-LemonTag-MBP and Cryo-Milled Method for SelQueX back exchange calculation** | **Apo LemonTag-MBP Standard Method** |
| **HDX reaction details** | 800 µL LB/$D_2O$ buffer containing 2.5 g/L LB, NaPi 50 mM and 150 mM NaCl, pD 7.0-7.2 ($pH_{read}$= 6.6-6.8) was added to 200 µL cell lysate | | 800 µL LB/$D_2O$ buffer containing 2.5 g/L LB, NaPi 50 mM and 150 mM NaCl, pD 7.0-7.2 ($pH_{read}$ = 6.6-6.8) was added to 200 µL purified LemonTag-MBP | 800 µL $D_2O$ buffer containing NaPi 50 mM and 150 mM NaCl, pD 7.0-7.2 ($pH_{read}$ = 6.6-6.8) was added to 200 µL purified LemonTag-MBP |
| **Temperature** | 22 °C | | | |
| **HDX time course (min), in 80% Deuterium** | 1, 10 and 30 | | 1,440 | |
| **HDX control samples** | purified protein (Dmax): maximum D-uptake (with 24 h labeling time); non-deuterated | | | |
| **Back exchange (mean)*** | 47.7% ± 9.7 calculated from $t_{HDX}$ = 30 min of peptides covering residues 340+ at the flexible C-termini[7] | | | 32.1± 2.0 % calculated from $t_{HDX}$ = 30 min of peptides covering residues 340+ at the flexible C-termini[7] |
| **# of Peptides** | 103 | | | |
| **Sequence coverage** | 89% LemonTag-MBP construct (98% for MBP) | | | |
| **Average peptide length / Redundancy** | 13.5 / 4.40 | | | |
| **Protease** | Pepsin | | | |
| **Replicates (biological or technical)** | 3 biological replicates | | | |
| **Repeatability (average standard deviation)** | 0.13 | 0.16 | 0.20 | 0.07 |

**Table S3. HDX-MS reporting table for LemonTag-MBP experiments.** *Back exchange calculated as the average for all peptides measured with error bars indicating the interquartile range of these values

### Suppelmentary information references


1. Li, L., Fierer, J. O., Rapoport, T. A. & Howarth, M. Structural Analysis and Optimization of the Covalent Association between SpyCatcher and a Peptide Tag. *J. Mol. Biol.* **426**, 309–317 (2014).
2. Suderman, R. J., Rice, D. A., Gibson, S. D., Strick, E. J. & Chao, D. M. Development of polyol-responsive antibody mimetics for single-step protein purification. *Protein Expr. Purif.* **134**, 114–124 (2017).
3. Wingfield, P. T. N-Terminal Methionine Processing. *Curr. Protoc. Protein Sci.* **88**, 6.14.1-6.14.3 (2017).
4. Venable, J. D., Okach, L., Agarwalla, S. & Brock, A. Subzero Temperature Chromatography for Reduced Back-Exchange and Improved Dynamic Range in Amide Hydrogen/Deuterium Exchange Mass Spectrometry. *Anal. Chem.* **84**, 9601–9608 (2012).
5. Abramson, J. *et al.* Accurate structure prediction of biomolecular interactions with AlphaFold 3. *Nature* **630**, 493–500 (2024).
6. Pettersen, E. F. *et al.* UCSF Chimera—A visualization system for exploratory research and analysis. *J. Comput. Chem.* **25**, 1605–1612 (2004).
7. Evenäs, J. *et al.* Ligand-induced structural changes to maltodextrin-binding protein as studied by solution NMR spectroscopy11Edited by P. E. Wright. *J. Mol. Biol.* **309**, 961–974 (2001).
