## Supplementary figures and images for "LemonCatcher Acidic Pull-Down Enables Selective In-Cell Hydrogen-Deuterium Exchange Mass Spectrometry"

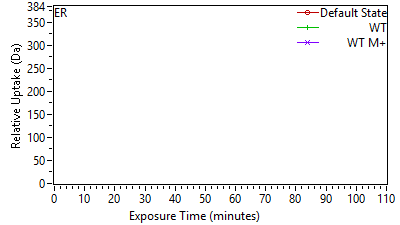

Supplement: Supporting Data File 1 [file 747387_file03.zip › uptake plots_cond 1_112025/01 ER[2-410] MGSSHHHHHHSSGLVP.png]

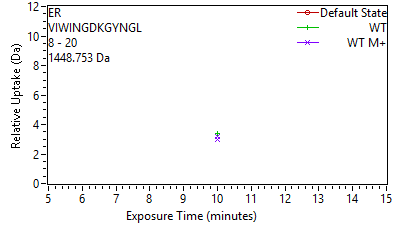

Supplement: Supporting Data File 1 [file 747387_file03.zip › uptake plots_cond 1_112025/02 ER[47-59] VIWINGDKGYNGL.png]

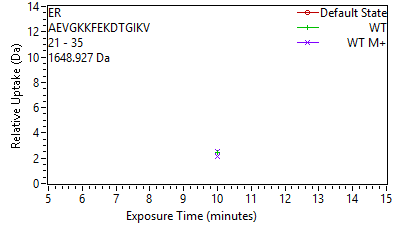

Supplement: Supporting Data File 1 [file 747387_file03.zip › uptake plots_cond 1_112025/03 ER[60-74] AEVGKKFEKDTGIKV.png]

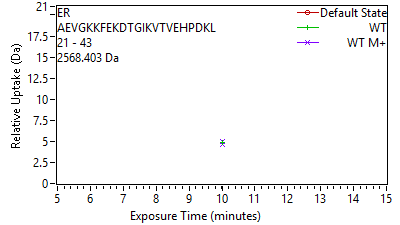

Supplement: Supporting Data File 1 [file 747387_file03.zip › uptake plots_cond 1_112025/04 ER[60-82] AEVGKKFEKDTGIKVT.png]

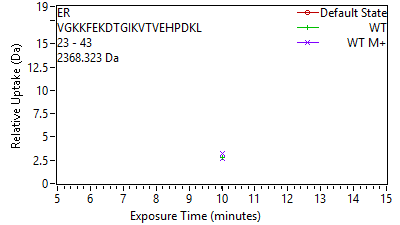

Supplement: Supporting Data File 1 [file 747387_file03.zip › uptake plots_cond 1_112025/05 ER[62-82] VGKKFEKDTGIKVTVE.png]

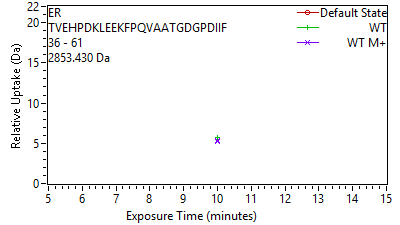

Supplement: Supporting Data File 1 [file 747387_file03.zip › uptake plots_cond 1_112025/06 ER[75-100] TVEHPDKLEEKFPQVA.png]

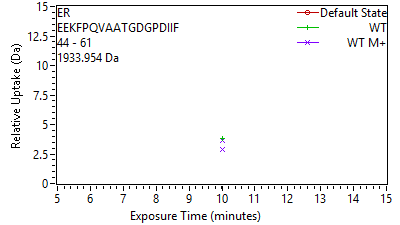

Supplement: Supporting Data File 1 [file 747387_file03.zip › uptake plots_cond 1_112025/07 ER[83-100] EEKFPQVAATGDGPDI.png]

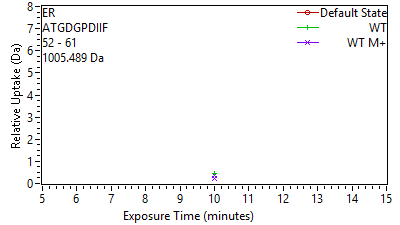

Supplement: Supporting Data File 1 [file 747387_file03.zip › uptake plots_cond 1_112025/08 ER[91-100] ATGDGPDIIF.png]

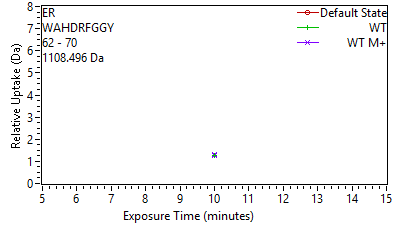

Supplement: Supporting Data File 1 [file 747387_file03.zip › uptake plots_cond 1_112025/09 ER[101-109] WAHDRFGGY.png]

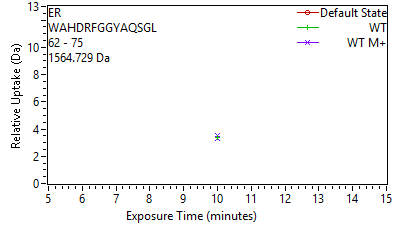

Supplement: Supporting Data File 1 [file 747387_file03.zip › uptake plots_cond 1_112025/10 ER[101-114] WAHDRFGGYAQSGL.png]

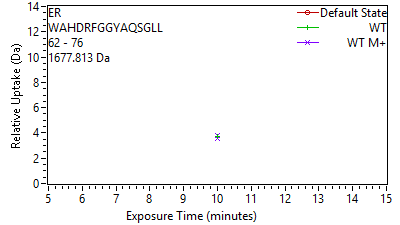

Supplement: Supporting Data File 1 [file 747387_file03.zip › uptake plots_cond 1_112025/11 ER[101-115] WAHDRFGGYAQSGLL.png]

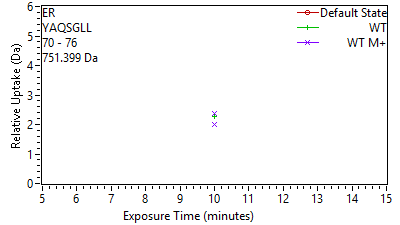

Supplement: Supporting Data File 1 [file 747387_file03.zip › uptake plots_cond 1_112025/12 ER[109-115] YAQSGLL.png]

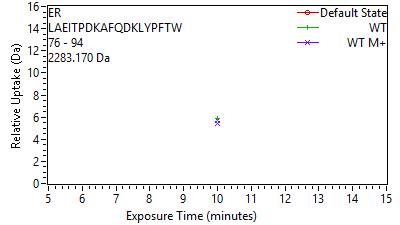

Supplement: Supporting Data File 1 [file 747387_file03.zip › uptake plots_cond 1_112025/13 ER[115-133] LAEITPDKAFQDKLYP.png]

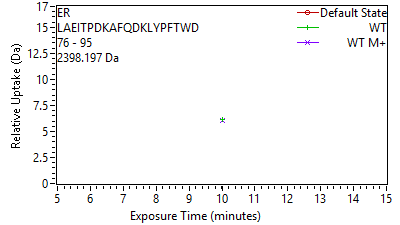

Supplement: Supporting Data File 1 [file 747387_file03.zip › uptake plots_cond 1_112025/14 ER[115-134] LAEITPDKAFQDKLYP.png]

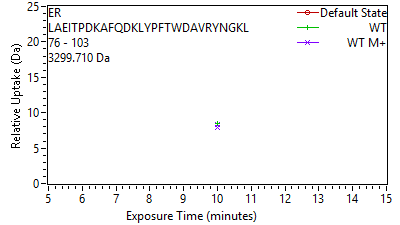

Supplement: Supporting Data File 1 [file 747387_file03.zip › uptake plots_cond 1_112025/15 ER[115-142] LAEITPDKAFQDKLYP.png]

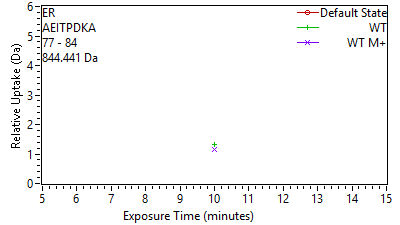

Supplement: Supporting Data File 1 [file 747387_file03.zip › uptake plots_cond 1_112025/16 ER[116-123] AEITPDKA.png]

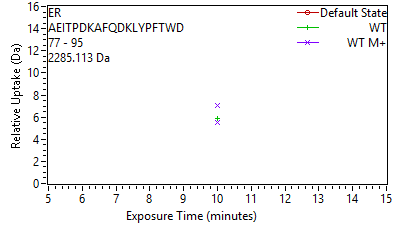

Supplement: Supporting Data File 1 [file 747387_file03.zip › uptake plots_cond 1_112025/17 ER[116-134] AEITPDKAFQDKLYPF.png]

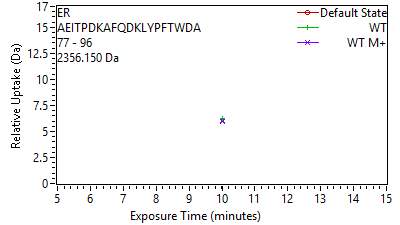

Supplement: Supporting Data File 1 [file 747387_file03.zip › uptake plots_cond 1_112025/18 ER[116-135] AEITPDKAFQDKLYPF.png]

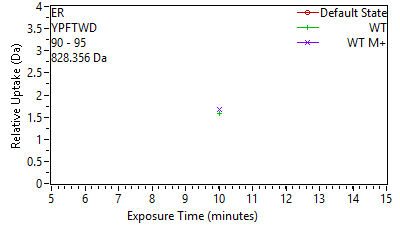

Supplement: Supporting Data File 1 [file 747387_file03.zip › uptake plots_cond 1_112025/19 ER[129-134] YPFTWD.png]

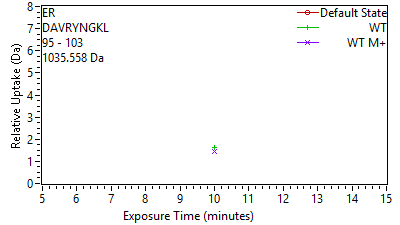

Supplement: Supporting Data File 1 [file 747387_file03.zip › uptake plots_cond 1_112025/20 ER[134-142] DAVRYNGKL.png]

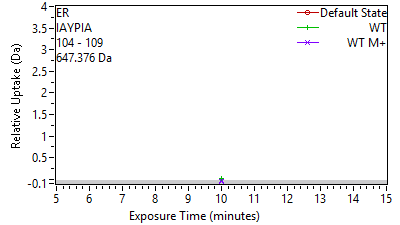

Supplement: Supporting Data File 1 [file 747387_file03.zip › uptake plots_cond 1_112025/21 ER[143-148] IAYPIA.png]

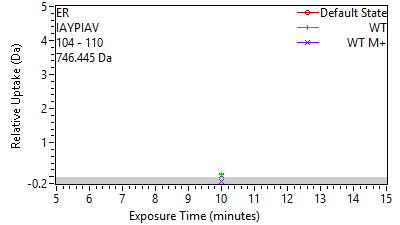

Supplement: Supporting Data File 1 [file 747387_file03.zip › uptake plots_cond 1_112025/22 ER[143-149] IAYPIAV.png]

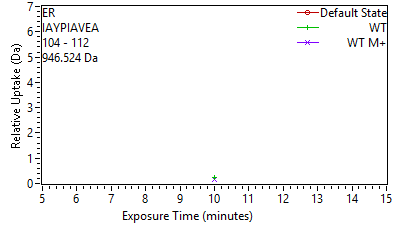

Supplement: Supporting Data File 1 [file 747387_file03.zip › uptake plots_cond 1_112025/23 ER[143-151] IAYPIAVEA.png]

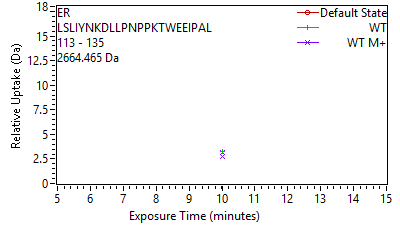

Supplement: Supporting Data File 1 [file 747387_file03.zip › uptake plots_cond 1_112025/24 ER[152-174] LSLIYNKDLLPNPPKT.png]

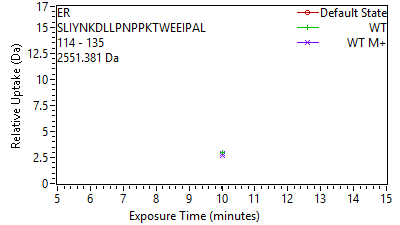

Supplement: Supporting Data File 1 [file 747387_file03.zip › uptake plots_cond 1_112025/25 ER[153-174] SLIYNKDLLPNPPKTW.png]

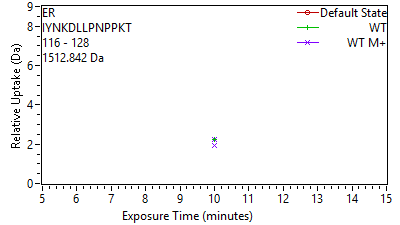

Supplement: Supporting Data File 1 [file 747387_file03.zip › uptake plots_cond 1_112025/26 ER[155-167] IYNKDLLPNPPKT.png]

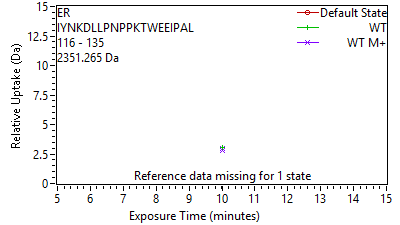

Supplement: Supporting Data File 1 [file 747387_file03.zip › uptake plots_cond 1_112025/27 ER[155-174] IYNKDLLPNPPKTWEE.png]

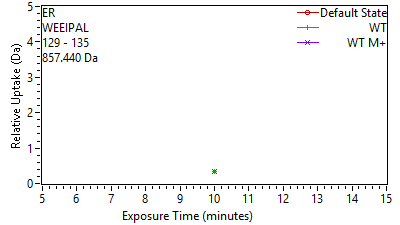

Supplement: Supporting Data File 1 [file 747387_file03.zip › uptake plots_cond 1_112025/28 ER[168-174] WEEIPAL.png]

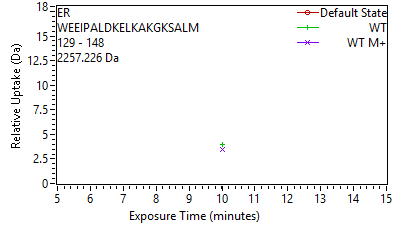

Supplement: Supporting Data File 1 [file 747387_file03.zip › uptake plots_cond 1_112025/29 ER[168-187] WEEIPALDKELKAKGK.png]

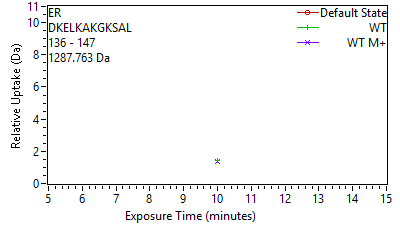

Supplement: Supporting Data File 1 [file 747387_file03.zip › uptake plots_cond 1_112025/30 ER[175-186] DKELKAKGKSAL.png]

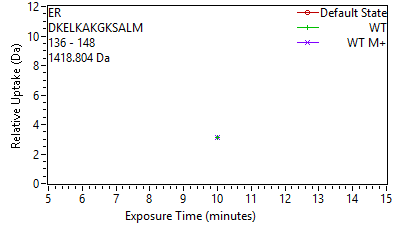

Supplement: Supporting Data File 1 [file 747387_file03.zip › uptake plots_cond 1_112025/31 ER[175-187] DKELKAKGKSALM.png]

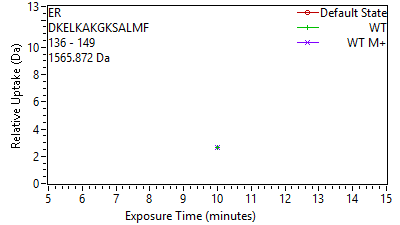

Supplement: Supporting Data File 1 [file 747387_file03.zip › uptake plots_cond 1_112025/32 ER[175-188] DKELKAKGKSALMF.png]

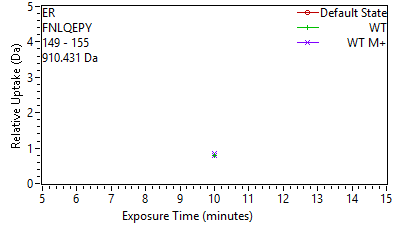

Supplement: Supporting Data File 1 [file 747387_file03.zip › uptake plots_cond 1_112025/33 ER[188-194] FNLQEPY.png]

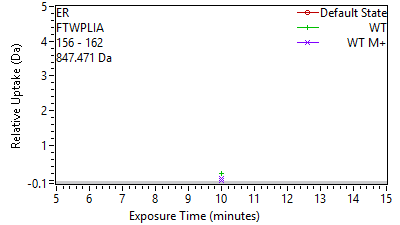

Supplement: Supporting Data File 1 [file 747387_file03.zip › uptake plots_cond 1_112025/34 ER[195-201] FTWPLIA.png]

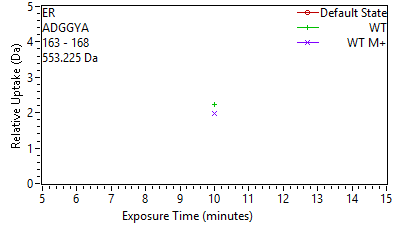

Supplement: Supporting Data File 1 [file 747387_file03.zip › uptake plots_cond 1_112025/35 ER[202-207] ADGGYA.png]

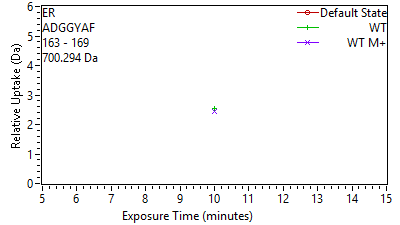

Supplement: Supporting Data File 1 [file 747387_file03.zip › uptake plots_cond 1_112025/36 ER[202-208] ADGGYAF.png]

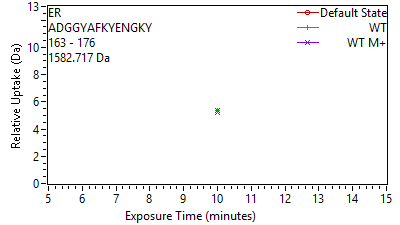

Supplement: Supporting Data File 1 [file 747387_file03.zip › uptake plots_cond 1_112025/37 ER[202-215] ADGGYAFKYENGKY.png]

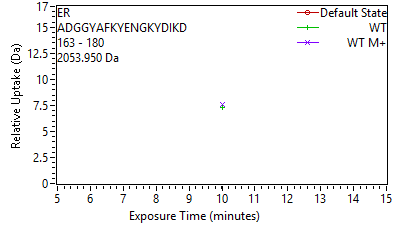

Supplement: Supporting Data File 1 [file 747387_file03.zip › uptake plots_cond 1_112025/38 ER[202-219] ADGGYAFKYENGKYDI.png]

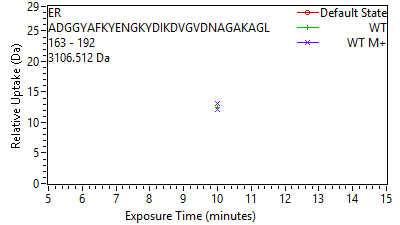

Supplement: Supporting Data File 1 [file 747387_file03.zip › uptake plots_cond 1_112025/39 ER[202-231] ADGGYAFKYENGKYDI.png]

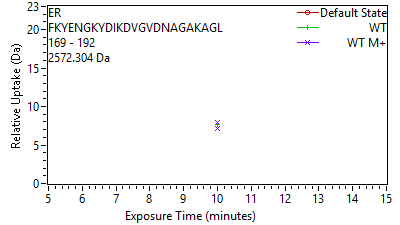

Supplement: Supporting Data File 1 [file 747387_file03.zip › uptake plots_cond 1_112025/40 ER[208-231] FKYENGKYDIKDVGVD.png]

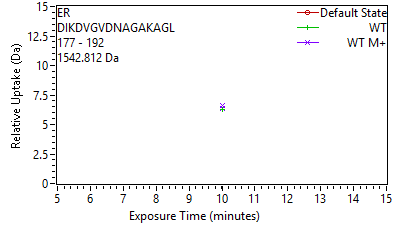

Supplement: Supporting Data File 1 [file 747387_file03.zip › uptake plots_cond 1_112025/41 ER[216-231] DIKDVGVDNAGAKAGL.png]

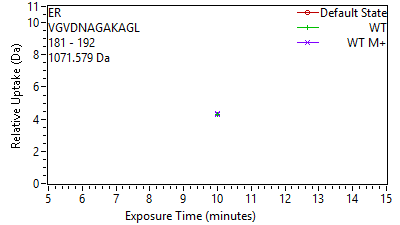

Supplement: Supporting Data File 1 [file 747387_file03.zip › uptake plots_cond 1_112025/42 ER[220-231] VGVDNAGAKAGL.png]

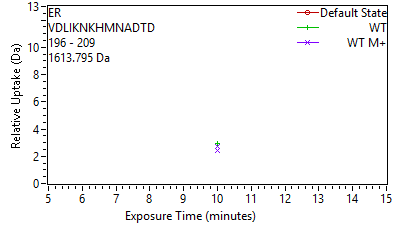

Supplement: Supporting Data File 1 [file 747387_file03.zip › uptake plots_cond 1_112025/43 ER[235-248] VDLIKNKHMNADTD.png]

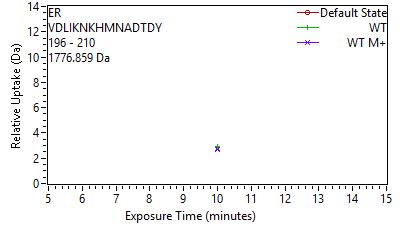

Supplement: Supporting Data File 1 [file 747387_file03.zip › uptake plots_cond 1_112025/44 ER[235-249] VDLIKNKHMNADTDY.png]

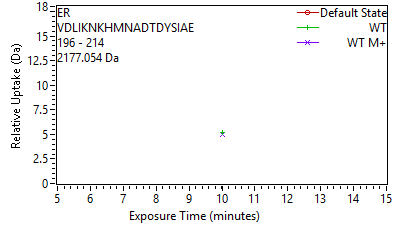

Supplement: Supporting Data File 1 [file 747387_file03.zip › uptake plots_cond 1_112025/45 ER[235-253] VDLIKNKHMNADTDYS.png]

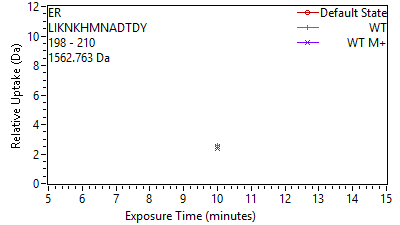

Supplement: Supporting Data File 1 [file 747387_file03.zip › uptake plots_cond 1_112025/46 ER[237-249] LIKNKHMNADTDY.png]

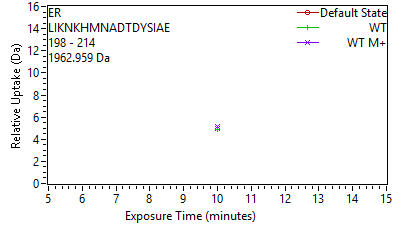

Supplement: Supporting Data File 1 [file 747387_file03.zip › uptake plots_cond 1_112025/47 ER[237-253] LIKNKHMNADTDYSIA.png]

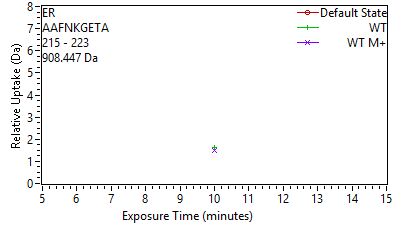

Supplement: Supporting Data File 1 [file 747387_file03.zip › uptake plots_cond 1_112025/48 ER[254-262] AAFNKGETA.png]

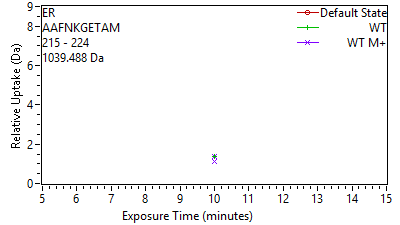

Supplement: Supporting Data File 1 [file 747387_file03.zip › uptake plots_cond 1_112025/49 ER[254-263] AAFNKGETAM.png]

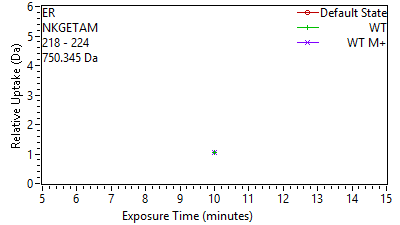

Supplement: Supporting Data File 1 [file 747387_file03.zip › uptake plots_cond 1_112025/50 ER[257-263] NKGETAM.png]

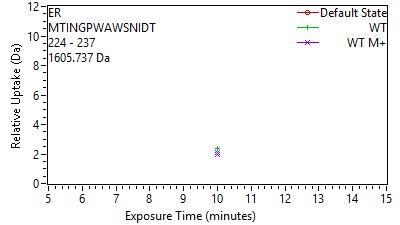

Supplement: Supporting Data File 1 [file 747387_file03.zip › uptake plots_cond 1_112025/51 ER[263-276] MTINGPWAWSNIDT.png]

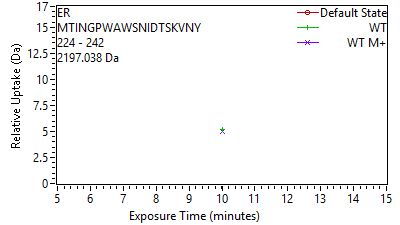

Supplement: Supporting Data File 1 [file 747387_file03.zip › uptake plots_cond 1_112025/52 ER[263-281] MTINGPWAWSNIDTSK.png]

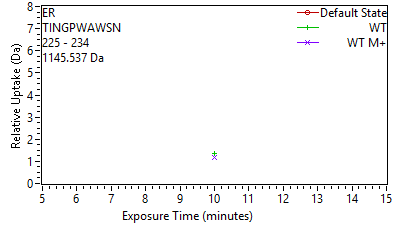

Supplement: Supporting Data File 1 [file 747387_file03.zip › uptake plots_cond 1_112025/53 ER[264-273] TINGPWAWSN.png]

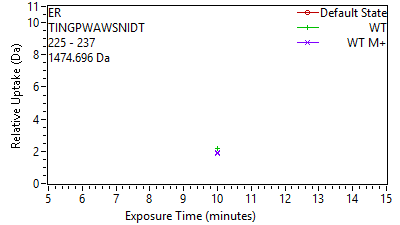

Supplement: Supporting Data File 1 [file 747387_file03.zip › uptake plots_cond 1_112025/54 ER[264-276] TINGPWAWSNIDT.png]

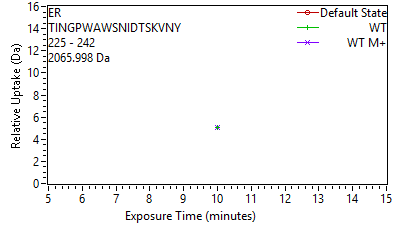

Supplement: Supporting Data File 1 [file 747387_file03.zip › uptake plots_cond 1_112025/55 ER[264-281] TINGPWAWSNIDTSKV.png]

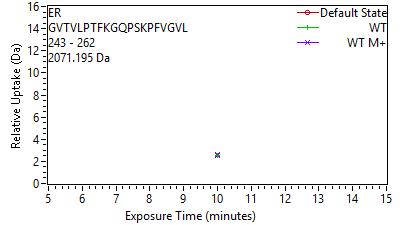

Supplement: Supporting Data File 1 [file 747387_file03.zip › uptake plots_cond 1_112025/56 ER[282-301] GVTVLPTFKGQPSKPF.png]

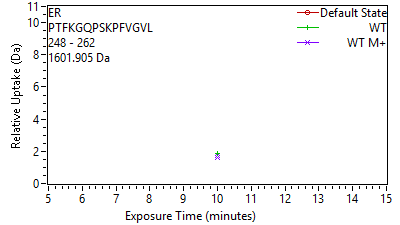

Supplement: Supporting Data File 1 [file 747387_file03.zip › uptake plots_cond 1_112025/57 ER[287-301] PTFKGQPSKPFVGVL.png]

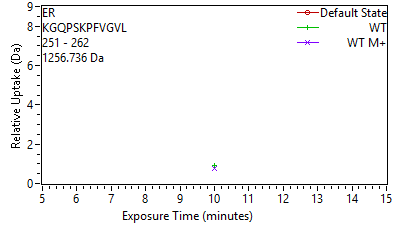

Supplement: Supporting Data File 1 [file 747387_file03.zip › uptake plots_cond 1_112025/58 ER[290-301] KGQPSKPFVGVL.png]

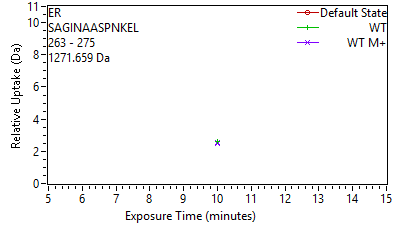

Supplement: Supporting Data File 1 [file 747387_file03.zip › uptake plots_cond 1_112025/59 ER[302-314] SAGINAASPNKEL.png]

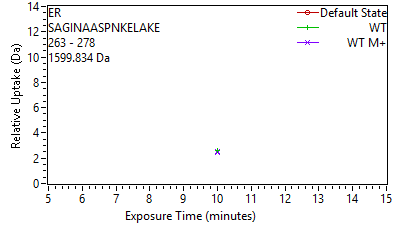

Supplement: Supporting Data File 1 [file 747387_file03.zip › uptake plots_cond 1_112025/60 ER[302-317] SAGINAASPNKELAKE.png]

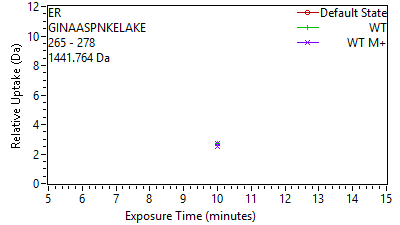

Supplement: Supporting Data File 1 [file 747387_file03.zip › uptake plots_cond 1_112025/61 ER[304-317] GINAASPNKELAKE.png]

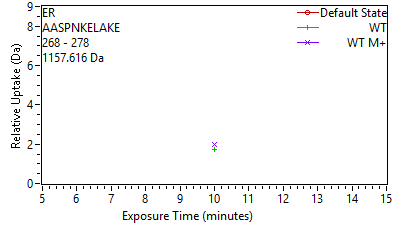

Supplement: Supporting Data File 1 [file 747387_file03.zip › uptake plots_cond 1_112025/62 ER[307-317] AASPNKELAKE.png]

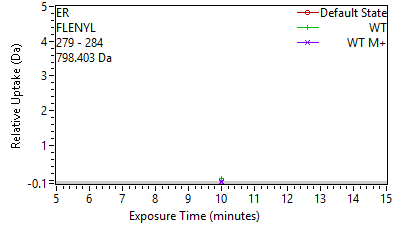

Supplement: Supporting Data File 1 [file 747387_file03.zip › uptake plots_cond 1_112025/63 ER[318-323] FLENYL.png]

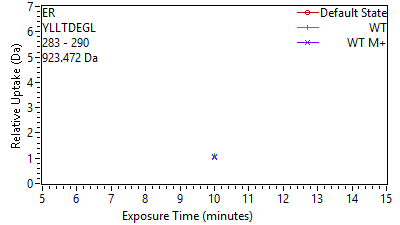

Supplement: Supporting Data File 1 [file 747387_file03.zip › uptake plots_cond 1_112025/64 ER[322-329] YLLTDEGL.png]

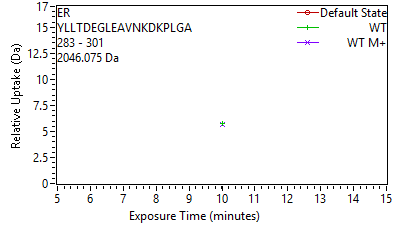

Supplement: Supporting Data File 1 [file 747387_file03.zip › uptake plots_cond 1_112025/65 ER[322-340] YLLTDEGLEAVNKDKP.png]

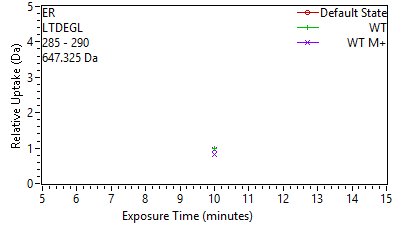

Supplement: Supporting Data File 1 [file 747387_file03.zip › uptake plots_cond 1_112025/66 ER[324-329] LTDEGL.png]

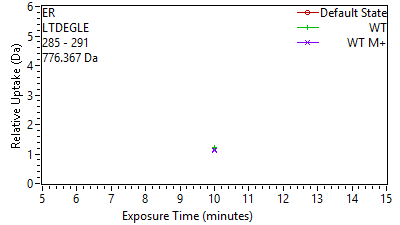

Supplement: Supporting Data File 1 [file 747387_file03.zip › uptake plots_cond 1_112025/67 ER[324-330] LTDEGLE.png]

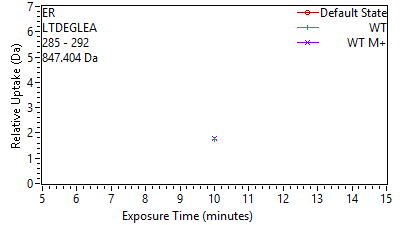

Supplement: Supporting Data File 1 [file 747387_file03.zip › uptake plots_cond 1_112025/68 ER[324-331] LTDEGLEA.png]

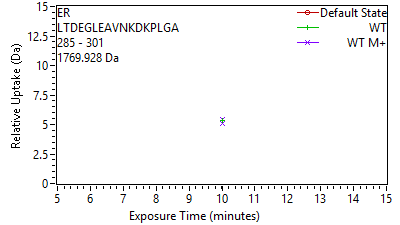

Supplement: Supporting Data File 1 [file 747387_file03.zip › uptake plots_cond 1_112025/69 ER[324-340] LTDEGLEAVNKDKPLG.png]

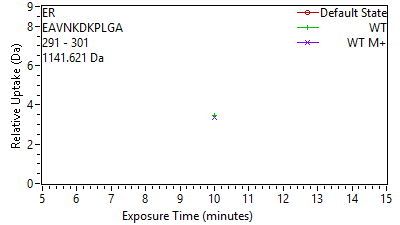

Supplement: Supporting Data File 1 [file 747387_file03.zip › uptake plots_cond 1_112025/70 ER[330-340] EAVNKDKPLGA.png]

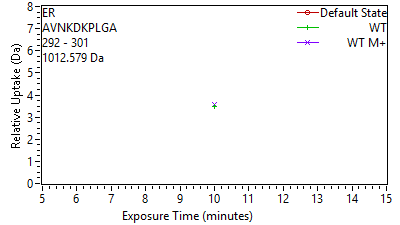

Supplement: Supporting Data File 1 [file 747387_file03.zip › uptake plots_cond 1_112025/71 ER[331-340] AVNKDKPLGA.png]

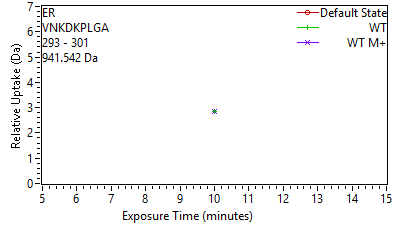

Supplement: Supporting Data File 1 [file 747387_file03.zip › uptake plots_cond 1_112025/72 ER[332-340] VNKDKPLGA.png]

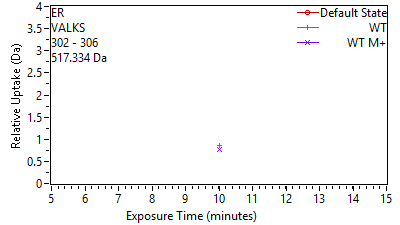

Supplement: Supporting Data File 1 [file 747387_file03.zip › uptake plots_cond 1_112025/73 ER[341-345] VALKS.png]

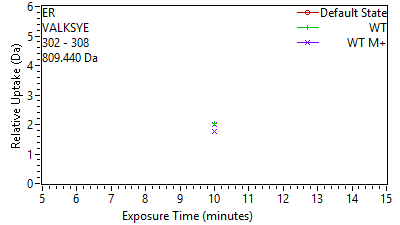

Supplement: Supporting Data File 1 [file 747387_file03.zip › uptake plots_cond 1_112025/74 ER[341-347] VALKSYE.png]

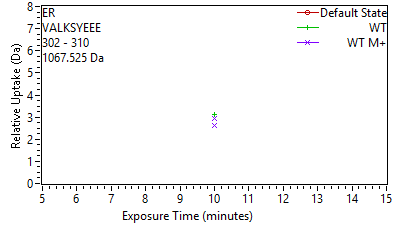

Supplement: Supporting Data File 1 [file 747387_file03.zip › uptake plots_cond 1_112025/75 ER[341-349] VALKSYEEE.png]

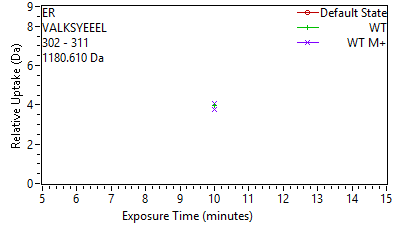

Supplement: Supporting Data File 1 [file 747387_file03.zip › uptake plots_cond 1_112025/76 ER[341-350] VALKSYEEEL.png]

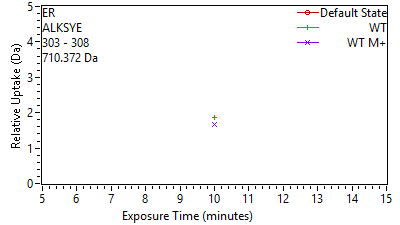

Supplement: Supporting Data File 1 [file 747387_file03.zip › uptake plots_cond 1_112025/77 ER[342-347] ALKSYE.png]

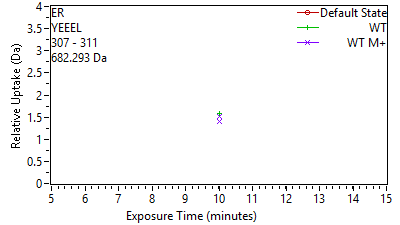

Supplement: Supporting Data File 1 [file 747387_file03.zip › uptake plots_cond 1_112025/78 ER[346-350] YEEEL.png]

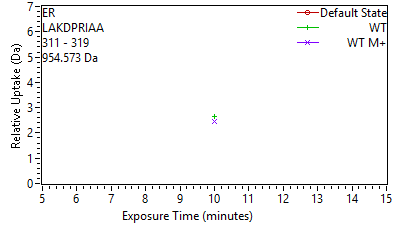

Supplement: Supporting Data File 1 [file 747387_file03.zip › uptake plots_cond 1_112025/79 ER[350-358] LAKDPRIAA.png]

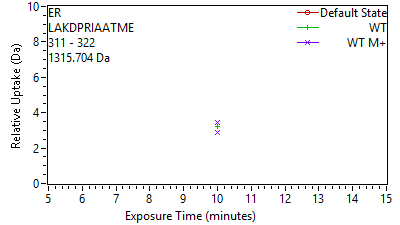

Supplement: Supporting Data File 1 [file 747387_file03.zip › uptake plots_cond 1_112025/80 ER[350-361] LAKDPRIAATME.png]

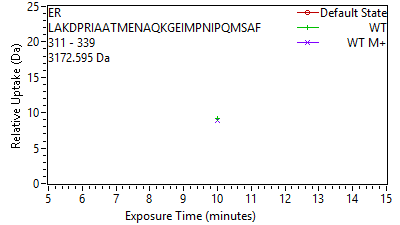

Supplement: Supporting Data File 1 [file 747387_file03.zip › uptake plots_cond 1_112025/81 ER[350-378] LAKDPRIAATMENAQK.png]

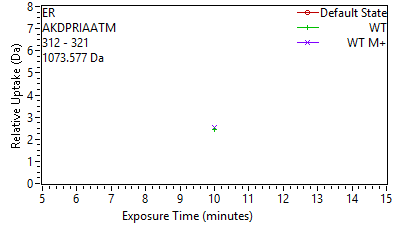

Supplement: Supporting Data File 1 [file 747387_file03.zip › uptake plots_cond 1_112025/82 ER[351-360] AKDPRIAATM.png]

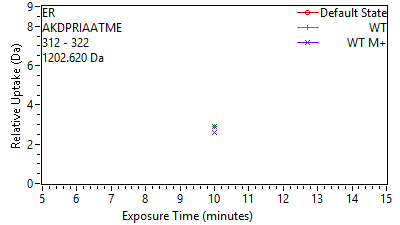

Supplement: Supporting Data File 1 [file 747387_file03.zip › uptake plots_cond 1_112025/83 ER[351-361] AKDPRIAATME.png]

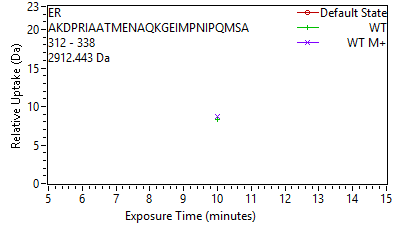

Supplement: Supporting Data File 1 [file 747387_file03.zip › uptake plots_cond 1_112025/84 ER[351-377] AKDPRIAATMENAQKG.png]

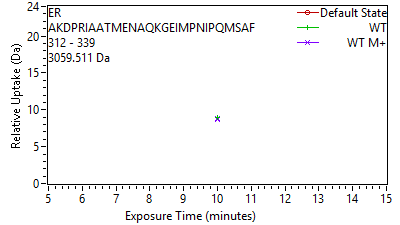

Supplement: Supporting Data File 1 [file 747387_file03.zip › uptake plots_cond 1_112025/85 ER[351-378] AKDPRIAATMENAQKG.png]

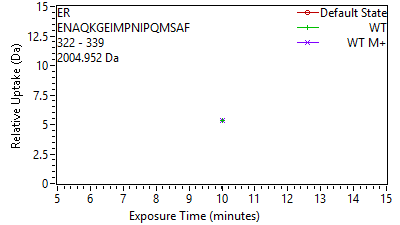

Supplement: Supporting Data File 1 [file 747387_file03.zip › uptake plots_cond 1_112025/86 ER[361-378] ENAQKGEIMPNIPQMS.png]

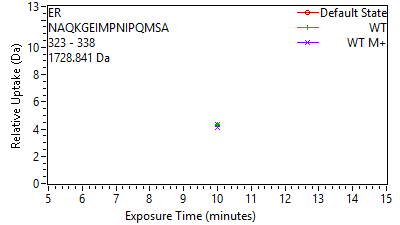

Supplement: Supporting Data File 1 [file 747387_file03.zip › uptake plots_cond 1_112025/87 ER[362-377] NAQKGEIMPNIPQMSA.png]

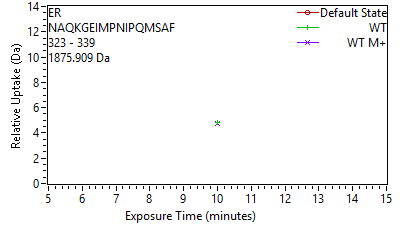

Supplement: Supporting Data File 1 [file 747387_file03.zip › uptake plots_cond 1_112025/88 ER[362-378] NAQKGEIMPNIPQMSA.png]

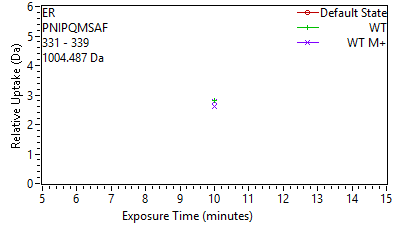

Supplement: Supporting Data File 1 [file 747387_file03.zip › uptake plots_cond 1_112025/89 ER[370-378] PNIPQMSAF.png]

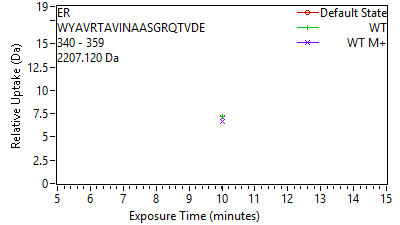

Supplement: Supporting Data File 1 [file 747387_file03.zip › uptake plots_cond 1_112025/90 ER[379-398] WYAVRTAVINAASGRQ.png]

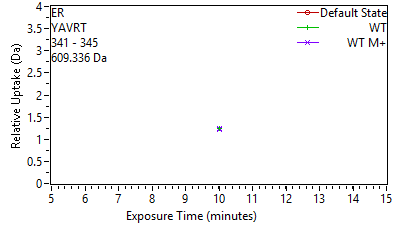

Supplement: Supporting Data File 1 [file 747387_file03.zip › uptake plots_cond 1_112025/91 ER[380-384] YAVRT.png]

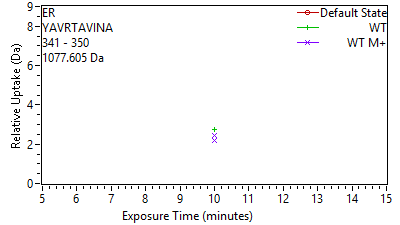

Supplement: Supporting Data File 1 [file 747387_file03.zip › uptake plots_cond 1_112025/92 ER[380-389] YAVRTAVINA.png]

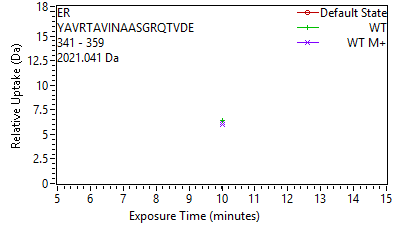

Supplement: Supporting Data File 1 [file 747387_file03.zip › uptake plots_cond 1_112025/93 ER[380-398] YAVRTAVINAASGRQT.png]

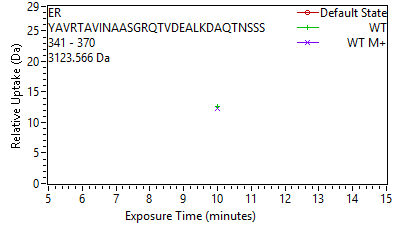

Supplement: Supporting Data File 1 [file 747387_file03.zip › uptake plots_cond 1_112025/94 ER[380-409] YAVRTAVINAASGRQT.png]

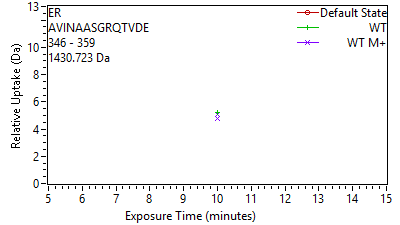

Supplement: Supporting Data File 1 [file 747387_file03.zip › uptake plots_cond 1_112025/95 ER[385-398] AVINAASGRQTVDE.png]

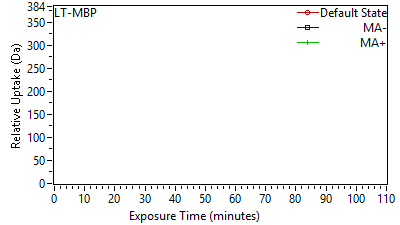

Supplement: Supporting Data File 2 [file 747387_file04.zip › uptake plots_cond 2_022026/001 LT-MBP[2-410] MGSSHHHHHHSSGLVP.png]

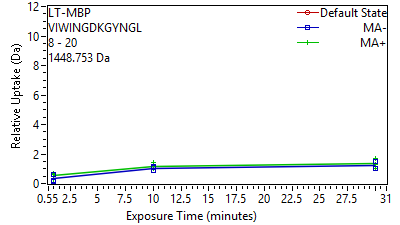

Supplement: Supporting Data File 2 [file 747387_file04.zip › uptake plots_cond 2_022026/002 LT-MBP[47-59] VIWINGDKGYNGL.png]

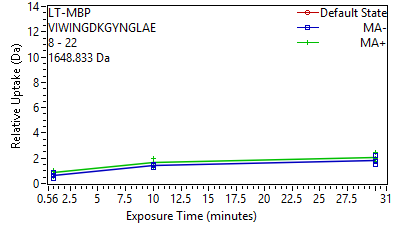

Supplement: Supporting Data File 2 [file 747387_file04.zip › uptake plots_cond 2_022026/003 LT-MBP[47-61] VIWINGDKGYNGLAE.png]

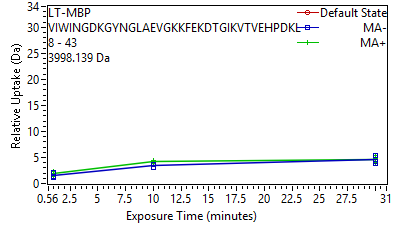

Supplement: Supporting Data File 2 [file 747387_file04.zip › uptake plots_cond 2_022026/004 LT-MBP[47-82] VIWINGDKGYNGLAEV.png]

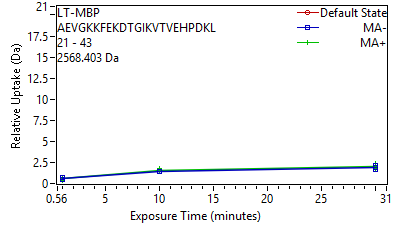

Supplement: Supporting Data File 2 [file 747387_file04.zip › uptake plots_cond 2_022026/005 LT-MBP[60-82] AEVGKKFEKDTGIKVT.png]
