## Supplementary figures and images for "LemonCatcher Acidic Pull-Down Enables Selective In-Cell Hydrogen-Deuterium Exchange Mass Spectrometry"

### Supporting Data File 4

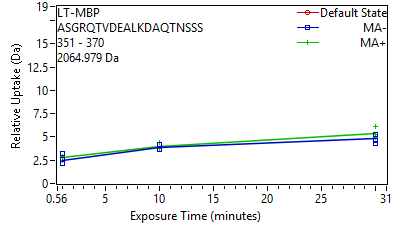

Supplement: Supporting Data File 2 [file 747387_file04.zip › uptake plots_cond 2_022026/106 LT-MBP[390-409] ASGRQTVDEALKDAQT.png]

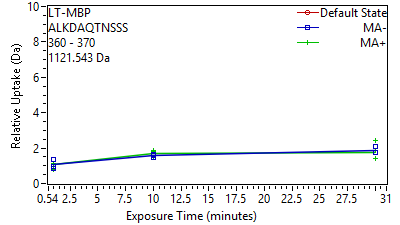

Supplement: Supporting Data File 2 [file 747387_file04.zip › uptake plots_cond 2_022026/107 LT-MBP[399-409] ALKDAQTNSSS.png]

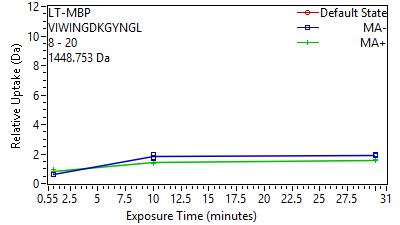

Supplement: Supporting Data File 3 [file 747387_file05.zip › uptake plots_cond 3_042026/02 LT-MBP[47-59] VIWINGDKGYNGL.png]

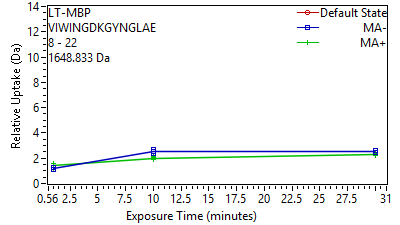

Supplement: Supporting Data File 3 [file 747387_file05.zip › uptake plots_cond 3_042026/03 LT-MBP[47-61] VIWINGDKGYNGLAE.png]

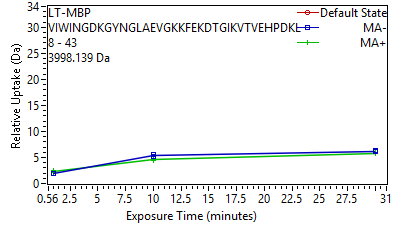

Supplement: Supporting Data File 3 [file 747387_file05.zip › uptake plots_cond 3_042026/04 LT-MBP[47-82] VIWINGDKGYNGLAEV.png]

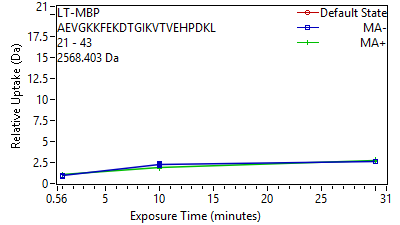

Supplement: Supporting Data File 3 [file 747387_file05.zip › uptake plots_cond 3_042026/05 LT-MBP[60-82] AEVGKKFEKDTGIKVT.png]

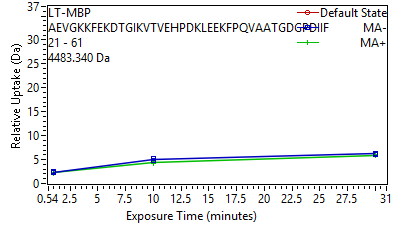

Supplement: Supporting Data File 3 [file 747387_file05.zip › uptake plots_cond 3_042026/06 LT-MBP[60-100] AEVGKKFEKDTGIKVT.png]

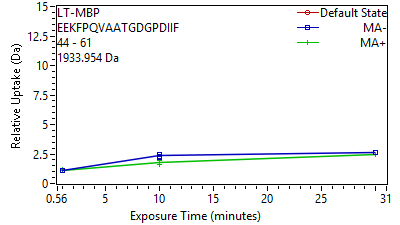

Supplement: Supporting Data File 3 [file 747387_file05.zip › uptake plots_cond 3_042026/07 LT-MBP[83-100] EEKFPQVAATGDGPDI.png]

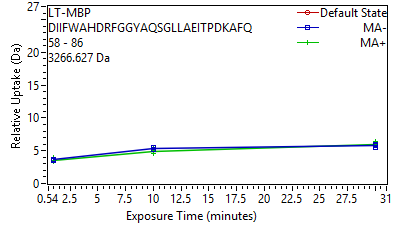

Supplement: Supporting Data File 3 [file 747387_file05.zip › uptake plots_cond 3_042026/08 LT-MBP[97-125] DIIFWAHDRFGGYAQS.png]

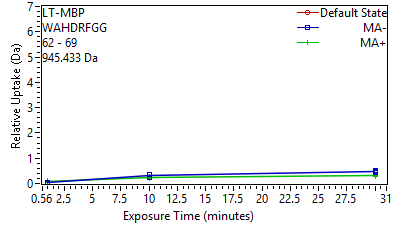

Supplement: Supporting Data File 3 [file 747387_file05.zip › uptake plots_cond 3_042026/09 LT-MBP[101-108] WAHDRFGG.png]

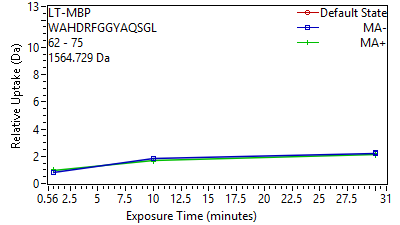

Supplement: Supporting Data File 3 [file 747387_file05.zip › uptake plots_cond 3_042026/10 LT-MBP[101-114] WAHDRFGGYAQSGL.png]

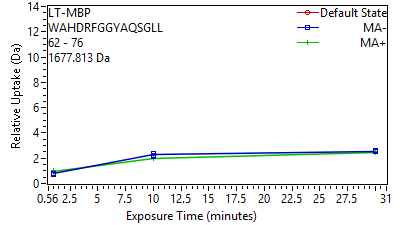

Supplement: Supporting Data File 3 [file 747387_file05.zip › uptake plots_cond 3_042026/11 LT-MBP[101-115] WAHDRFGGYAQSGLL.png]

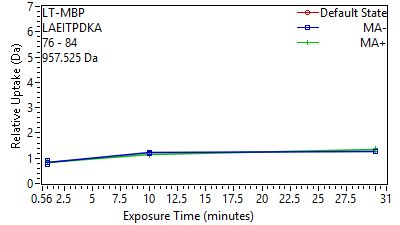

Supplement: Supporting Data File 3 [file 747387_file05.zip › uptake plots_cond 3_042026/12 LT-MBP[115-123] LAEITPDKA.png]

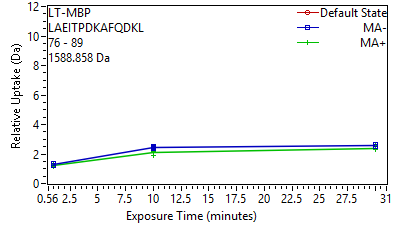

Supplement: Supporting Data File 3 [file 747387_file05.zip › uptake plots_cond 3_042026/13 LT-MBP[115-128] LAEITPDKAFQDKL.png]

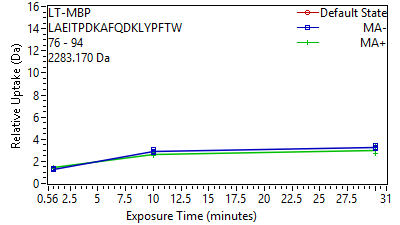

Supplement: Supporting Data File 3 [file 747387_file05.zip › uptake plots_cond 3_042026/14 LT-MBP[115-133] LAEITPDKAFQDKLYP.png]

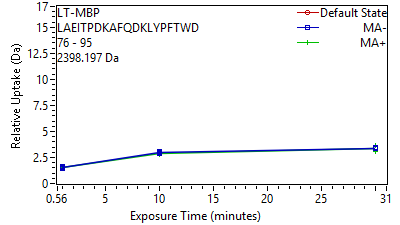

Supplement: Supporting Data File 3 [file 747387_file05.zip › uptake plots_cond 3_042026/15 LT-MBP[115-134] LAEITPDKAFQDKLYP.png]

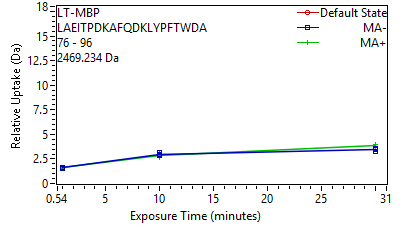

Supplement: Supporting Data File 3 [file 747387_file05.zip › uptake plots_cond 3_042026/16 LT-MBP[115-135] LAEITPDKAFQDKLYP.png]

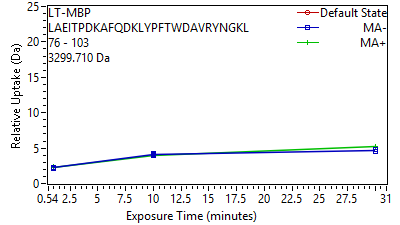

Supplement: Supporting Data File 3 [file 747387_file05.zip › uptake plots_cond 3_042026/17 LT-MBP[115-142] LAEITPDKAFQDKLYP.png]

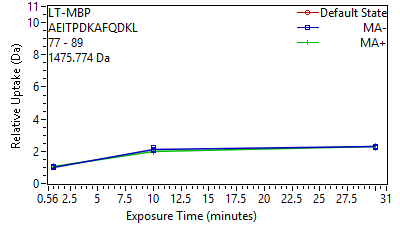

Supplement: Supporting Data File 3 [file 747387_file05.zip › uptake plots_cond 3_042026/18 LT-MBP[116-128] AEITPDKAFQDKL.png]

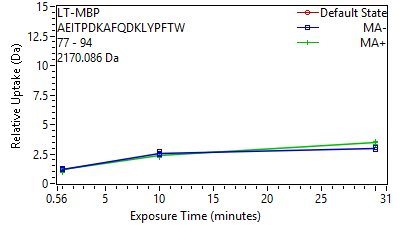

Supplement: Supporting Data File 3 [file 747387_file05.zip › uptake plots_cond 3_042026/19 LT-MBP[116-133] AEITPDKAFQDKLYPF.png]

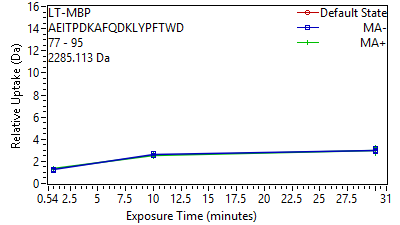

Supplement: Supporting Data File 3 [file 747387_file05.zip › uptake plots_cond 3_042026/20 LT-MBP[116-134] AEITPDKAFQDKLYPF.png]

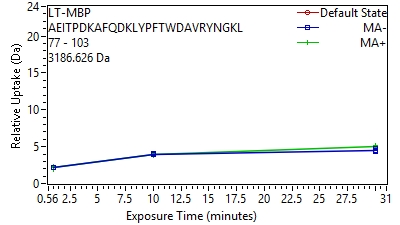

Supplement: Supporting Data File 3 [file 747387_file05.zip › uptake plots_cond 3_042026/21 LT-MBP[116-142] AEITPDKAFQDKLYPF.png]

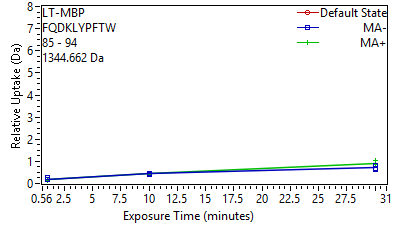

Supplement: Supporting Data File 3 [file 747387_file05.zip › uptake plots_cond 3_042026/22 LT-MBP[124-133] FQDKLYPFTW.png]

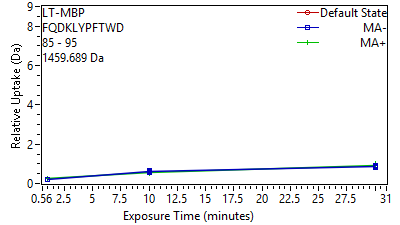

Supplement: Supporting Data File 3 [file 747387_file05.zip › uptake plots_cond 3_042026/23 LT-MBP[124-134] FQDKLYPFTWD.png]

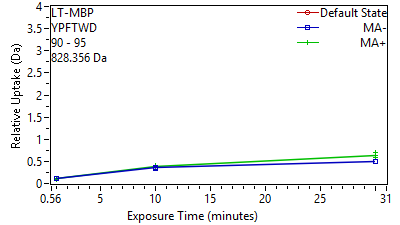

Supplement: Supporting Data File 3 [file 747387_file05.zip › uptake plots_cond 3_042026/24 LT-MBP[129-134] YPFTWD.png]

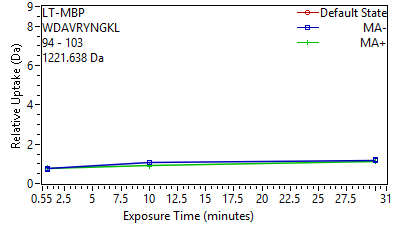

Supplement: Supporting Data File 3 [file 747387_file05.zip › uptake plots_cond 3_042026/25 LT-MBP[133-142] WDAVRYNGKL.png]

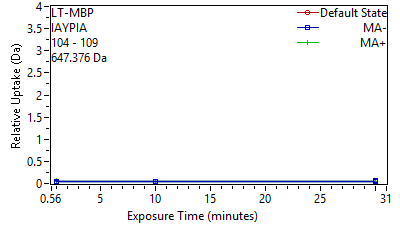

Supplement: Supporting Data File 3 [file 747387_file05.zip › uptake plots_cond 3_042026/26 LT-MBP[143-148] IAYPIA.png]

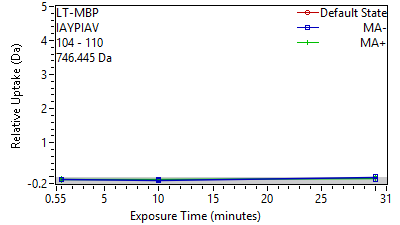

Supplement: Supporting Data File 3 [file 747387_file05.zip › uptake plots_cond 3_042026/27 LT-MBP[143-149] IAYPIAV.png]

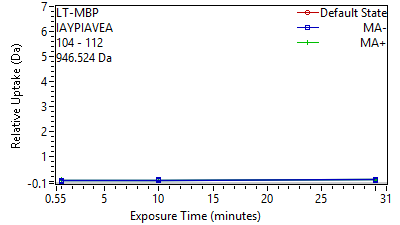

Supplement: Supporting Data File 3 [file 747387_file05.zip › uptake plots_cond 3_042026/28 LT-MBP[143-151] IAYPIAVEA.png]

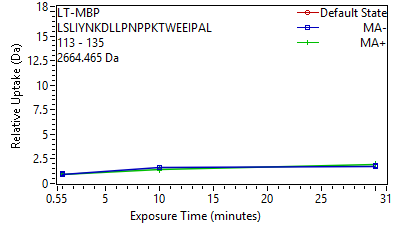

Supplement: Supporting Data File 3 [file 747387_file05.zip › uptake plots_cond 3_042026/29 LT-MBP[152-174] LSLIYNKDLLPNPPKT.png]

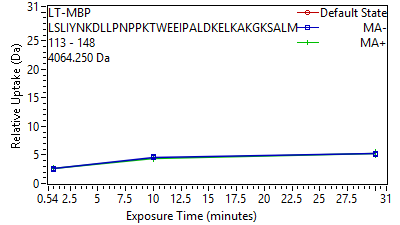

Supplement: Supporting Data File 3 [file 747387_file05.zip › uptake plots_cond 3_042026/30 LT-MBP[152-187] LSLIYNKDLLPNPPKT.png]

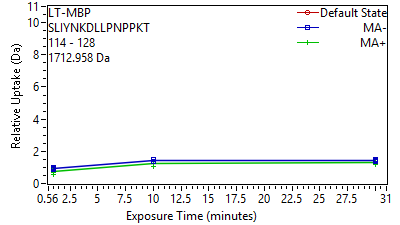

Supplement: Supporting Data File 3 [file 747387_file05.zip › uptake plots_cond 3_042026/31 LT-MBP[153-167] SLIYNKDLLPNPPKT.png]

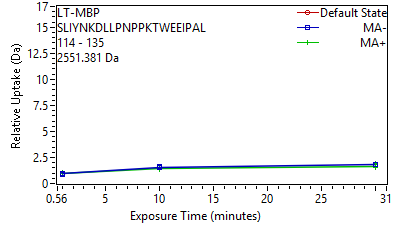

Supplement: Supporting Data File 3 [file 747387_file05.zip › uptake plots_cond 3_042026/32 LT-MBP[153-174] SLIYNKDLLPNPPKTW.png]

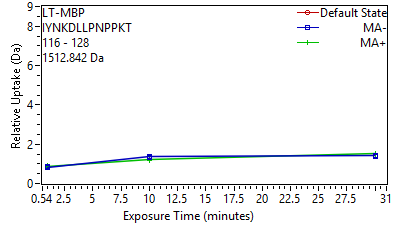

Supplement: Supporting Data File 3 [file 747387_file05.zip › uptake plots_cond 3_042026/33 LT-MBP[155-167] IYNKDLLPNPPKT.png]

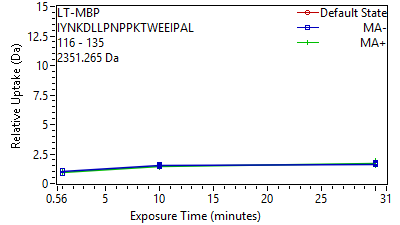

Supplement: Supporting Data File 3 [file 747387_file05.zip › uptake plots_cond 3_042026/34 LT-MBP[155-174] IYNKDLLPNPPKTWEE.png]

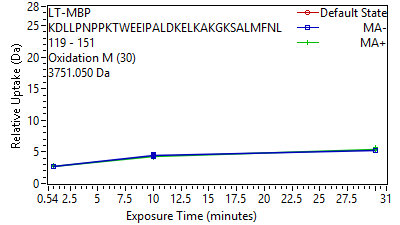

Supplement: Supporting Data File 3 [file 747387_file05.zip › uptake plots_cond 3_042026/35 LT-MBP[158-190] KDLLPNPPKTWEEIPA.png]

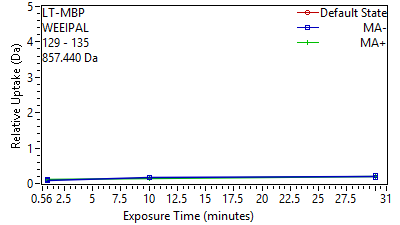

Supplement: Supporting Data File 3 [file 747387_file05.zip › uptake plots_cond 3_042026/36 LT-MBP[168-174] WEEIPAL.png]

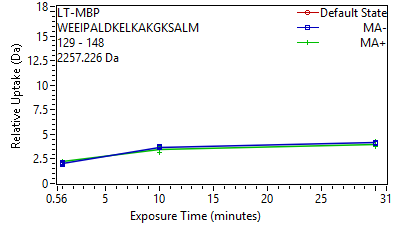

Supplement: Supporting Data File 3 [file 747387_file05.zip › uptake plots_cond 3_042026/37 LT-MBP[168-187] WEEIPALDKELKAKGK.png]

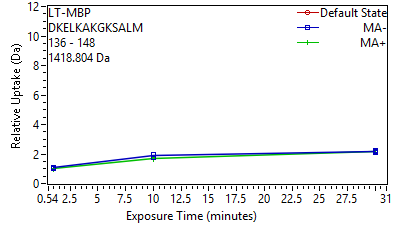

Supplement: Supporting Data File 3 [file 747387_file05.zip › uptake plots_cond 3_042026/38 LT-MBP[175-187] DKELKAKGKSALM.png]

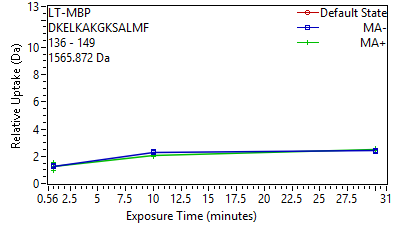

Supplement: Supporting Data File 3 [file 747387_file05.zip › uptake plots_cond 3_042026/39 LT-MBP[175-188] DKELKAKGKSALMF.png]

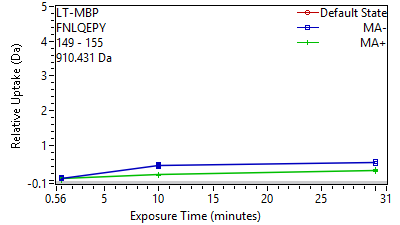

Supplement: Supporting Data File 3 [file 747387_file05.zip › uptake plots_cond 3_042026/40 LT-MBP[188-194] FNLQEPY.png]

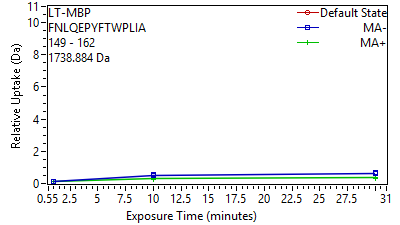

Supplement: Supporting Data File 3 [file 747387_file05.zip › uptake plots_cond 3_042026/41 LT-MBP[188-201] FNLQEPYFTWPLIA.png]

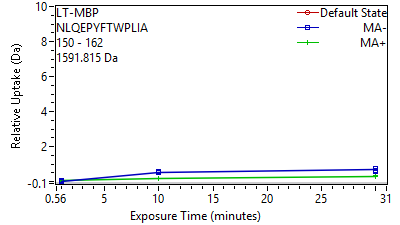

Supplement: Supporting Data File 3 [file 747387_file05.zip › uptake plots_cond 3_042026/42 LT-MBP[189-201] NLQEPYFTWPLIA.png]

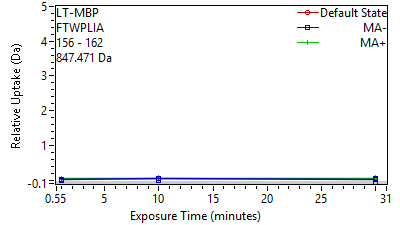

Supplement: Supporting Data File 3 [file 747387_file05.zip › uptake plots_cond 3_042026/43 LT-MBP[195-201] FTWPLIA.png]

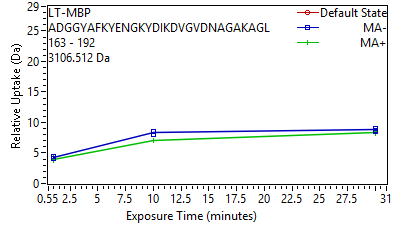

Supplement: Supporting Data File 3 [file 747387_file05.zip › uptake plots_cond 3_042026/44 LT-MBP[202-231] ADGGYAFKYENGKYDI.png]

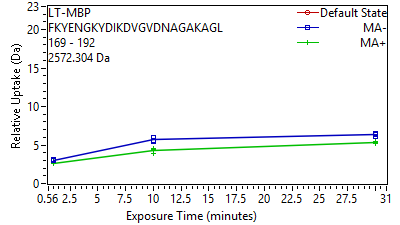

Supplement: Supporting Data File 3 [file 747387_file05.zip › uptake plots_cond 3_042026/45 LT-MBP[208-231] FKYENGKYDIKDVGVD.png]

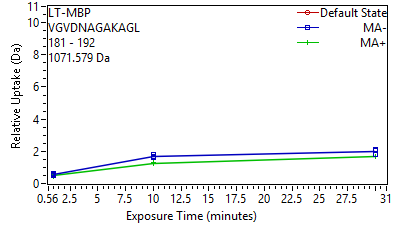

Supplement: Supporting Data File 3 [file 747387_file05.zip › uptake plots_cond 3_042026/46 LT-MBP[220-231] VGVDNAGAKAGL.png]

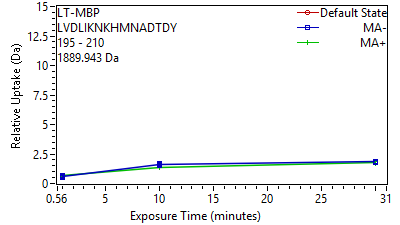

Supplement: Supporting Data File 3 [file 747387_file05.zip › uptake plots_cond 3_042026/47 LT-MBP[234-249] LVDLIKNKHMNADTDY.png]

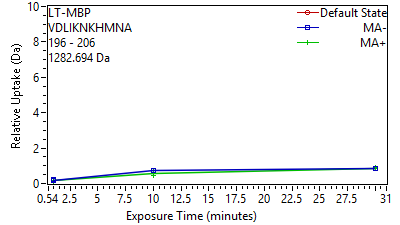

Supplement: Supporting Data File 3 [file 747387_file05.zip › uptake plots_cond 3_042026/48 LT-MBP[235-245] VDLIKNKHMNA.png]

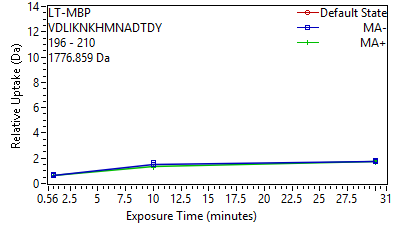

Supplement: Supporting Data File 3 [file 747387_file05.zip › uptake plots_cond 3_042026/49 LT-MBP[235-249] VDLIKNKHMNADTDY.png]

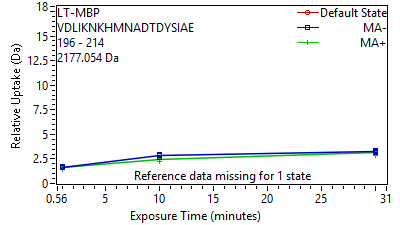

Supplement: Supporting Data File 3 [file 747387_file05.zip › uptake plots_cond 3_042026/50 LT-MBP[235-253] VDLIKNKHMNADTDYS.png]

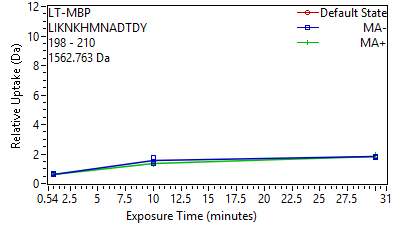

Supplement: Supporting Data File 3 [file 747387_file05.zip › uptake plots_cond 3_042026/51 LT-MBP[237-249] LIKNKHMNADTDY.png]

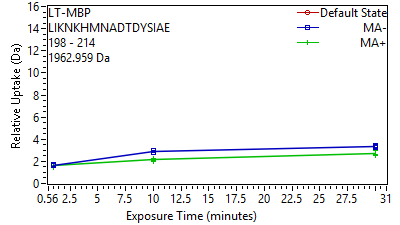

Supplement: Supporting Data File 3 [file 747387_file05.zip › uptake plots_cond 3_042026/52 LT-MBP[237-253] LIKNKHMNADTDYSIA.png]

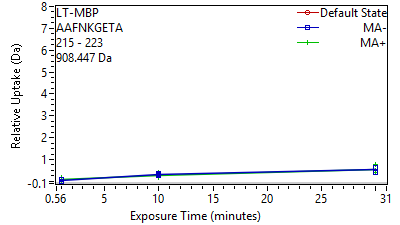

Supplement: Supporting Data File 3 [file 747387_file05.zip › uptake plots_cond 3_042026/53 LT-MBP[254-262] AAFNKGETA.png]

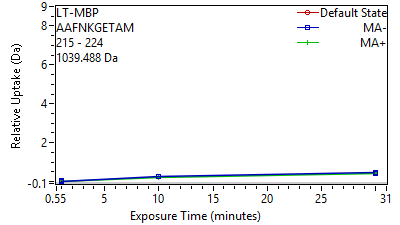

Supplement: Supporting Data File 3 [file 747387_file05.zip › uptake plots_cond 3_042026/54 LT-MBP[254-263] AAFNKGETAM.png]

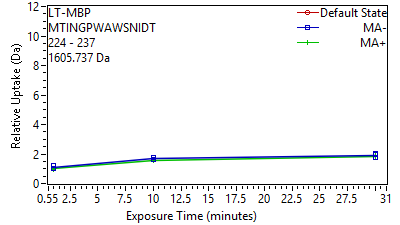

Supplement: Supporting Data File 3 [file 747387_file05.zip › uptake plots_cond 3_042026/55 LT-MBP[263-276] MTINGPWAWSNIDT.png]

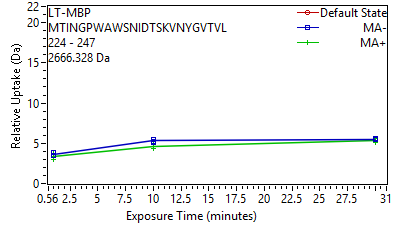

Supplement: Supporting Data File 3 [file 747387_file05.zip › uptake plots_cond 3_042026/56 LT-MBP[263-286] MTINGPWAWSNIDTSK.png]

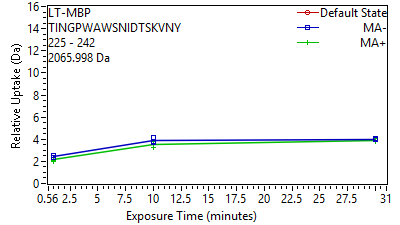

Supplement: Supporting Data File 3 [file 747387_file05.zip › uptake plots_cond 3_042026/57 LT-MBP[264-281] TINGPWAWSNIDTSKV.png]

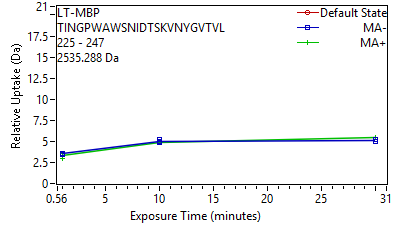

Supplement: Supporting Data File 3 [file 747387_file05.zip › uptake plots_cond 3_042026/58 LT-MBP[264-286] TINGPWAWSNIDTSKV.png]

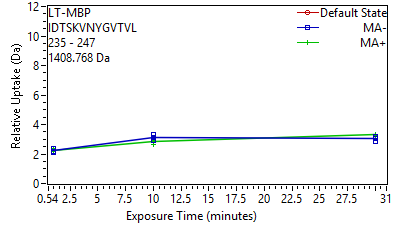

Supplement: Supporting Data File 3 [file 747387_file05.zip › uptake plots_cond 3_042026/59 LT-MBP[274-286] IDTSKVNYGVTVL.png]

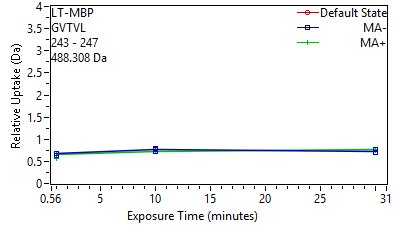

Supplement: Supporting Data File 3 [file 747387_file05.zip › uptake plots_cond 3_042026/60 LT-MBP[282-286] GVTVL.png]

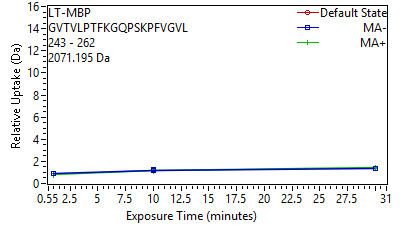

Supplement: Supporting Data File 3 [file 747387_file05.zip › uptake plots_cond 3_042026/61 LT-MBP[282-301] GVTVLPTFKGQPSKPF.png]

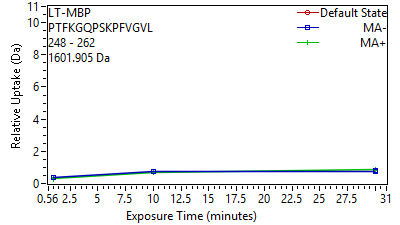

Supplement: Supporting Data File 3 [file 747387_file05.zip › uptake plots_cond 3_042026/62 LT-MBP[287-301] PTFKGQPSKPFVGVL.png]

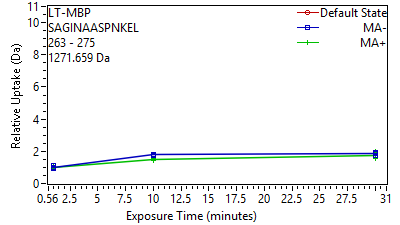

Supplement: Supporting Data File 3 [file 747387_file05.zip › uptake plots_cond 3_042026/63 LT-MBP[302-314] SAGINAASPNKEL.png]

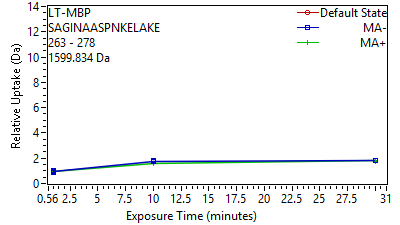

Supplement: Supporting Data File 3 [file 747387_file05.zip › uptake plots_cond 3_042026/64 LT-MBP[302-317] SAGINAASPNKELAKE.png]

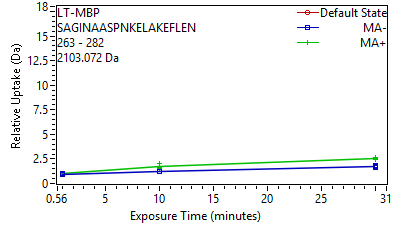

Supplement: Supporting Data File 3 [file 747387_file05.zip › uptake plots_cond 3_042026/65 LT-MBP[302-321] SAGINAASPNKELAKE.png]

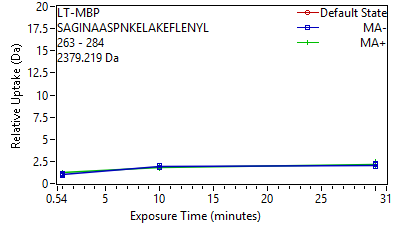

Supplement: Supporting Data File 3 [file 747387_file05.zip › uptake plots_cond 3_042026/66 LT-MBP[302-323] SAGINAASPNKELAKE.png]

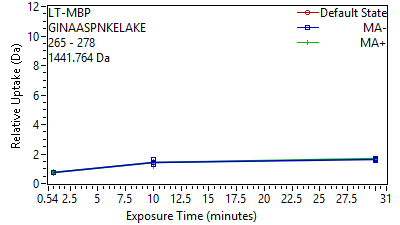

Supplement: Supporting Data File 3 [file 747387_file05.zip › uptake plots_cond 3_042026/67 LT-MBP[304-317] GINAASPNKELAKE.png]

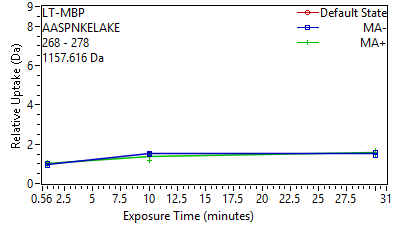

Supplement: Supporting Data File 3 [file 747387_file05.zip › uptake plots_cond 3_042026/68 LT-MBP[307-317] AASPNKELAKE.png]

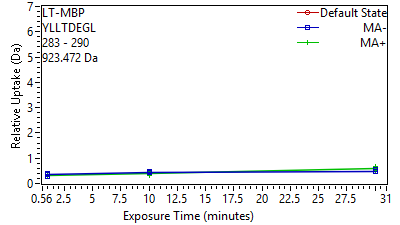

Supplement: Supporting Data File 3 [file 747387_file05.zip › uptake plots_cond 3_042026/69 LT-MBP[322-329] YLLTDEGL.png]

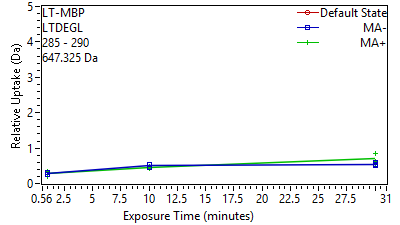

Supplement: Supporting Data File 3 [file 747387_file05.zip › uptake plots_cond 3_042026/70 LT-MBP[324-329] LTDEGL.png]

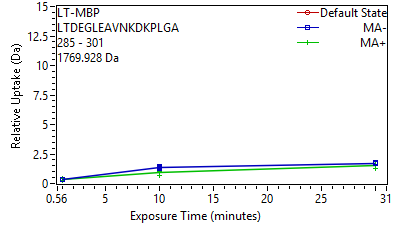

Supplement: Supporting Data File 3 [file 747387_file05.zip › uptake plots_cond 3_042026/71 LT-MBP[324-340] LTDEGLEAVNKDKPLG.png]

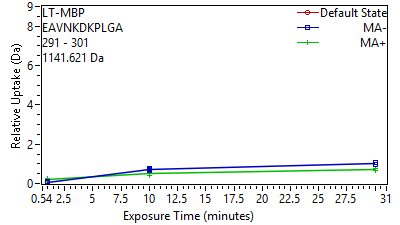

Supplement: Supporting Data File 3 [file 747387_file05.zip › uptake plots_cond 3_042026/72 LT-MBP[330-340] EAVNKDKPLGA.png]

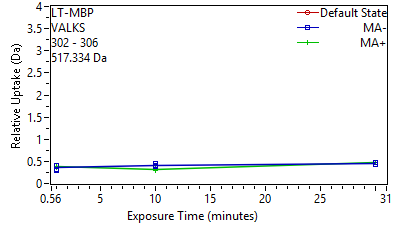

Supplement: Supporting Data File 3 [file 747387_file05.zip › uptake plots_cond 3_042026/73 LT-MBP[341-345] VALKS.png]

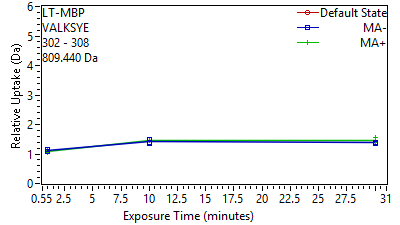

Supplement: Supporting Data File 3 [file 747387_file05.zip › uptake plots_cond 3_042026/74 LT-MBP[341-347] VALKSYE.png]

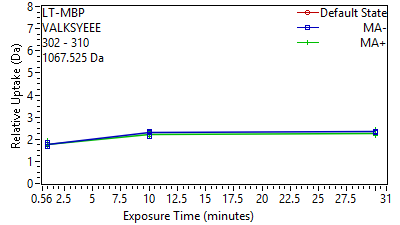

Supplement: Supporting Data File 3 [file 747387_file05.zip › uptake plots_cond 3_042026/75 LT-MBP[341-349] VALKSYEEE.png]

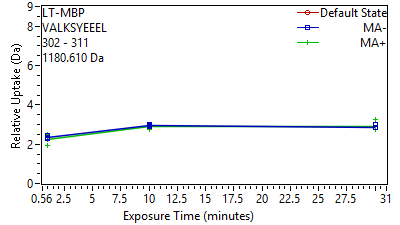

Supplement: Supporting Data File 3 [file 747387_file05.zip › uptake plots_cond 3_042026/76 LT-MBP[341-350] VALKSYEEEL.png]

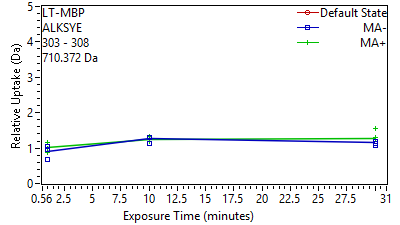

Supplement: Supporting Data File 3 [file 747387_file05.zip › uptake plots_cond 3_042026/77 LT-MBP[342-347] ALKSYE.png]

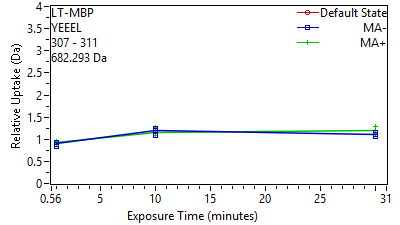

Supplement: Supporting Data File 3 [file 747387_file05.zip › uptake plots_cond 3_042026/78 LT-MBP[346-350] YEEEL.png]

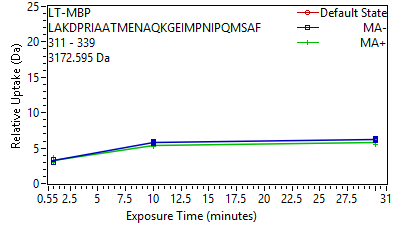

Supplement: Supporting Data File 3 [file 747387_file05.zip › uptake plots_cond 3_042026/79 LT-MBP[350-378] LAKDPRIAATMENAQK.png]

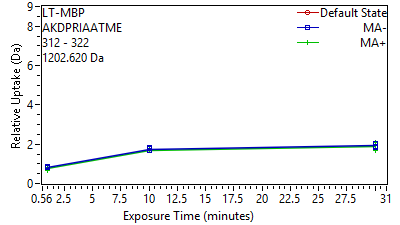

Supplement: Supporting Data File 3 [file 747387_file05.zip › uptake plots_cond 3_042026/80 LT-MBP[351-361] AKDPRIAATME.png]

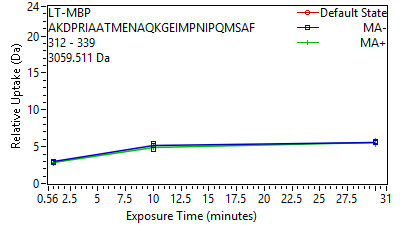

Supplement: Supporting Data File 3 [file 747387_file05.zip › uptake plots_cond 3_042026/81 LT-MBP[351-378] AKDPRIAATMENAQKG.png]

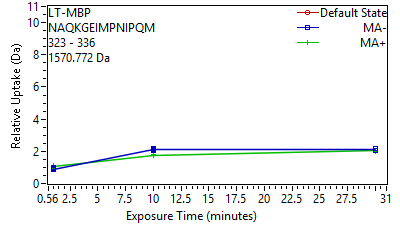

Supplement: Supporting Data File 3 [file 747387_file05.zip › uptake plots_cond 3_042026/82 LT-MBP[362-375] NAQKGEIMPNIPQM.png]

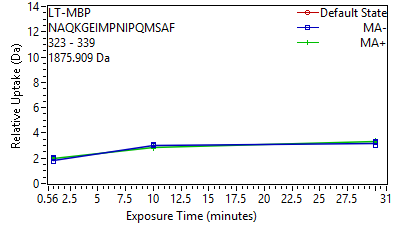

Supplement: Supporting Data File 3 [file 747387_file05.zip › uptake plots_cond 3_042026/83 LT-MBP[362-378] NAQKGEIMPNIPQMSA.png]

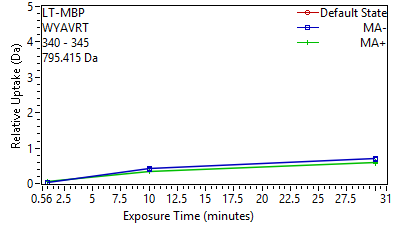

Supplement: Supporting Data File 3 [file 747387_file05.zip › uptake plots_cond 3_042026/84 LT-MBP[379-384] WYAVRT.png]

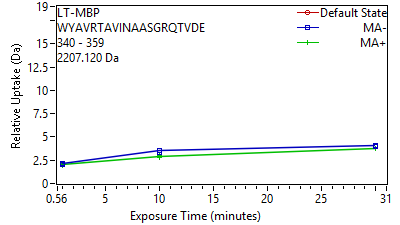

Supplement: Supporting Data File 3 [file 747387_file05.zip › uptake plots_cond 3_042026/85 LT-MBP[379-398] WYAVRTAVINAASGRQ.png]

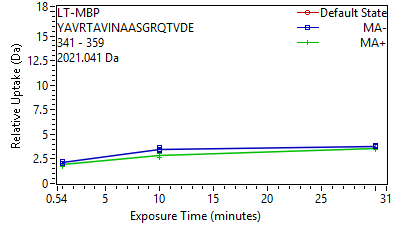

Supplement: Supporting Data File 3 [file 747387_file05.zip › uptake plots_cond 3_042026/86 LT-MBP[380-398] YAVRTAVINAASGRQT.png]

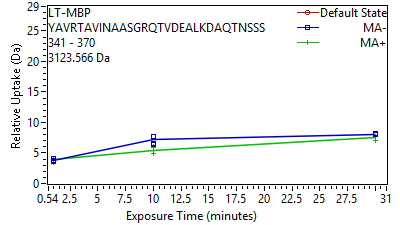

Supplement: Supporting Data File 3 [file 747387_file05.zip › uptake plots_cond 3_042026/87 LT-MBP[380-409] YAVRTAVINAASGRQT.png]

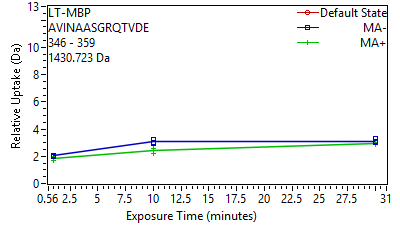

Supplement: Supporting Data File 3 [file 747387_file05.zip › uptake plots_cond 3_042026/88 LT-MBP[385-398] AVINAASGRQTVDE.png]

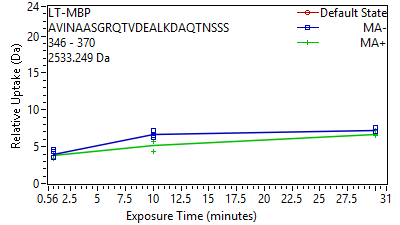

Supplement: Supporting Data File 3 [file 747387_file05.zip › uptake plots_cond 3_042026/89 LT-MBP[385-409] AVINAASGRQTVDEAL.png]

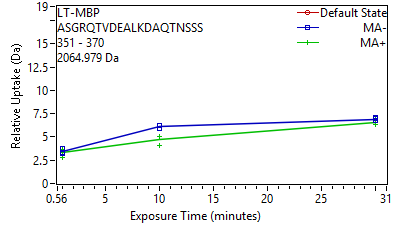

Supplement: Supporting Data File 3 [file 747387_file05.zip › uptake plots_cond 3_042026/90 LT-MBP[390-409] ASGRQTVDEALKDAQT.png]

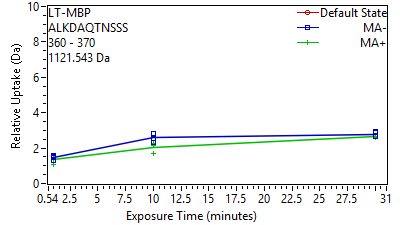

Supplement: Supporting Data File 3 [file 747387_file05.zip › uptake plots_cond 3_042026/91 LT-MBP[399-409] ALKDAQTNSSS.png]

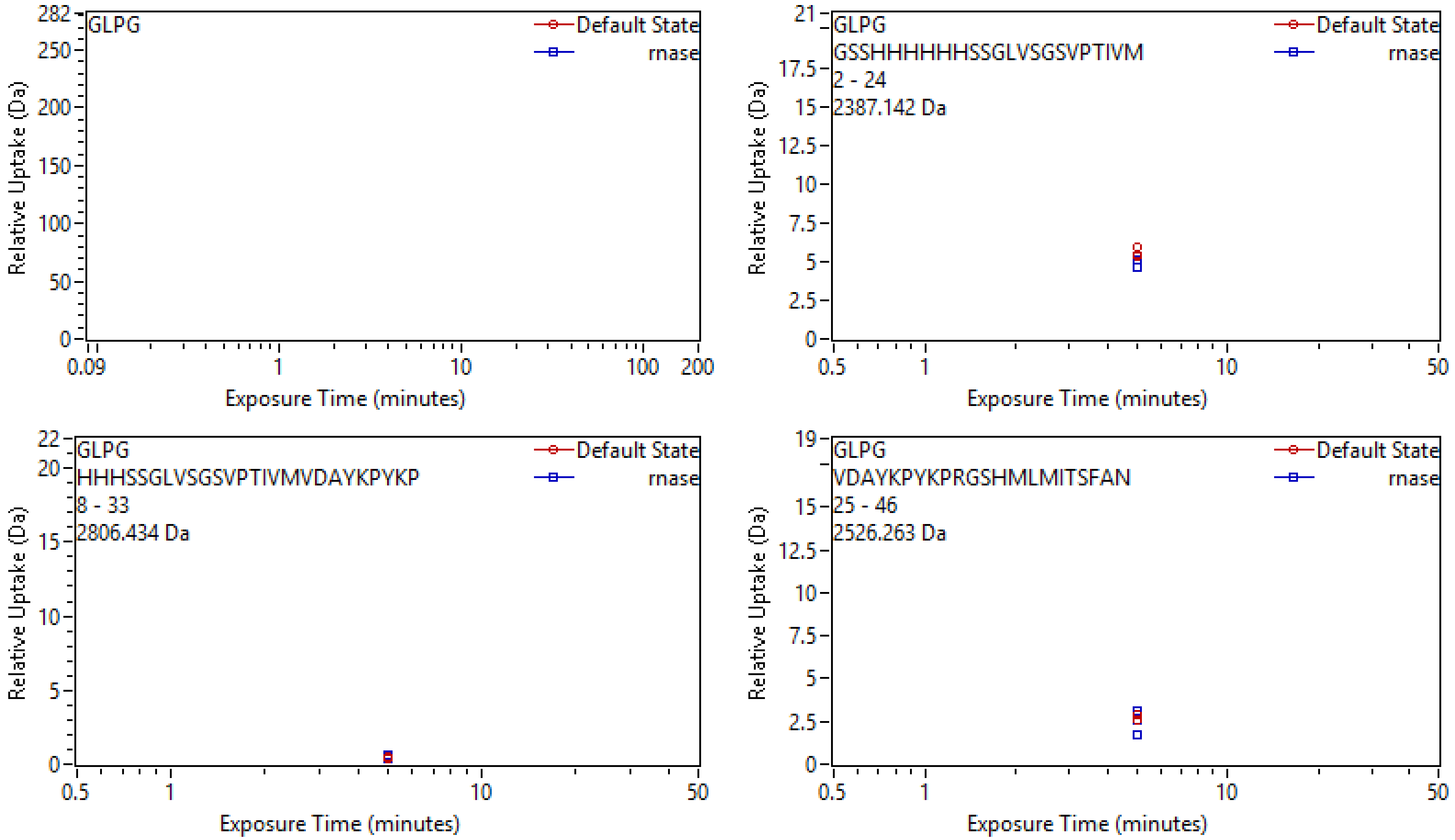

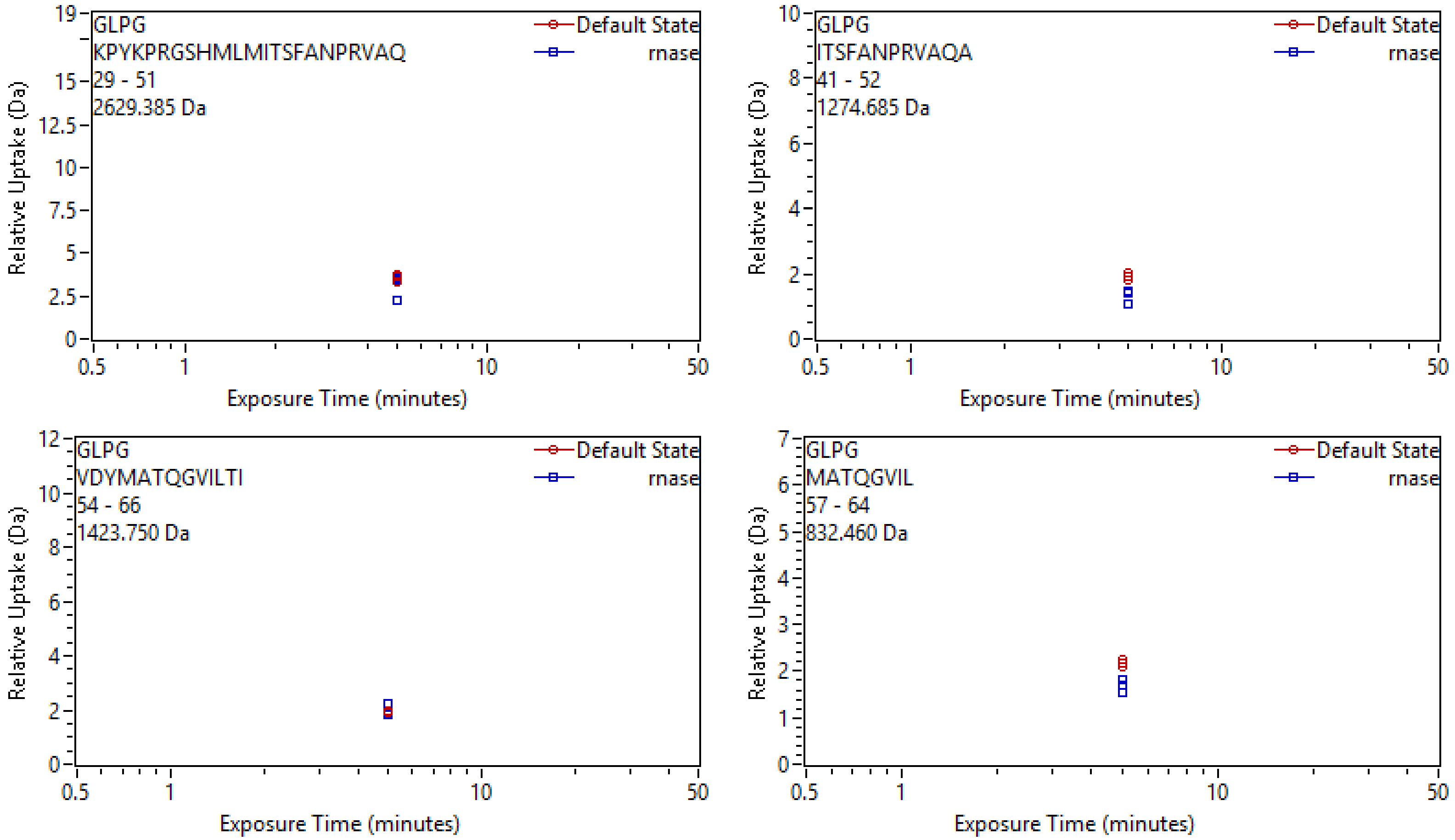

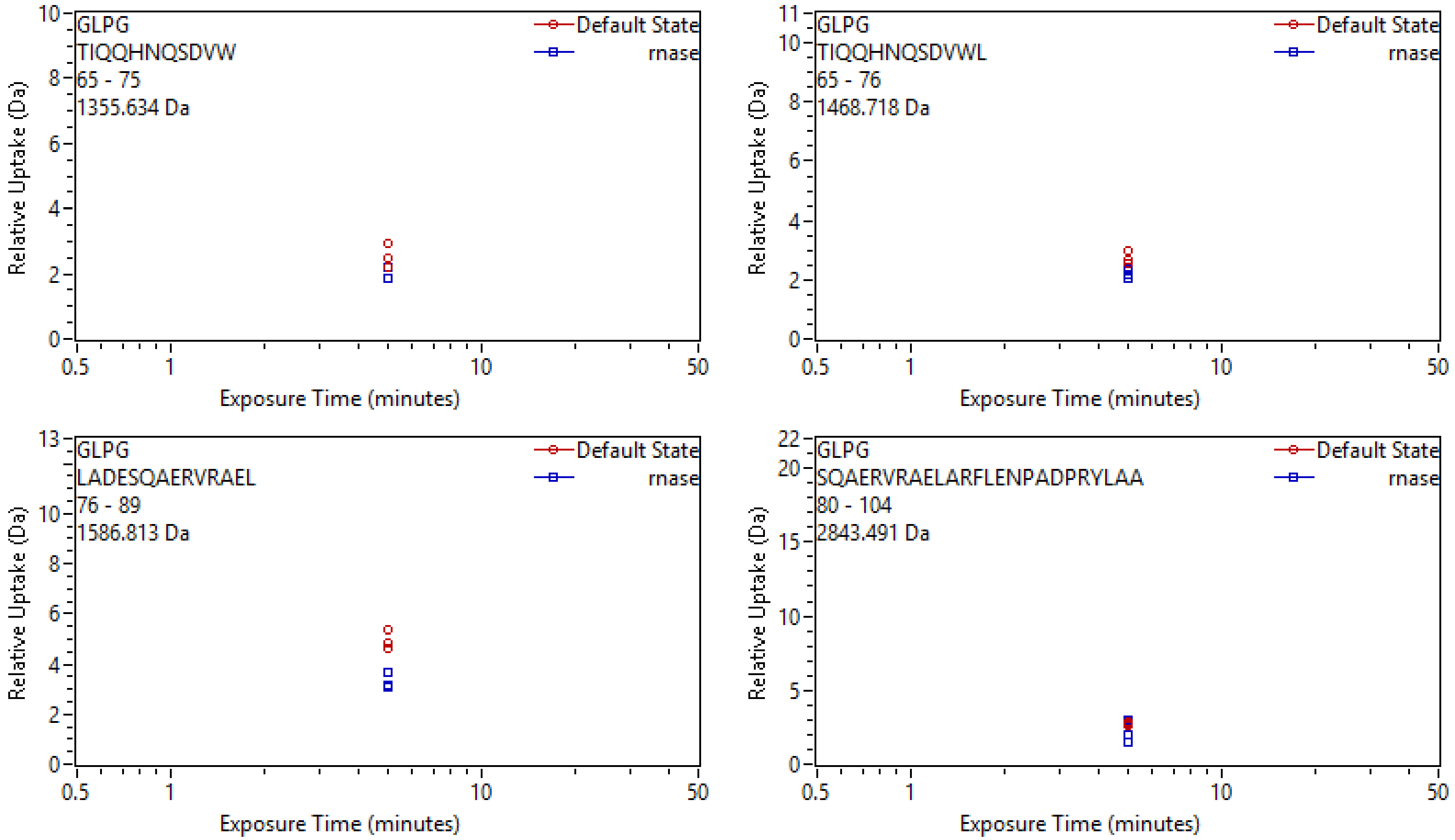

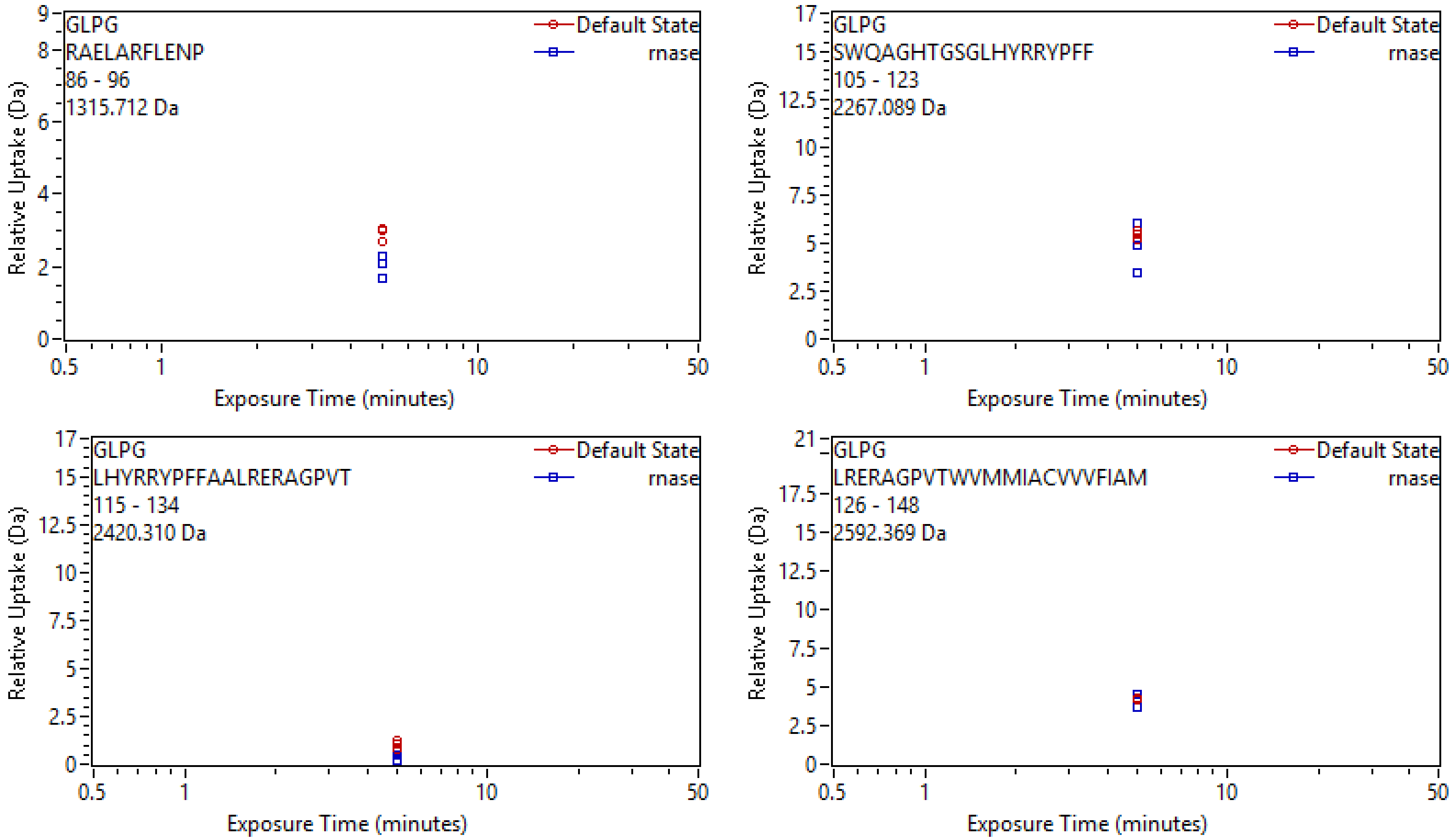

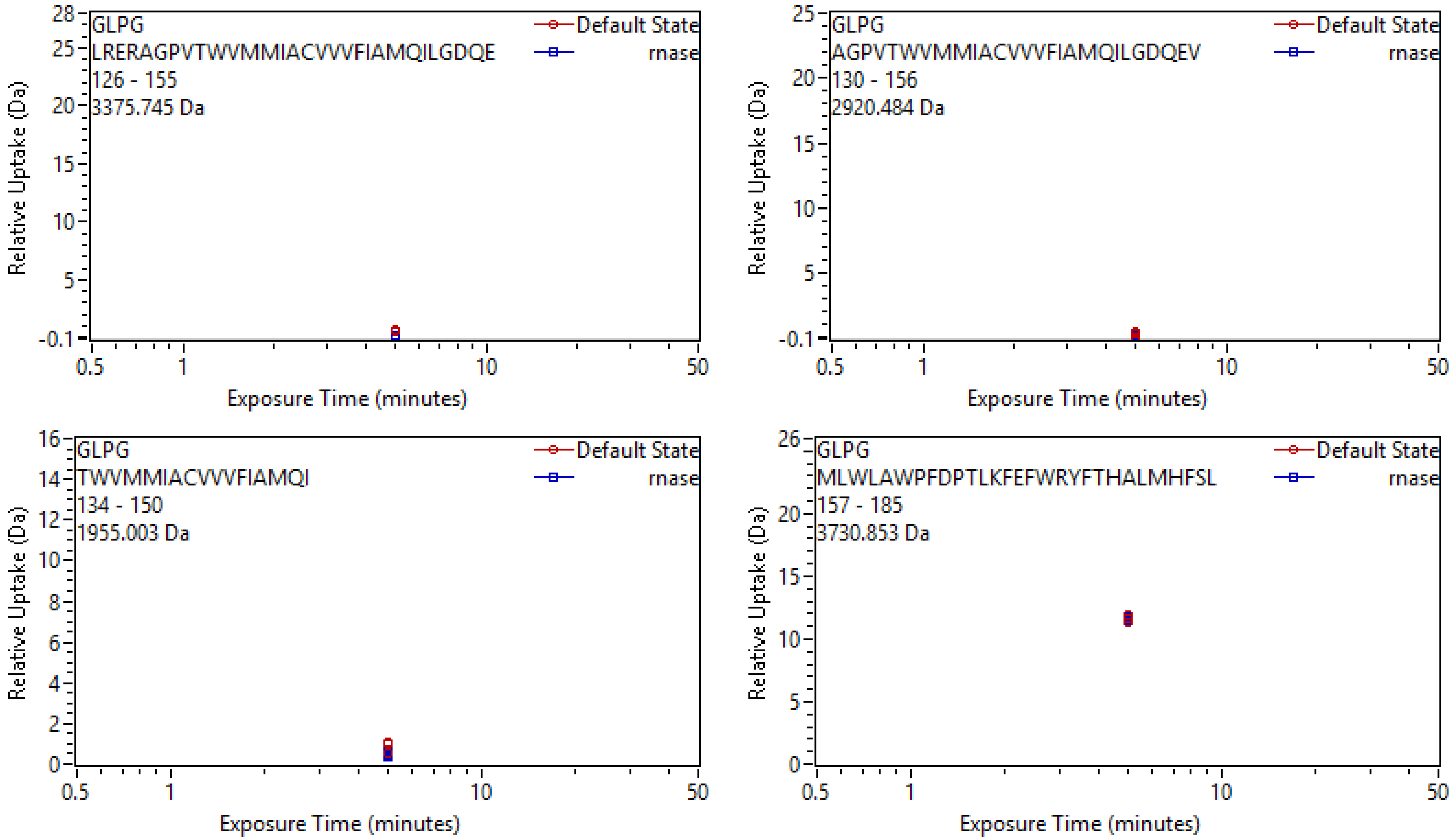

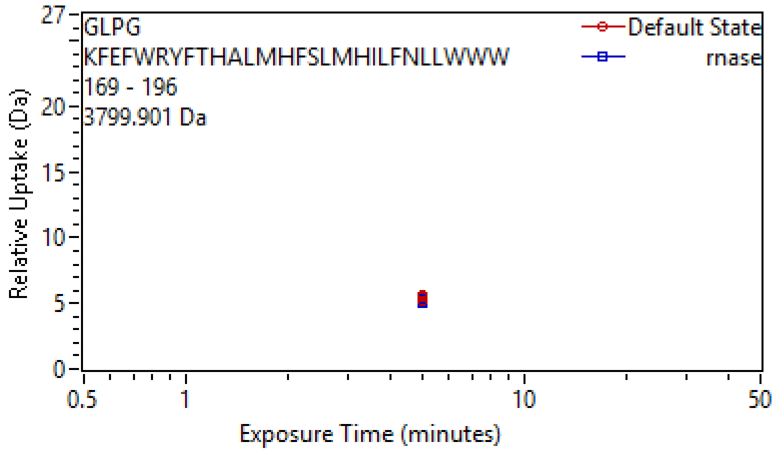

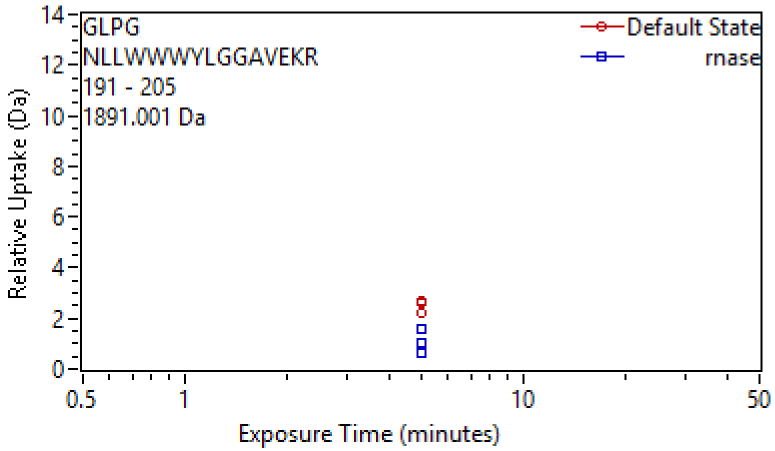

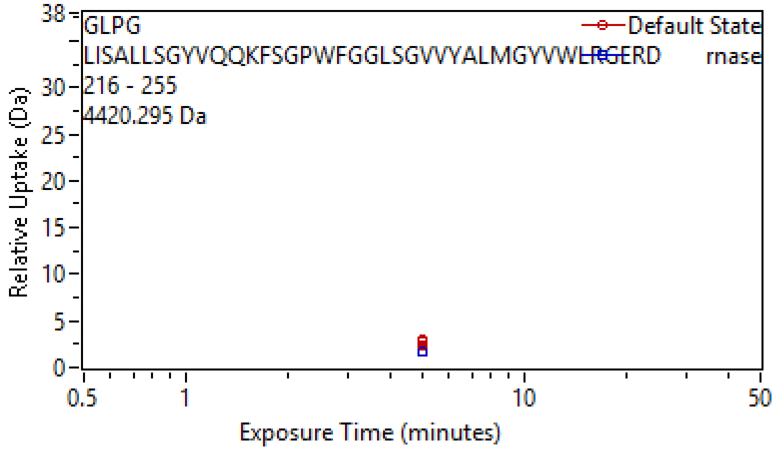

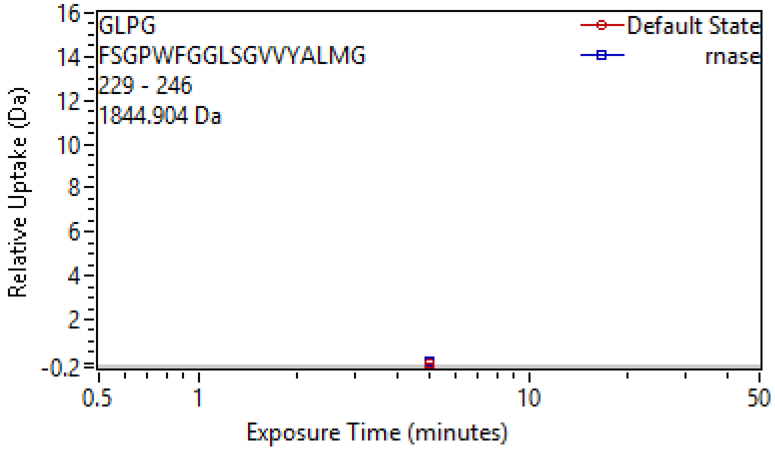

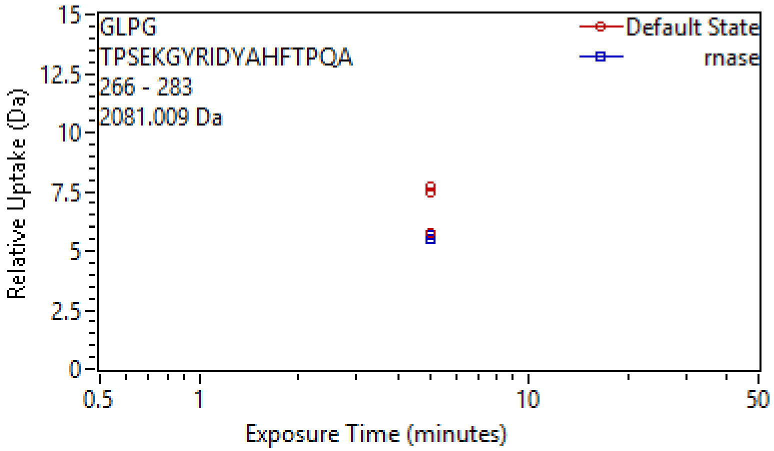
