## Supplementary figures and images for "LemonCatcher Acidic Pull-Down Enables Selective In-Cell Hydrogen-Deuterium Exchange Mass Spectrometry"

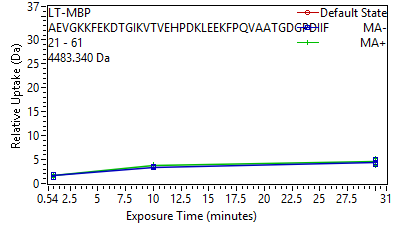

Supplement: Supporting Data File 2 [file 747387_file04.zip › uptake plots_cond 2_022026/006 LT-MBP[60-100] AEVGKKFEKDTGIKVT.png]

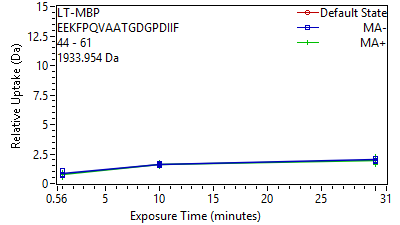

Supplement: Supporting Data File 2 [file 747387_file04.zip › uptake plots_cond 2_022026/007 LT-MBP[83-100] EEKFPQVAATGDGPDI.png]

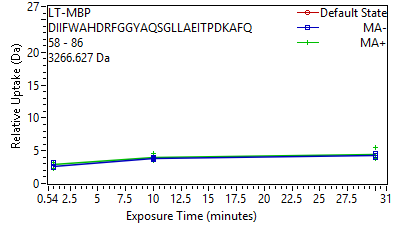

Supplement: Supporting Data File 2 [file 747387_file04.zip › uptake plots_cond 2_022026/008 LT-MBP[97-125] DIIFWAHDRFGGYAQS.png]

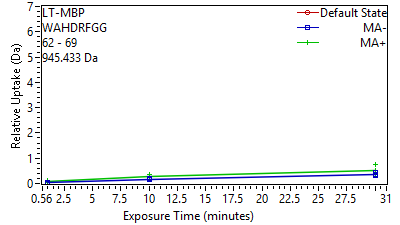

Supplement: Supporting Data File 2 [file 747387_file04.zip › uptake plots_cond 2_022026/009 LT-MBP[101-108] WAHDRFGG.png]

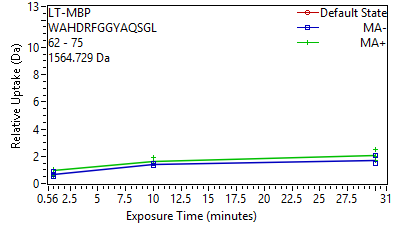

Supplement: Supporting Data File 2 [file 747387_file04.zip › uptake plots_cond 2_022026/010 LT-MBP[101-114] WAHDRFGGYAQSGL.png]

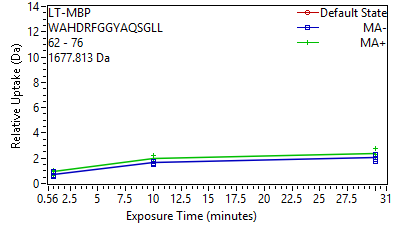

Supplement: Supporting Data File 2 [file 747387_file04.zip › uptake plots_cond 2_022026/011 LT-MBP[101-115] WAHDRFGGYAQSGLL.png]

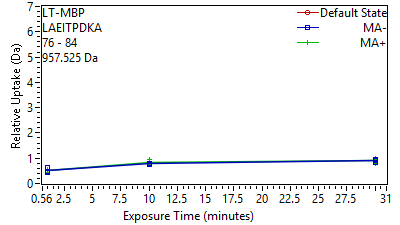

Supplement: Supporting Data File 2 [file 747387_file04.zip › uptake plots_cond 2_022026/012 LT-MBP[115-123] LAEITPDKA.png]

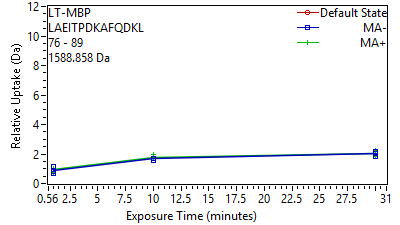

Supplement: Supporting Data File 2 [file 747387_file04.zip › uptake plots_cond 2_022026/013 LT-MBP[115-128] LAEITPDKAFQDKL.png]

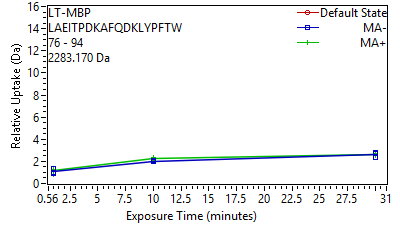

Supplement: Supporting Data File 2 [file 747387_file04.zip › uptake plots_cond 2_022026/014 LT-MBP[115-133] LAEITPDKAFQDKLYP.png]

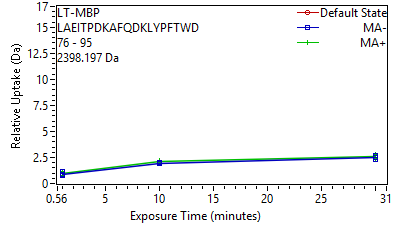

Supplement: Supporting Data File 2 [file 747387_file04.zip › uptake plots_cond 2_022026/015 LT-MBP[115-134] LAEITPDKAFQDKLYP.png]

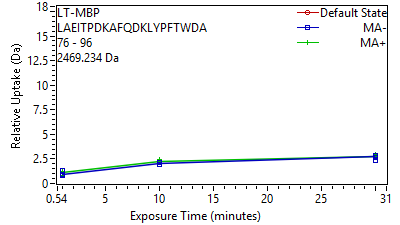

Supplement: Supporting Data File 2 [file 747387_file04.zip › uptake plots_cond 2_022026/016 LT-MBP[115-135] LAEITPDKAFQDKLYP.png]

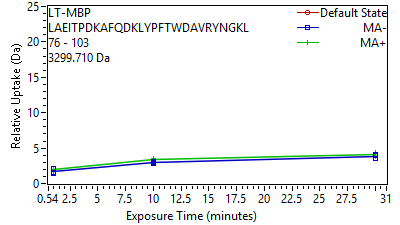

Supplement: Supporting Data File 2 [file 747387_file04.zip › uptake plots_cond 2_022026/017 LT-MBP[115-142] LAEITPDKAFQDKLYP.png]

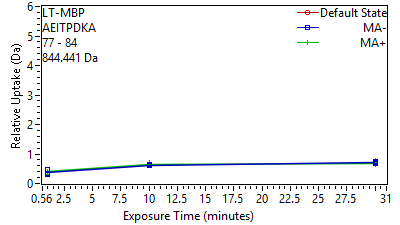

Supplement: Supporting Data File 2 [file 747387_file04.zip › uptake plots_cond 2_022026/018 LT-MBP[116-123] AEITPDKA.png]

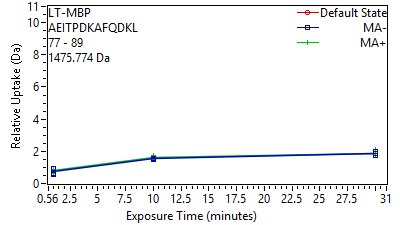

Supplement: Supporting Data File 2 [file 747387_file04.zip › uptake plots_cond 2_022026/019 LT-MBP[116-128] AEITPDKAFQDKL.png]

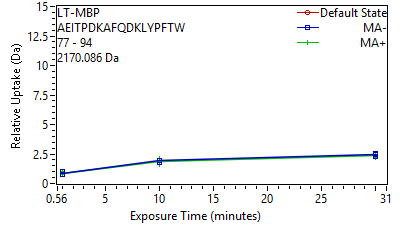

Supplement: Supporting Data File 2 [file 747387_file04.zip › uptake plots_cond 2_022026/020 LT-MBP[116-133] AEITPDKAFQDKLYPF.png]

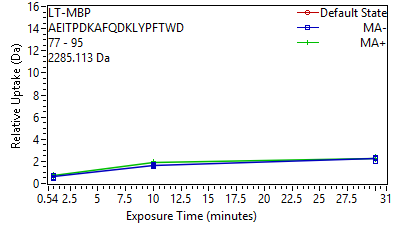

Supplement: Supporting Data File 2 [file 747387_file04.zip › uptake plots_cond 2_022026/021 LT-MBP[116-134] AEITPDKAFQDKLYPF.png]

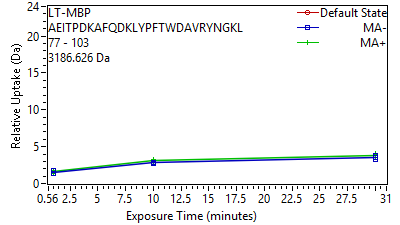

Supplement: Supporting Data File 2 [file 747387_file04.zip › uptake plots_cond 2_022026/022 LT-MBP[116-142] AEITPDKAFQDKLYPF.png]

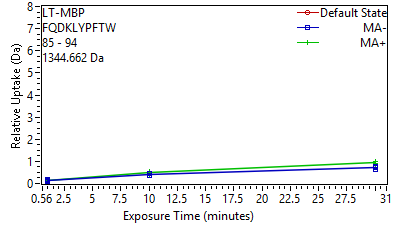

Supplement: Supporting Data File 2 [file 747387_file04.zip › uptake plots_cond 2_022026/023 LT-MBP[124-133] FQDKLYPFTW.png]

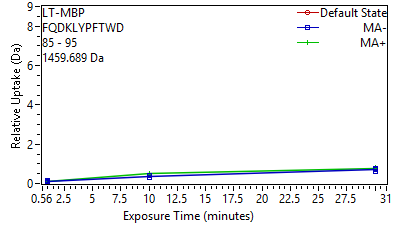

Supplement: Supporting Data File 2 [file 747387_file04.zip › uptake plots_cond 2_022026/024 LT-MBP[124-134] FQDKLYPFTWD.png]

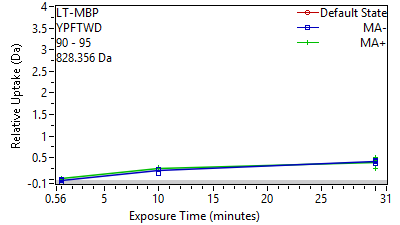

Supplement: Supporting Data File 2 [file 747387_file04.zip › uptake plots_cond 2_022026/025 LT-MBP[129-134] YPFTWD.png]

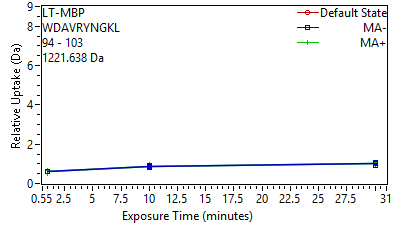

Supplement: Supporting Data File 2 [file 747387_file04.zip › uptake plots_cond 2_022026/026 LT-MBP[133-142] WDAVRYNGKL.png]

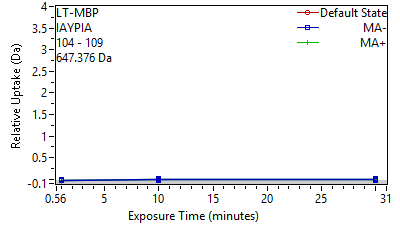

Supplement: Supporting Data File 2 [file 747387_file04.zip › uptake plots_cond 2_022026/027 LT-MBP[143-148] IAYPIA.png]

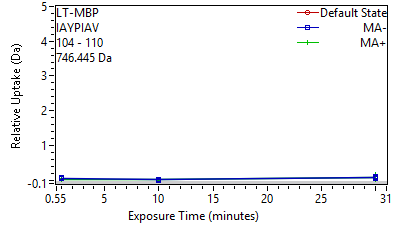

Supplement: Supporting Data File 2 [file 747387_file04.zip › uptake plots_cond 2_022026/028 LT-MBP[143-149] IAYPIAV.png]

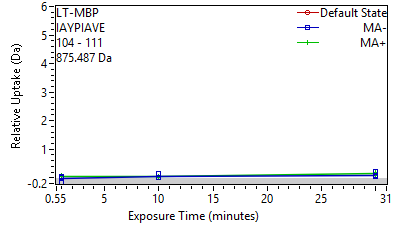

Supplement: Supporting Data File 2 [file 747387_file04.zip › uptake plots_cond 2_022026/029 LT-MBP[143-150] IAYPIAVE.png]

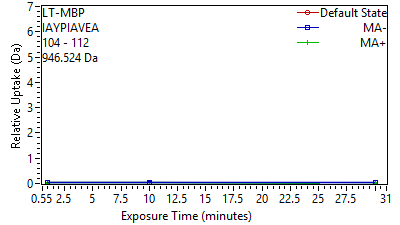

Supplement: Supporting Data File 2 [file 747387_file04.zip › uptake plots_cond 2_022026/030 LT-MBP[143-151] IAYPIAVEA.png]

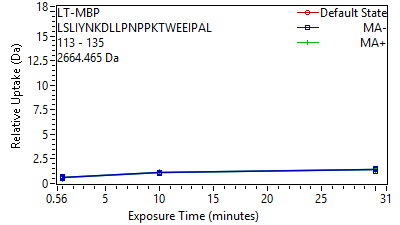

Supplement: Supporting Data File 2 [file 747387_file04.zip › uptake plots_cond 2_022026/031 LT-MBP[152-174] LSLIYNKDLLPNPPKT.png]

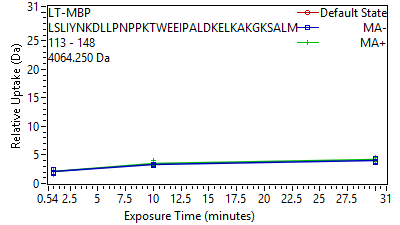

Supplement: Supporting Data File 2 [file 747387_file04.zip › uptake plots_cond 2_022026/032 LT-MBP[152-187] LSLIYNKDLLPNPPKT.png]

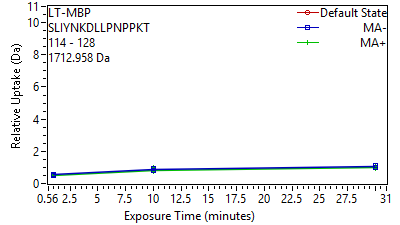

Supplement: Supporting Data File 2 [file 747387_file04.zip › uptake plots_cond 2_022026/033 LT-MBP[153-167] SLIYNKDLLPNPPKT.png]

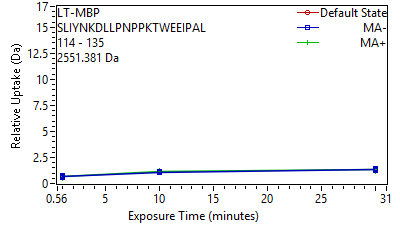

Supplement: Supporting Data File 2 [file 747387_file04.zip › uptake plots_cond 2_022026/034 LT-MBP[153-174] SLIYNKDLLPNPPKTW.png]

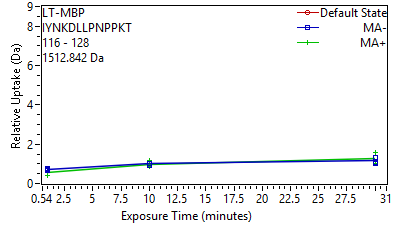

Supplement: Supporting Data File 2 [file 747387_file04.zip › uptake plots_cond 2_022026/035 LT-MBP[155-167] IYNKDLLPNPPKT.png]

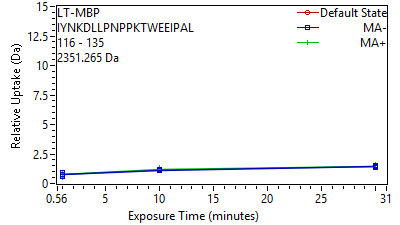

Supplement: Supporting Data File 2 [file 747387_file04.zip › uptake plots_cond 2_022026/036 LT-MBP[155-174] IYNKDLLPNPPKTWEE.png]

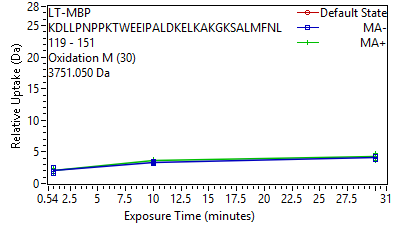

Supplement: Supporting Data File 2 [file 747387_file04.zip › uptake plots_cond 2_022026/037 LT-MBP[158-190] KDLLPNPPKTWEEIPA.png]

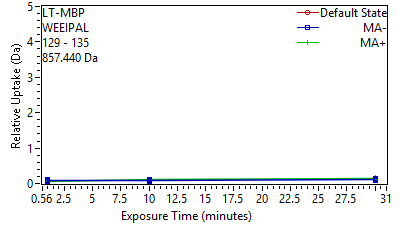

Supplement: Supporting Data File 2 [file 747387_file04.zip › uptake plots_cond 2_022026/038 LT-MBP[168-174] WEEIPAL.png]

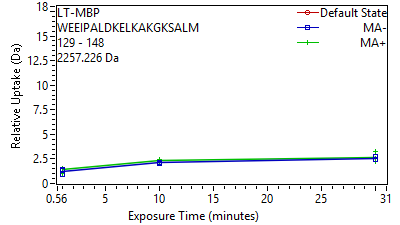

Supplement: Supporting Data File 2 [file 747387_file04.zip › uptake plots_cond 2_022026/039 LT-MBP[168-187] WEEIPALDKELKAKGK.png]

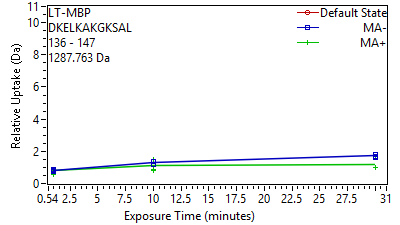

Supplement: Supporting Data File 2 [file 747387_file04.zip › uptake plots_cond 2_022026/040 LT-MBP[175-186] DKELKAKGKSAL.png]

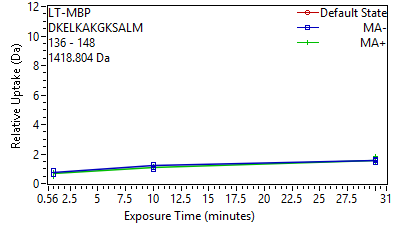

Supplement: Supporting Data File 2 [file 747387_file04.zip › uptake plots_cond 2_022026/041 LT-MBP[175-187] DKELKAKGKSALM.png]

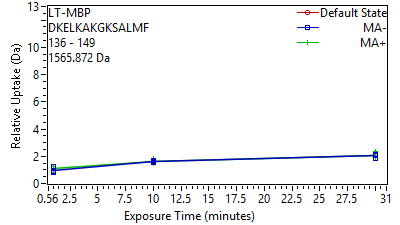

Supplement: Supporting Data File 2 [file 747387_file04.zip › uptake plots_cond 2_022026/042 LT-MBP[175-188] DKELKAKGKSALMF.png]

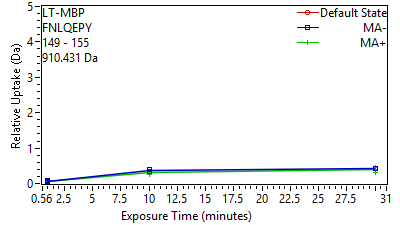

Supplement: Supporting Data File 2 [file 747387_file04.zip › uptake plots_cond 2_022026/043 LT-MBP[188-194] FNLQEPY.png]

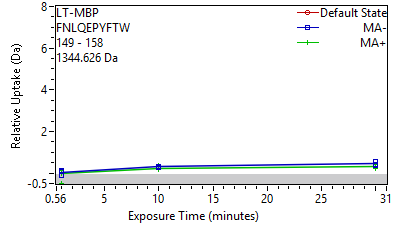

Supplement: Supporting Data File 2 [file 747387_file04.zip › uptake plots_cond 2_022026/044 LT-MBP[188-197] FNLQEPYFTW.png]

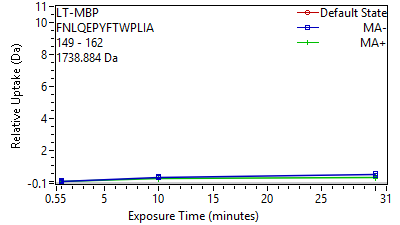

Supplement: Supporting Data File 2 [file 747387_file04.zip › uptake plots_cond 2_022026/045 LT-MBP[188-201] FNLQEPYFTWPLIA.png]

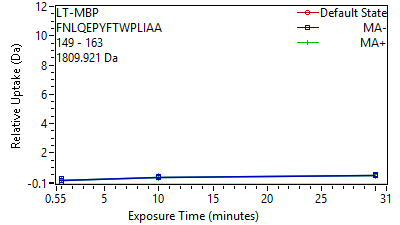

Supplement: Supporting Data File 2 [file 747387_file04.zip › uptake plots_cond 2_022026/046 LT-MBP[188-202] FNLQEPYFTWPLIAA.png]

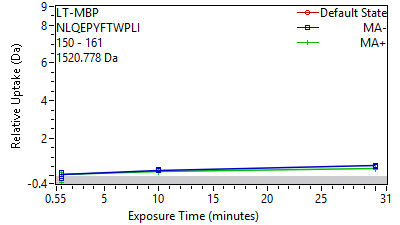

Supplement: Supporting Data File 2 [file 747387_file04.zip › uptake plots_cond 2_022026/047 LT-MBP[189-200] NLQEPYFTWPLI.png]

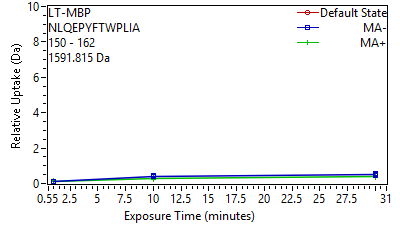

Supplement: Supporting Data File 2 [file 747387_file04.zip › uptake plots_cond 2_022026/048 LT-MBP[189-201] NLQEPYFTWPLIA.png]

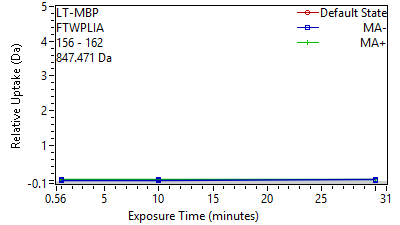

Supplement: Supporting Data File 2 [file 747387_file04.zip › uptake plots_cond 2_022026/049 LT-MBP[195-201] FTWPLIA.png]

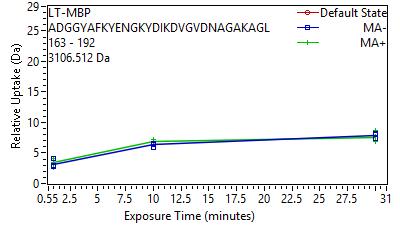

Supplement: Supporting Data File 2 [file 747387_file04.zip › uptake plots_cond 2_022026/050 LT-MBP[202-231] ADGGYAFKYENGKYDI.png]

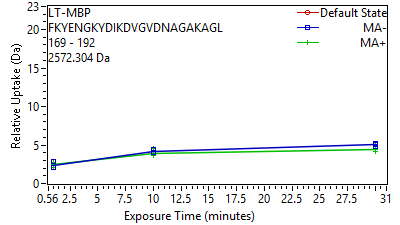

Supplement: Supporting Data File 2 [file 747387_file04.zip › uptake plots_cond 2_022026/051 LT-MBP[208-231] FKYENGKYDIKDVGVD.png]

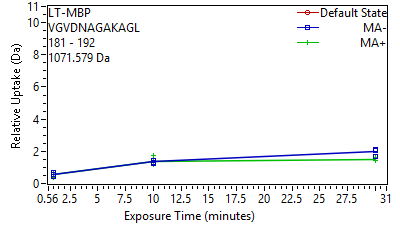

Supplement: Supporting Data File 2 [file 747387_file04.zip › uptake plots_cond 2_022026/052 LT-MBP[220-231] VGVDNAGAKAGL.png]

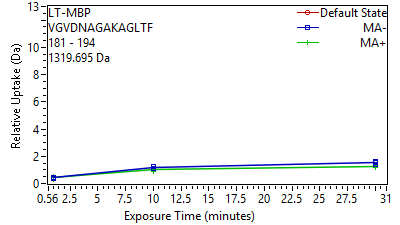

Supplement: Supporting Data File 2 [file 747387_file04.zip › uptake plots_cond 2_022026/053 LT-MBP[220-233] VGVDNAGAKAGLTF.png]

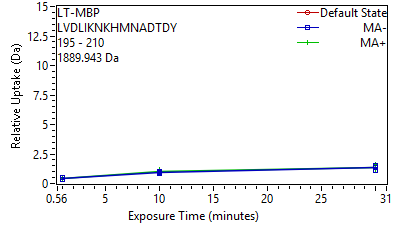

Supplement: Supporting Data File 2 [file 747387_file04.zip › uptake plots_cond 2_022026/054 LT-MBP[234-249] LVDLIKNKHMNADTDY.png]

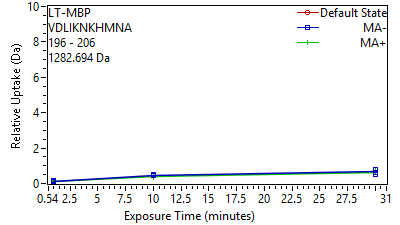

Supplement: Supporting Data File 2 [file 747387_file04.zip › uptake plots_cond 2_022026/055 LT-MBP[235-245] VDLIKNKHMNA.png]

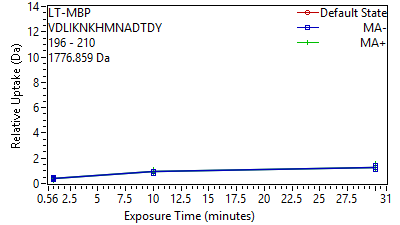

Supplement: Supporting Data File 2 [file 747387_file04.zip › uptake plots_cond 2_022026/056 LT-MBP[235-249] VDLIKNKHMNADTDY.png]

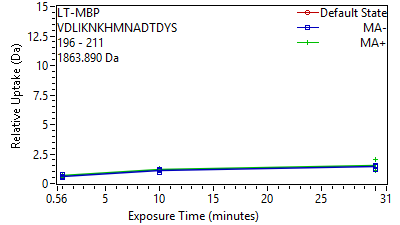

Supplement: Supporting Data File 2 [file 747387_file04.zip › uptake plots_cond 2_022026/057 LT-MBP[235-250] VDLIKNKHMNADTDYS.png]

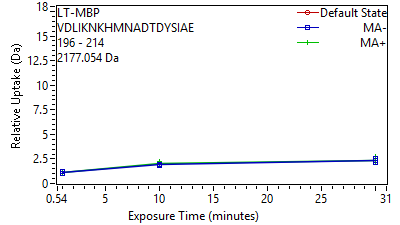

Supplement: Supporting Data File 2 [file 747387_file04.zip › uptake plots_cond 2_022026/058 LT-MBP[235-253] VDLIKNKHMNADTDYS.png]

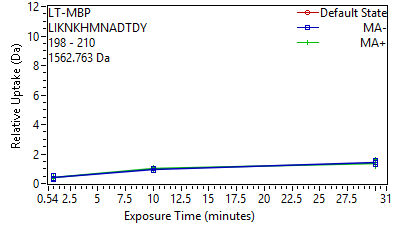

Supplement: Supporting Data File 2 [file 747387_file04.zip › uptake plots_cond 2_022026/059 LT-MBP[237-249] LIKNKHMNADTDY.png]

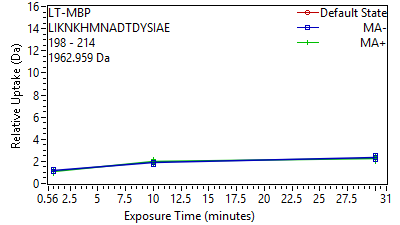

Supplement: Supporting Data File 2 [file 747387_file04.zip › uptake plots_cond 2_022026/060 LT-MBP[237-253] LIKNKHMNADTDYSIA.png]

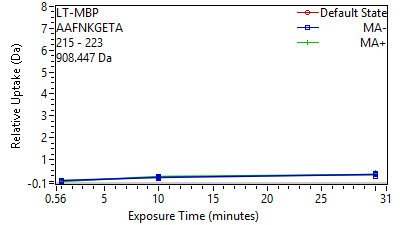

Supplement: Supporting Data File 2 [file 747387_file04.zip › uptake plots_cond 2_022026/061 LT-MBP[254-262] AAFNKGETA.png]

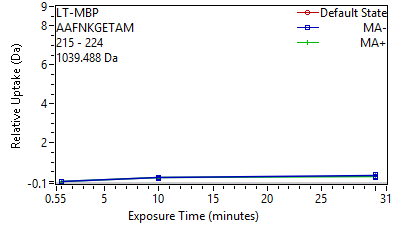

Supplement: Supporting Data File 2 [file 747387_file04.zip › uptake plots_cond 2_022026/062 LT-MBP[254-263] AAFNKGETAM.png]

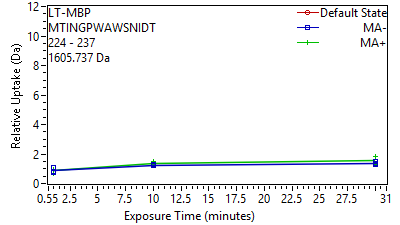

Supplement: Supporting Data File 2 [file 747387_file04.zip › uptake plots_cond 2_022026/063 LT-MBP[263-276] MTINGPWAWSNIDT.png]

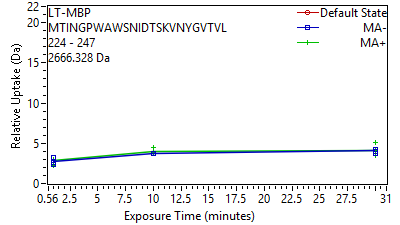

Supplement: Supporting Data File 2 [file 747387_file04.zip › uptake plots_cond 2_022026/064 LT-MBP[263-286] MTINGPWAWSNIDTSK.png]

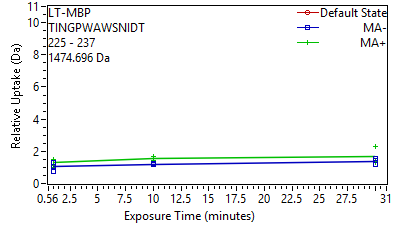

Supplement: Supporting Data File 2 [file 747387_file04.zip › uptake plots_cond 2_022026/065 LT-MBP[264-276] TINGPWAWSNIDT.png]

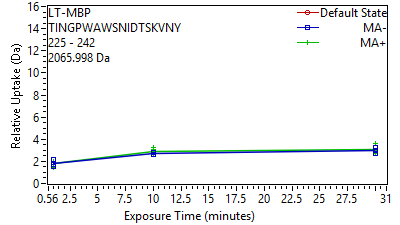

Supplement: Supporting Data File 2 [file 747387_file04.zip › uptake plots_cond 2_022026/066 LT-MBP[264-281] TINGPWAWSNIDTSKV.png]

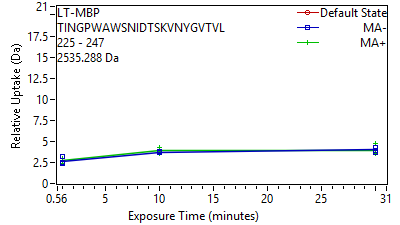

Supplement: Supporting Data File 2 [file 747387_file04.zip › uptake plots_cond 2_022026/067 LT-MBP[264-286] TINGPWAWSNIDTSKV.png]

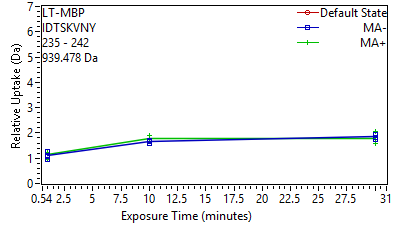

Supplement: Supporting Data File 2 [file 747387_file04.zip › uptake plots_cond 2_022026/068 LT-MBP[274-281] IDTSKVNY.png]

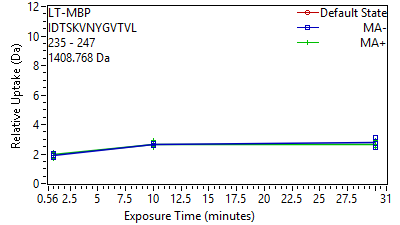

Supplement: Supporting Data File 2 [file 747387_file04.zip › uptake plots_cond 2_022026/069 LT-MBP[274-286] IDTSKVNYGVTVL.png]

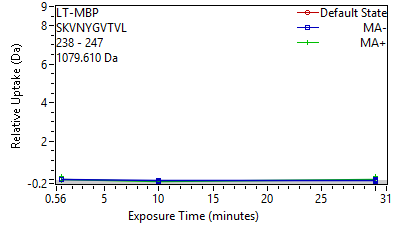

Supplement: Supporting Data File 2 [file 747387_file04.zip › uptake plots_cond 2_022026/070 LT-MBP[277-286] SKVNYGVTVL.png]

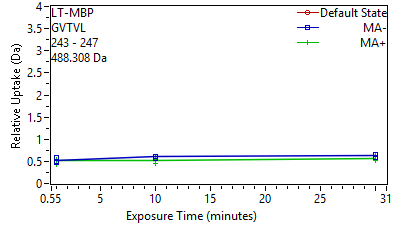

Supplement: Supporting Data File 2 [file 747387_file04.zip › uptake plots_cond 2_022026/071 LT-MBP[282-286] GVTVL.png]

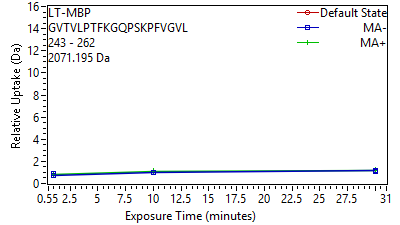

Supplement: Supporting Data File 2 [file 747387_file04.zip › uptake plots_cond 2_022026/072 LT-MBP[282-301] GVTVLPTFKGQPSKPF.png]

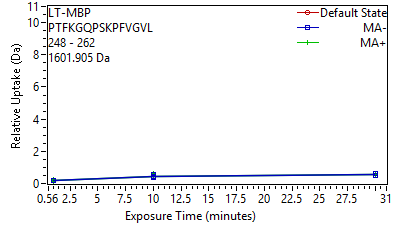

Supplement: Supporting Data File 2 [file 747387_file04.zip › uptake plots_cond 2_022026/073 LT-MBP[287-301] PTFKGQPSKPFVGVL.png]

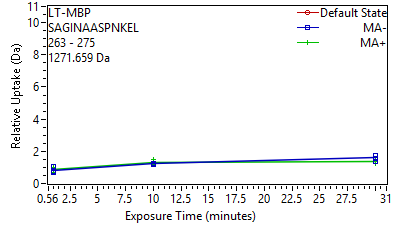

Supplement: Supporting Data File 2 [file 747387_file04.zip › uptake plots_cond 2_022026/074 LT-MBP[302-314] SAGINAASPNKEL.png]

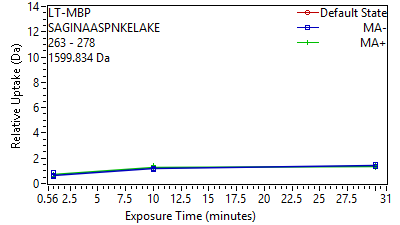

Supplement: Supporting Data File 2 [file 747387_file04.zip › uptake plots_cond 2_022026/075 LT-MBP[302-317] SAGINAASPNKELAKE.png]

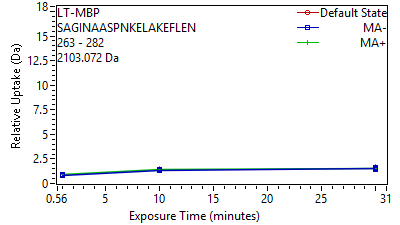

Supplement: Supporting Data File 2 [file 747387_file04.zip › uptake plots_cond 2_022026/076 LT-MBP[302-321] SAGINAASPNKELAKE.png]

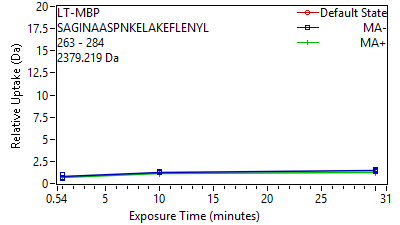

Supplement: Supporting Data File 2 [file 747387_file04.zip › uptake plots_cond 2_022026/077 LT-MBP[302-323] SAGINAASPNKELAKE.png]

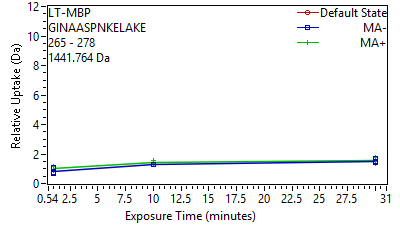

Supplement: Supporting Data File 2 [file 747387_file04.zip › uptake plots_cond 2_022026/078 LT-MBP[304-317] GINAASPNKELAKE.png]

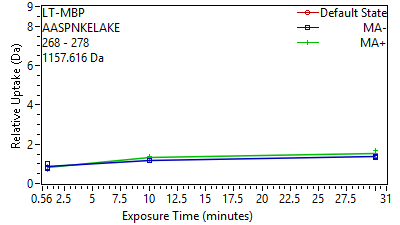

Supplement: Supporting Data File 2 [file 747387_file04.zip › uptake plots_cond 2_022026/079 LT-MBP[307-317] AASPNKELAKE.png]

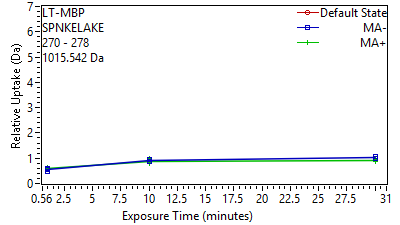

Supplement: Supporting Data File 2 [file 747387_file04.zip › uptake plots_cond 2_022026/080 LT-MBP[309-317] SPNKELAKE.png]

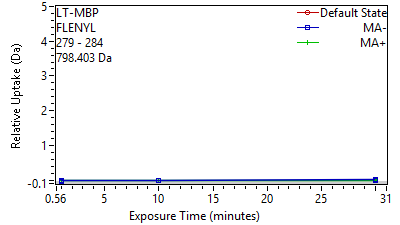

Supplement: Supporting Data File 2 [file 747387_file04.zip › uptake plots_cond 2_022026/081 LT-MBP[318-323] FLENYL.png]

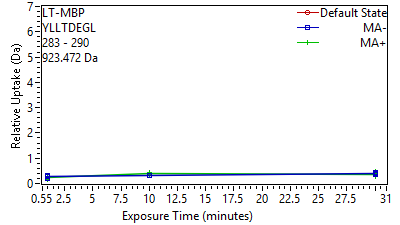

Supplement: Supporting Data File 2 [file 747387_file04.zip › uptake plots_cond 2_022026/082 LT-MBP[322-329] YLLTDEGL.png]

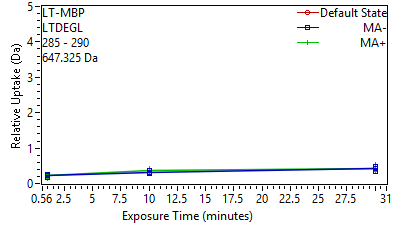

Supplement: Supporting Data File 2 [file 747387_file04.zip › uptake plots_cond 2_022026/083 LT-MBP[324-329] LTDEGL.png]

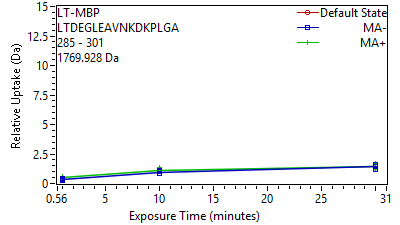

Supplement: Supporting Data File 2 [file 747387_file04.zip › uptake plots_cond 2_022026/084 LT-MBP[324-340] LTDEGLEAVNKDKPLG.png]

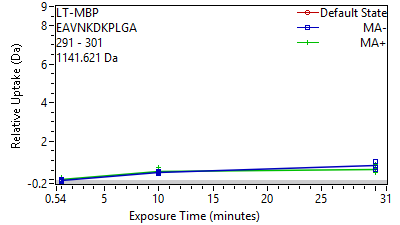

Supplement: Supporting Data File 2 [file 747387_file04.zip › uptake plots_cond 2_022026/085 LT-MBP[330-340] EAVNKDKPLGA.png]

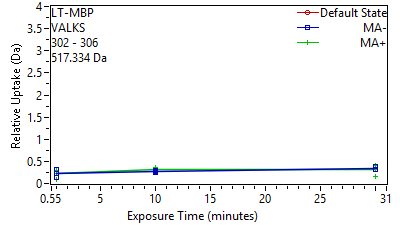

Supplement: Supporting Data File 2 [file 747387_file04.zip › uptake plots_cond 2_022026/086 LT-MBP[341-345] VALKS.png]

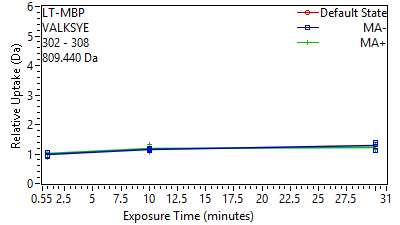

Supplement: Supporting Data File 2 [file 747387_file04.zip › uptake plots_cond 2_022026/087 LT-MBP[341-347] VALKSYE.png]

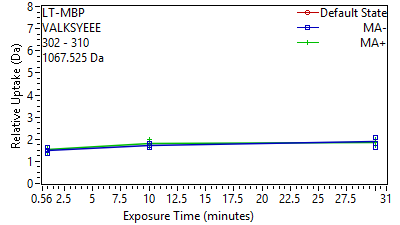

Supplement: Supporting Data File 2 [file 747387_file04.zip › uptake plots_cond 2_022026/088 LT-MBP[341-349] VALKSYEEE.png]

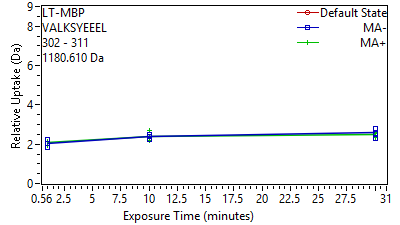

Supplement: Supporting Data File 2 [file 747387_file04.zip › uptake plots_cond 2_022026/089 LT-MBP[341-350] VALKSYEEEL.png]

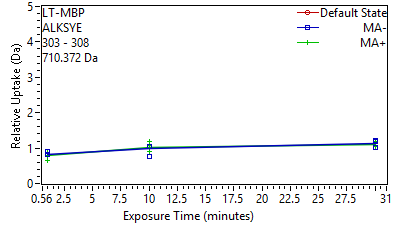

Supplement: Supporting Data File 2 [file 747387_file04.zip › uptake plots_cond 2_022026/090 LT-MBP[342-347] ALKSYE.png]

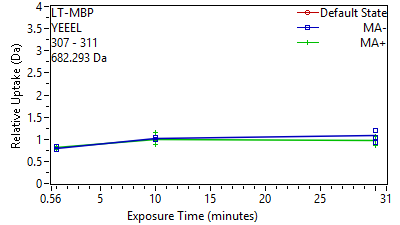

Supplement: Supporting Data File 2 [file 747387_file04.zip › uptake plots_cond 2_022026/091 LT-MBP[346-350] YEEEL.png]

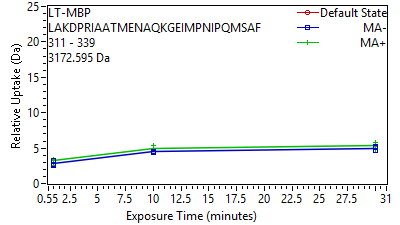

Supplement: Supporting Data File 2 [file 747387_file04.zip › uptake plots_cond 2_022026/092 LT-MBP[350-378] LAKDPRIAATMENAQK.png]

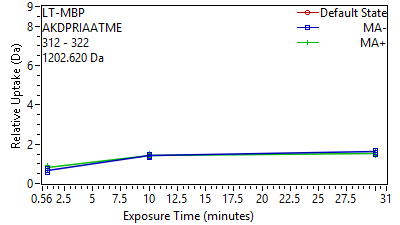

Supplement: Supporting Data File 2 [file 747387_file04.zip › uptake plots_cond 2_022026/093 LT-MBP[351-361] AKDPRIAATME.png]

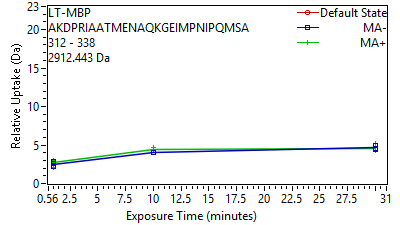

Supplement: Supporting Data File 2 [file 747387_file04.zip › uptake plots_cond 2_022026/094 LT-MBP[351-377] AKDPRIAATMENAQKG.png]

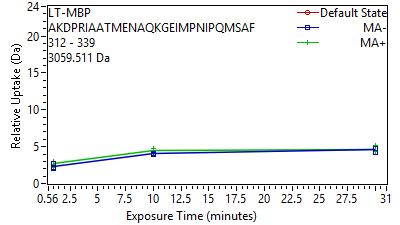

Supplement: Supporting Data File 2 [file 747387_file04.zip › uptake plots_cond 2_022026/095 LT-MBP[351-378] AKDPRIAATMENAQKG.png]

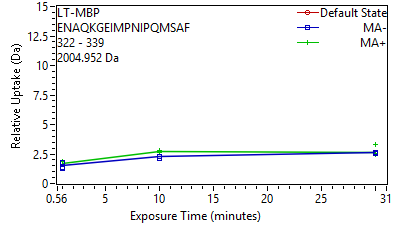

Supplement: Supporting Data File 2 [file 747387_file04.zip › uptake plots_cond 2_022026/096 LT-MBP[361-378] ENAQKGEIMPNIPQMS.png]

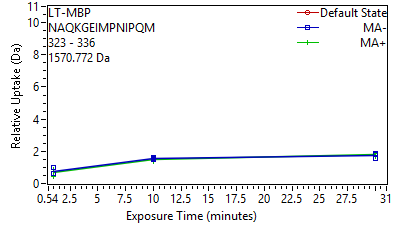

Supplement: Supporting Data File 2 [file 747387_file04.zip › uptake plots_cond 2_022026/097 LT-MBP[362-375] NAQKGEIMPNIPQM.png]

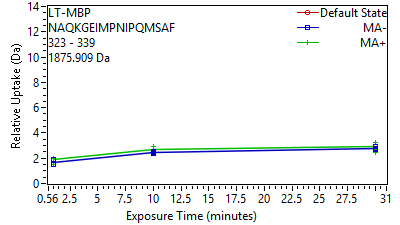

Supplement: Supporting Data File 2 [file 747387_file04.zip › uptake plots_cond 2_022026/098 LT-MBP[362-378] NAQKGEIMPNIPQMSA.png]

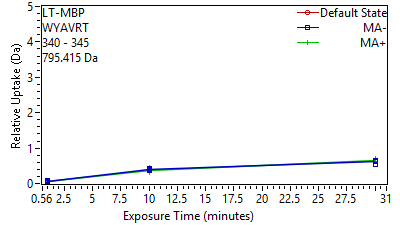

Supplement: Supporting Data File 2 [file 747387_file04.zip › uptake plots_cond 2_022026/099 LT-MBP[379-384] WYAVRT.png]

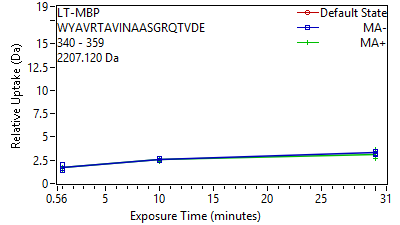

Supplement: Supporting Data File 2 [file 747387_file04.zip › uptake plots_cond 2_022026/100 LT-MBP[379-398] WYAVRTAVINAASGRQ.png]

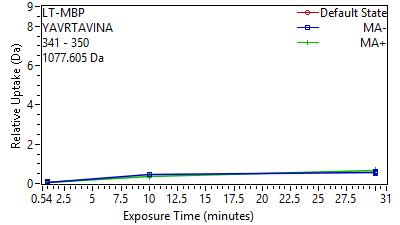

Supplement: Supporting Data File 2 [file 747387_file04.zip › uptake plots_cond 2_022026/101 LT-MBP[380-389] YAVRTAVINA.png]

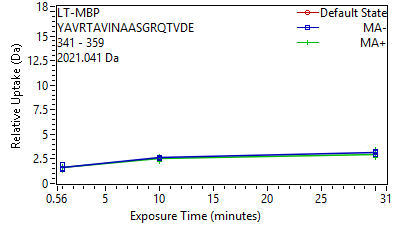

Supplement: Supporting Data File 2 [file 747387_file04.zip › uptake plots_cond 2_022026/102 LT-MBP[380-398] YAVRTAVINAASGRQT.png]

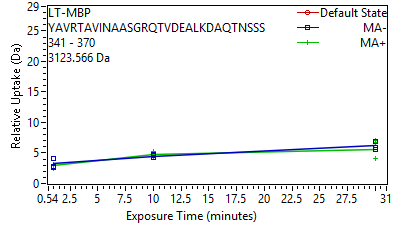

Supplement: Supporting Data File 2 [file 747387_file04.zip › uptake plots_cond 2_022026/103 LT-MBP[380-409] YAVRTAVINAASGRQT.png]

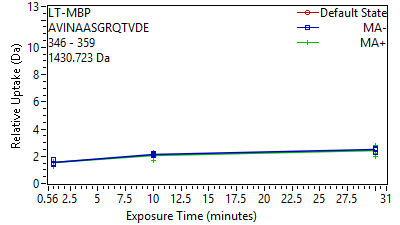

Supplement: Supporting Data File 2 [file 747387_file04.zip › uptake plots_cond 2_022026/104 LT-MBP[385-398] AVINAASGRQTVDE.png]

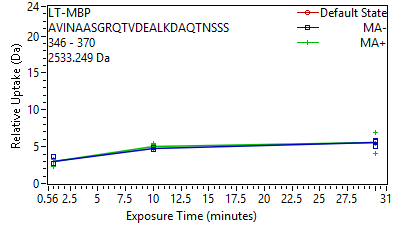

Supplement: Supporting Data File 2 [file 747387_file04.zip › uptake plots_cond 2_022026/105 LT-MBP[385-409] AVINAASGRQTVDEAL.png]
